# A Tunable Pulmonary Organoid Model Demonstrates Compositionally Driven Epithelial Plasticity and Immune Polarization

**DOI:** 10.1101/2025.06.05.658120

**Authors:** Sophie E. Edelstein, Satoshi Mizoguchi, Maria Tomàs Gracia, Nuoya Wang, Vi Lee, Hahram Kim, Connor Haynes, Colten Danelski, Tomoshi Tsuchiya, Maor Sauler, Micha Sam Brickman Raredon

**Affiliations:** Department of Anesthesiology, Yale School of Medicine, New Haven, CT 06520, USA; Vascular Biology and Therapeutics Program, Yale School of Medicine, New Haven, CT 06520, USA; Medical Student Training Program, Yale School of Medicine, New Haven, CT 06520, USA; Department of Biomedical Engineering, Yale University, New Haven, CT 06511, USA; Department of Thoracic Surgery, University of Toyama, Toyama, 9300194, Japan; Department of Pulmonary, Critical Care and Sleep Medicine, Yale School of Medicine, New Haven, CT 06520, USA; Center for Pulmonary Injury, Inflammation, Repair and Therapeutics (CPIRT), Yale School of Medicine, New Haven, CT 06520, USA; Program in Translational Biomedicine (PTB), Yale School of Medicine, New Haven, CT 06520, USA

## Abstract

Aberrant epithelial regeneration and immune remodeling are hallmarks of chronic lung diseases such as idiopathic pulmonary fibrosis (IPF), COPD, and post-viral syndromes. Yet how cellular context shapes these trajectories remains unresolved. We present a tunable, primary rat-derived lung organoid model that systematically varies immune, epithelial, and mesenchymal inputs to reveal how composition alone dictates epithelial plasticity and macrophage polarization. Across organoid conditions that varied by relative starting lineage ratios, we observed the spontaneous emergence of disease-relevant transitional cell states, including *Sox9^+^* stressed progenitors, RAS-like intermediates, and hillock-like cells, alongside distinct macrophage activation profiles. In mesenchyme-rich contexts, epithelial-immune-mesenchymal crosstalk appeared to reinforce inflammatory signaling and stabilize transitional persistence, while immune-dominant inputs favored ATI-like repair and squamous remodeling. Hillock-like cells displayed context-specific polarization and expressed immune-regulatory genes, suggesting a role as epithelial orchestrators that help calibrate inflammatory response during regeneration. Connectomic analysis via NICHES revealed that regenerative outcomes were associated with dynamic multicellular signaling networks that integrate stress sensing, immune coordination, and epithelial resilience. This platform provides a tractable system for modeling milieu-specific repair and regenerative mechanisms and could inform therapeutic strategies aimed at redirecting epithelial fate in chronic lung disease.

## Introduction

Lung homeostasis, injury repair, and disease progression are all governed by epithelial plasticity (<u>Beppu et al., 2023</u>). Regeneration of the lung is regulated by a finely tuned system of interactions between epithelial, immune, and mesenchymal cells. Based on cues from their microenvironment, epithelial cells transition through multiple phenotypic and transcriptomic states, giving rise to transitional populations that include, but are not limited to: hillock, activated respiratory airway secretory (aRAS), alveolar type 0 (AT0) and damage associated transient progenitor (DATP) cells in response to injury and during the repair process (<u>Lin et al., 2024; Kadur Lakshminarasimha Murthy et al., 2022; Basil et al., 2021; Tata & Rajagopal, 2017; Choi et al., 2020</u>). These transitional cell types have become of significant interest to the pulmonary biology community in recent years due to their flexible lineage potential, transient activation following injury, and capacity to mediate divergent outcomes ranging from effective regeneration to maladaptive remodeling (<u>Choi et al., 2020; Konkimalla et al., 2023</u>).

Traditional 2D epithelial culture systems fail to recapitulate native tissue architecture and lack immune and mesenchymal components, which are essential for regulating epithelial steady state and fate (<u>Barnes et al., 2023; Jones et al., 2023</u>). Conversely, while in vivo animal models provide a physiologically relevant context, they do not allow for systematic dissection of how specific cellular lineages and the specific ratios of these lineages shape epithelial behavior under controlled conditions (<u>Murrow et al., 2017</u>).

Mesenchymal populations are central regulators of epithelial and immune cell behavior, mediating a vast array of context-dependent behaviors (<u>Zepp et al., 2017; Ghonim et al., 2023</u>). Epithelial-mesenchymal signaling coordinates spatial pattering and cell fate decisions in the developing lung, primarily through canonical pathways such as WNT and BMP (<u>Goltsis et al., 2024; Xu et al., 2011; Frum et al., 2023</u>). Moreover, mesenchymal cells are also capable of modulating immune cell polarization and cytokine production, thus modifying the inflammatory milieu and calibrating repair pathways (<u>Morrison et al., 2017; Mo et al., 2022</u>). These regulatory axes come together during injury and involve epithelial and immune coordination through IL-33, IL-1B. and TNFα to orchestrate tissue remodeling (<u>de Kleer et al., 2016; Wang et al., 2024; Katsura et al., 2019</u>). That being said, the specific mechanisms by which mesenchymal presence or absence dictates epithelial-immune coordination in the rat lung are difficult to isolate in vivo.

Pulmonary organoids are promising models for studying epithelial plasticity and multicellular dynamics in defined systems (<u>Nikolic & Rawlins, 2017</u>); however, most depend on genetic manipulations, exogenous injury cues, or pre-programmed differentiation protocols to induce epithelial transitions (<u>Leko et al., 2023; Sachs et al., 2019; Abo et al., 2022</u>). Here, we present a lung organoid model that leverages primary, rat-derived populations and allows for the study of epithelial plasticity and immune cell behavior, specifically macrophage polarization, without relying on pre-programmed or exogenous injury signals.

To investigate the influence of the relative lineage ratios at seeding on epithelial plasticity under uniform culture conditions, we isolated two distinct populations from the adult rat lung: (1) a pulmonary dissociation (PD), generated by enzymatic digestion of whole lung tissue and containing a heterogenous mix of epithelial, mesenchymal, endothelial, and immune cells; and (2) a bronchoalveolar lavage isolate collected via perfusion of the lungs, through the trachea, with cold saline to collect non-adherent immune cells from the surfaces of the airways. These populations were combined in different ratios to generate three organoid conditions: 100% PD (PD_3D), 100% BAL (BAL_3D), and a 1:1 mix of BAL to PD (Mixed_3D).

The spontaneous formation of reparative-like conditions occurred in all cultures, consistent with previous work demonstrating that engineered tissues often remodel under stress when taken out of native physiological regulation (<u>Zhang et al., 2024; Liu et al., 2015; Patel et al., 2018</u>). Despite the same media, matrix, and environmental parameters being used for all cultures (embryonic progenitor media dosed with potent growth factors and a 5% CO_2_ environment [<u>Nichane et al., 2017</u>]), similar, as well as distinct epithelial and immune cell states reproducibly emerged in the three conditions (PD_3D, Mixed_3D, BAL_3D). Among these derivatives were epithelial sub-types associated with injury such as *Krt13^+^*hillock-like cells and *Scgb3a2^+^/Sftpc^+^/Scgb1a1^+^*activated RAS cells, along with macrophage polarization states that were condition-specific (<u>Basil et al., 2022; Lin et al., 2024</u>). We propose that the appearance of these cell states was not the consequence of external perturbations but rather reflect variation in initial cellular composition. Our system does not deliberately impose damage using chemical, mechanical, or physical stimuli such as bleomycin treatment or cigarette smoke treatment (<u>Suezawa et al., 2021; Khedoe et al., 2023</u>). Organoid cultures, however, inherently mimic some stress-inducing characteristics such as local hypoxia, limitations in nutrient diffusion, and lack of physiological feedback leading to the priming of cells toward a transitional or reparative state (<u>Nwokoye & Abilez, 2024; Corro et al., 2020</u>).

In the results reported herein, we present a model system that implies that the ratios of epithelium, immune, endothelium, and mesenchyme relative to one another at the start of culture dictate the trajectory of multicellular interactions in a stress-permissive milieu, laying a foundation for understanding how lung tissues transition between homeostatic, reparative, and pathological states. Furthermore, differences in the starting cell suspension composition allowed for the emergence of transitional epithelial states and polarized macrophage patterns. Specifically, these findings suggest that cellular ratios can direct divergent regenerative trajectories whereby immune enrichment favors certain reparative or inflammatory fate and mesenchymal presence facilitates epithelial state transitions. By enabling the systematic variation in discrete lineage inputs, our model provides an experimental framework to interrogate how compositional changes influences developmental and injury-associated programs—a principle with broad implications for regenerative medicine and therapeutic reprogramming for lung repair.

## Results

### A compositionally tunable lung organoid system enables controlled investigation of epithelial and immune state emergence

The generation of our model started with two biologically accessible inputs derived from the native rat lung, an organ whose developmental trajectory, anatomical architecture, and cellular composition closely resembles that of the human lung and thus serves as an experimentally informative system for studying epithelial dynamics in health and disease (<u>Warburton et al., 2000; O’Reilly & Thébaud, 2014; Burri, 1974; Nardell & Brody, 1982</u>). The two populations utilized are as follows: (1) a pulmonary dissociation (PD) obtained by enzymatic digestion of whole lung tissue resulting in a heterogenous mixture of epithelial, mesenchymal, endothelial, and immune cells; and (2) a bronchoalveolar lavage (BAL) isolate enriched for non-adherent immune cells such as macrophages, neutrophils, B cells, and T cells, as well as a small, rare population of miscellaneous epithelium (**Fig. 1A**).

**Figure 1.**
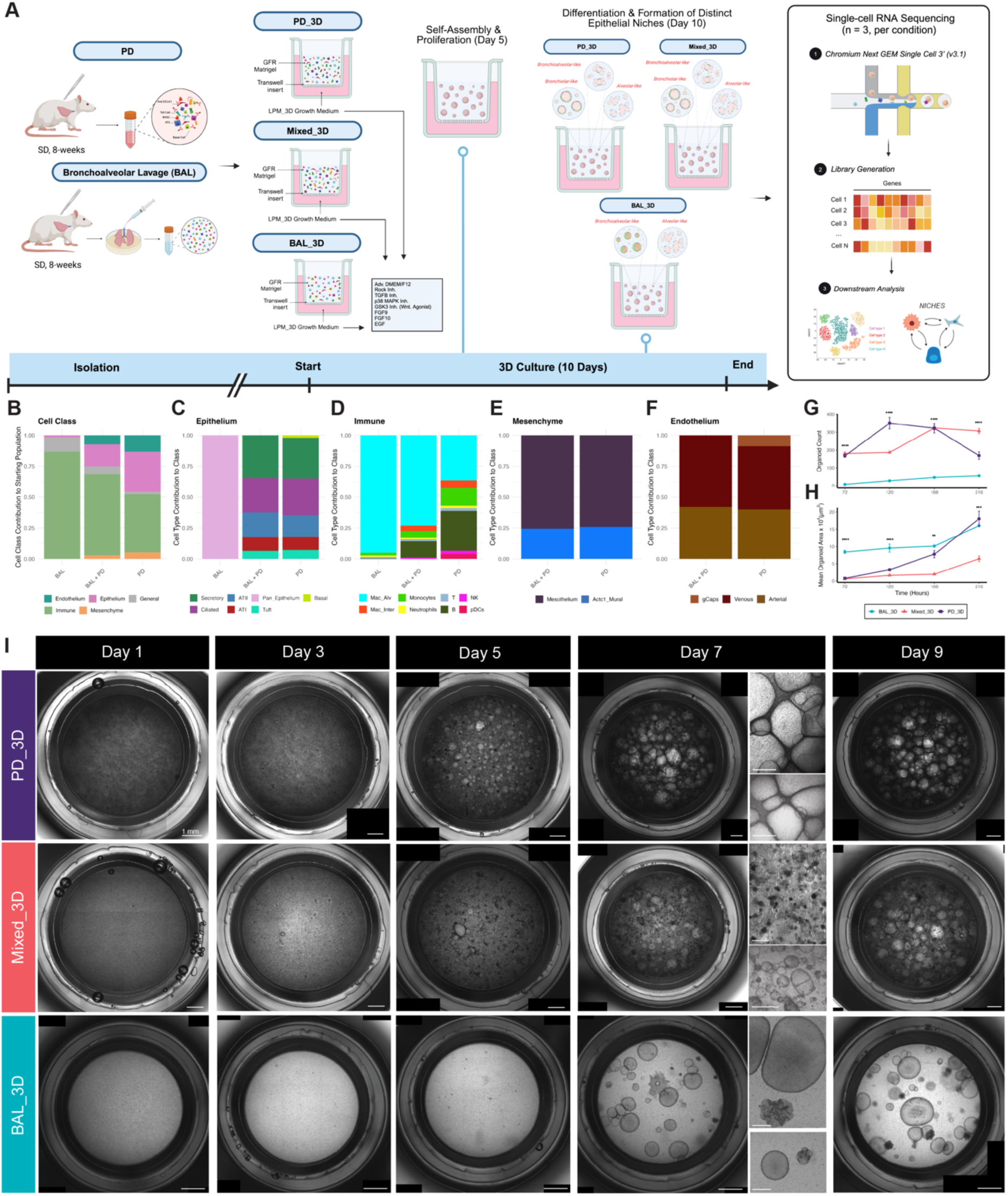
A tunable 3D lung organoid system enables controlled modeling of epithelial-immune-mesenchymal interactions. **A)** Schematic overview of experimental design. Cells were isolated from rat lungs via either pulmonary dissociation (PD) or bronchoalveolar lavage (BAL) at 8 weeks of age, then combined to generate three experimental groups: PD_3D (native lung dissociation), BAL_3D (immune-enriched), and Mixed_3D (1:1 mixture of PD and BAL cells). Organoids were maintained in LPM-3D progenitor medium through Day 1O. Organoids were harvested for single-cell RNA sequencing (10x Genomics Chromium, n *=* 3 replicates per condition) and processed for downstream transcriptomic and ligand-receptor signaling analysis (NICHES). **B-F)** Quantification of major cell class distributions across conditions using scRNA-seq data. Bar graphs show proportions of: **B)** major cell lineages. including a “General” category representing cycling cells that could not be definitively assigned to a single lineage; **C)** epithelial subsets, **D)** immune subsets, **E)** mesenchymal populations, and **F)** endothelial cells. **G-H)** Quantification of morphological metrics from 3D organoid cultures over time. Organoid number **(G)** and mean organoid area **(H)** were quantified using QuPath on stitched brighlfield images (n *=* 3 wells per condition per timepoint) (see *Methods)* , ***, p ≤ 0.001; *, p ≤ 0.01; p ≤ 0.05; ns, not significant; statistical comparisons performed using Welch’s I-tests with Benjamini-Hochberg correction for multiple comparisons. **I)** Brightfield images of 3D organoid cultures across time (Days 1, 3, 5, 7, 9). Representative wells are shown for each condition. Insets on Day 7 show zoomed views of organoid morphology. Scale bars, 1 mm (main panels) and 200 µm (insets).

scRNA-seq of these starting populations confirmed that PD (n = 3 - 9,730 cells) contained a diverse set of epithelial subsets, namely alveolar type 1 (*Pdpn⁺/Ager⁺)* (ATI), alveolar type 2 (*Sftpc⁺/Napsa⁺/Lamp3⁺)* (ATII), basal (*Krt5⁺/Tp63⁺)*, secretory (*Scgb3a1⁺/Scgb3a2⁺)*, ciliated (*Ccdc153⁺/Pifo⁺)*, and tuft cells (*Dclk1⁺/Trpm5⁺)*, as well as mural (*Acta2⁺/Actc1⁺*) mesenchyme, *Msln⁺* mesothelium, arterial (*Gja5^+^/Dll4^+^*), venous (*Slc6a2^+^/Tmem252^+^*), and capillary (*Wif1^+^/Aplnr^+^*) endothelial cells, and a variety of innate and adaptive immune components (**Fig. 1B-F; Supplemental Fig. 2D-X**). In contrast, BAL (n = 32,009 cells) was largely restricted to myeloid populations (*Pparg⁺/Prodh2⁺*) and lymphoid populations (*Cd3e⁺/Cd79b⁺*), with only a small population of epithelial cells (**Fig. 1B-D; Supplemental Fig. 1A-D**).

To generate three unique starting populations of varying compositions, PD and BAL isolates were used individually or in combination: PD_3D (100% PD), BAL_3D (100% BAL), and Mixed_3D (a 50:50 mix of BAL and PD) (**Fig. 1A**). All cell suspension mixtures were resuspended in Matrigel and maintained in identical cell culture media supplemented with broadly supportive growth factors and small molecules, including Fgf10, Egf, and inhibitors of TGFβ and GSK3 signaling and a 5% CO_2_, 37°C environment (<u>Nichane et al., 2017</u>) (**Fig. 1A**).

Organoids self-assembled in all three systems over the 10-day culture period, but each showed distinct growth kinetics and morphology (**Fig. 1G-H**). By day 3, PD_3D organoids had expanded rapidly, in both number or organoids (count) and average surface area, showing early branching with increased structural complexity. By day 7, PD_3D organoids expanded to the point of physical contact, thereby forming a dense, interconnected web-like structure. Even though organoid area continued to increase (**Fig. 1H**, **Fig. 2A**), the number of discrete organoids in this condition decreased over time, demonstrating that individual organoids fused as they grew larger. Hematoxylin and eosin (H&E) staining demonstrates that the largest structures in this condition were enriched for ATI-like cells—the thin, flattened epithelial cells responsible for gas exchange in the body (**Fig. 2K, N**).

**Figure 2.**
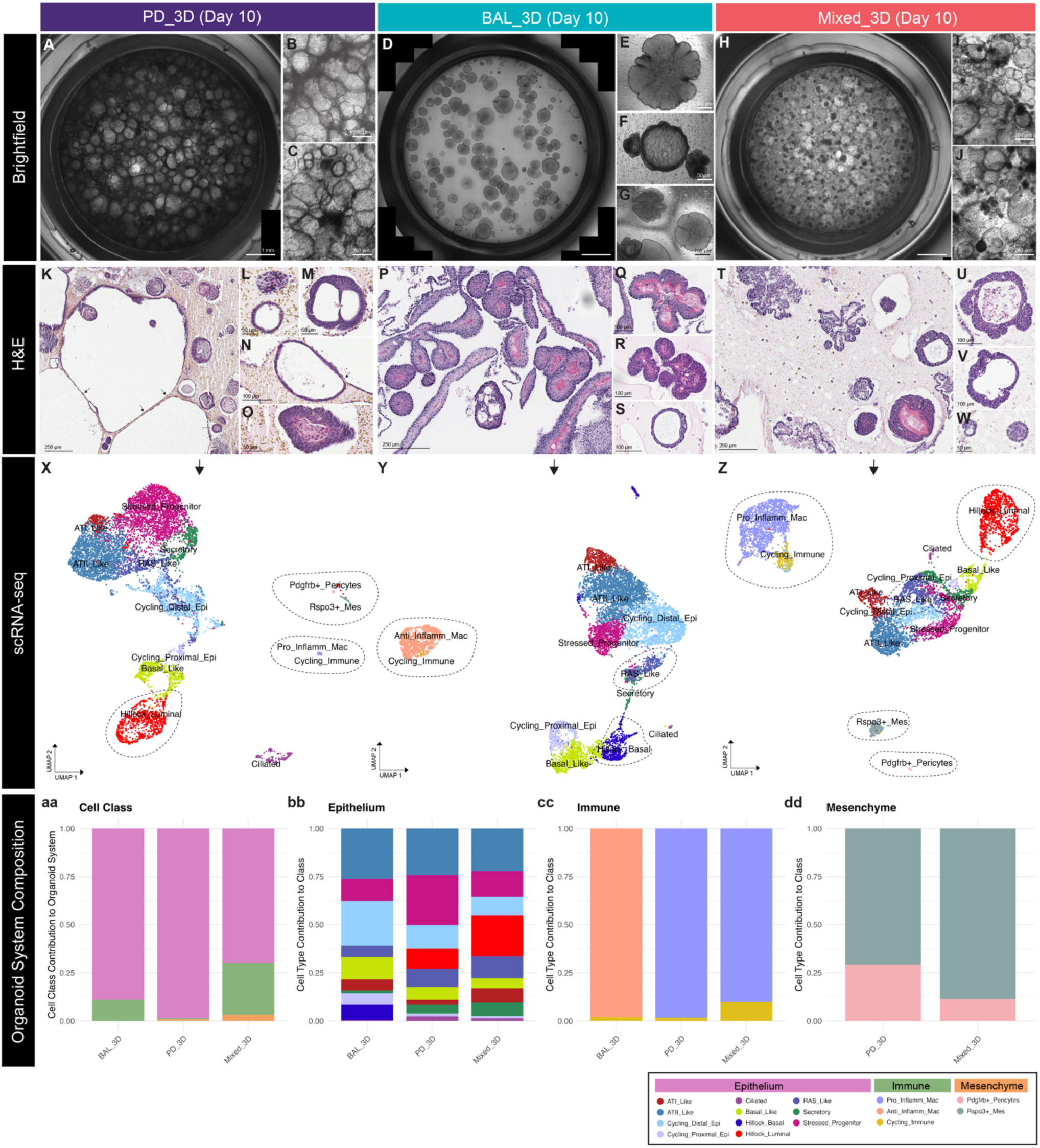
Day 10 organoid systems exhibit condition-specific morphologies, epithelial architectures, and cellular compositions. **A-C, D-F, H..J**) Brightfie1d images of representative organoid cultures on Day 10 for PD_3D **(A-C)**, BAL_3D **(D-F)**, and Mixed_3D **(H.J)**. Panels A, D, and H show stitched whole-well images (scale bar, 1mm); insets show zoomed in organoids highlighting distinct morphologies across conditions (scale bars provided on individual images due to variation). **K-M, N-O; P-R, s; T-V, W)** Hematoxylin and eosin (H&E) staining shows condition-specific epithelial architectures, inducting variations in epithelial layering, lumen formation, and structural complexity. Scale bars are sho’Nfl on each image. Arrows in Kand N highlight the thin, flattened epithelial cells morphologically consistent with alveolar type I (ATl)-like cells. **X-Z)** UMAP projections of condition-levelsingle-cell RNA-seq data for PD_30 (**X**), BAL_30 **(Y)**, and Mixed_3O **(Z)**, with cells colored by annotated identity (see *legend in figure).* Circled populations mark epithelial, immune, and mesenchymal subtypes of relevance, including Hillock cells, secretory-alveolar intermediates (AT0/RAS), polarized macrophages, Rspo3+ mesenchyme, and Pdgfrb+ pericytes. Embeddings shown using node-aligned global object cell type annotations mapped back onto the condition level embeddings. **aa-dd)** Within-class breakdowns of cell type composition across conditions on day 10. Stacked bar plots show the relative proportions of specific cell types within each major class: cell classes (aa), epithelial subsets (bb), immune subsets (cc), and mesenchymal populations (dd), normalized to 1 within each dass per condition. PD_3D and Mixed_3D cultures maintained.

In both growth kinetics and structural complexity, Mixed_3D organoids displayed an intermediate phenotype between that of PD_3D and BAL_3D structures. Counts over time were greater than in BAL_3D, but less than in PD_3D (**Fig. 1G**). Similarly, the average area of the structures increased over the culture period, though they remained smaller on average than BAL_3D by day 10 (**Fig. 1H**). Structurally, the Mixed_3D cultures contained a blend of dense, compact structures analogous to spheroids as well as larger, branched structures with early complexity comparable to PD_3D, but without the extensive fusion of structures (**Fig. 2H-J**).

Organoids in the BAL_3D system self-assembled at slower rate and these cultures produced fewer structures compared to Mixed_3D and PD_3D (**Fig. 1G-H**). However, the structures that did form were consistently larger (**Fig. 2H**). A subset of the structures in this condition displayed a floret-like morphology with branching extensions indicative of lung-budding architecture (**Fig. 2E-G**). This could indicate that the higher ratio of immune cells present and lack of mesenchyme at the start of culture in this model may have contributed to unique differentiation trajectories of the epithelium.

To relate the observed morphologies to underlying cell fate outcomes, we performed scRNA-seq on each system (condition) at Day 10 (see Methods). Lineage outputs were observably different across conditions: PD_3D and Mixed_3D maintained epithelial, immune, and mesenchymal cells whereas BAL_3D maintained only epithelium and immune cells (**Fig. 2X-Z**), the two lineages that had been present in the starting suspension. Endothelial populations were lost by Day 10 in both PD_3D and Mixed_3D, likely due to the bias of the growth medium towards the epithelium, not the endothelium. Pro-endothelial growth factors such as VEGF and FGF2 may have better supported the maintenance of the endothelial cells if they had been added to the growth medium (<u>Seo et al., 2016; Worsdorfer et al., 2019</u>) (**Fig. 1F**). *Pdgfrb⁺* pericytes and *Rspo3*⁺ fibroblasts were selectively maintained in Mixed_3D and PD_3D, in varying proportions (**Fig. 2bb, 2dd**).

BAL_3D, which lacked mesenchymal cells, had the second largest percentage of ATI-like cells BAL_3D: 5.16 ± 3.88%) after Mixed_3D (5.60 ± 1.70%), and more than PD_3D (2.68 ± 0.54%). This finding is inconsistent with previous reports demonstrating the reliance of alveolar epithelial differentiation on mesenchyme-derived signaling pathways such as Wnt, BMP, FGF, TGF-β, and YAP/TAZ (<u>Zhao et al., 2013; Jeon et al., 2022; Nabhan et al., 2022</u>). Although our data do not rule out a supportive role for mesenchyme, they indicate other cues such as immune-derived signals may be more associated with the increased ATI-differentiation observed in BAL_3D. Specifically, expression of insulin-like growth factor 1 (*Igf1*) and interleukin-1 beta (*Il1b*) were highest amongst the immune cells in BAL_3D. These findings are in line with previous work demonstrating that Igf1 can stimulate ATII-to-ATI differentiation via the activation of *Wnt5a* signaling as well as additional work showing that *Il1b* is capable of priming ATII cells for alveolar regeneration by induction toward a damage-associated state required for the eventual differentiation into mature ATI cells (<u>Ghosh et al., 2013; Choi et al., 2020</u>). Our results raise the possibility that in the absence of mesenchymal cells, immune-derived factors such as *Igf1* and *Il1b* may play a compensatory role in shaping ATI-like differentiation dynamics.

Epithelial composition and relative fractions of subtypes varied by input. PD_3D cultures generated a full spectrum of epithelial states, including ATII-like, ATI-like, basal, secretory, hillock, and RAS-like subtypes (**Fig. 2X**). BAL_3D cultures, despite beginning with only 4.8 ± 7.3% epithelial cells, gave rise to a comparable range of epithelial types (**Fig. 1C**; **Fig. 2bb**). Basal cells were most abundant in BAL_3D (9.33%) compared to PD_3D (7.1%) and Mixed_3D (5.9%). Transitional remodeling populations such as RAS-like cells and Sox9^+^ cells with stress associated transcription profiles were identified in all three systems. BAL_3D, however, uniquely gave rise to a basal-polarized hillock-like state whereas the mesenchyme-containing systems, Mixed_3D and PD_3D, produced a luminal-polarized hillock-like state (**Fig. 2bb**; **Fig. 2X, Z**).

Finally, albeit a small population, both Mixed_3D and PD_3D maintained Pdgfrb^+^ pericytes (**Fig. 2bb; 2X, Z**). Given these observations, our data underscores how initial cellular composition does not necessarily dictate epithelial diversity through generation of condition-specific lineages but by modifying the relative distribution and co-occurrence of similar cell states.

Together, these data position our organoid platform as a compositionally tunable and biologically grounded system in which divergent cell states emerge from different starting populations under the same environmental conditions. The significant expansion of epithelium from BAL samples despite their small contribution on Day 0, the survival bias towards mesenchymal and loss of endothelial populations, and the arrangement of distinct tissue morphologies and lineage distributions all highlight the ability of this platform to model how initial cellular context influences subsequent epithelial and immune specification and differentiation.

### Emergence of Polarized Hillock Cells Reveals Condition-Specific Epithelial Trajectories

Using scRNAseq data to characterize the epithelial landscape across conditions at day 10, we identified a large population of *Krt13^+^*/*Sprr1a⁺* cells co-expressing either *Krt5* or *Adm*. Given the absence of cells with the same transcriptomic profile in our starting samples, we posit that these cells arose de novo in our system, indicating their innate ability to respond to remodeling cues. In alignment with recent findings presented in <u>Montoro et al., 2018</u> and <u>Lin et al., 2024</u>, we defined this subset of epithelium as ‘Hillock-like.’ Hillock cells have been identified in the proximal airway epithelium— specifically, over the junctions between cartilage rings of the trachea and along the posterior membrane—of both mice and humans but not rat (<u>Lin et al., 2024; Otto et al., 2025</u>). ‘Hillocks’ are defined as standalone islands of *Krt13^+^* epithelial cells that form stacked structures above a layer of proliferating basal cells (<u>Lin et al., 2024</u>). These cells have been reported to exist in healthy humans and mice, but following denuded injury, they expand rapidly and display rapid turnover, contributing to regeneration of the airway epithelium (<u>Montoro et al., 2018; Lin et al., 2024; Otto et al., 2025</u>).

To validate the significance of our findings in organoid culture, we performed immunohistochemical (IHC) staining for *Krt13* on longitudinally sectioned native rat trachea, where hillock cells have not previously been described. We identified organized clusters of *Krt13^+^* cells at both the basement membrane (basal) and atop the cells at the basement membrane (luminal) (**Fig. 3B-C**). These structures mimicked the characteristic anatomy of hillock domains seen in mouse and human airways: a layer of stratified squamous *Krt13^+^* cells overlaying a basal stem cell pool expressing *Krt5* (<u>Montoro et al., 2018; Lin et al., 2024</u>). To confirm the presence of these cells in vitro, as well as the protein-level, as opposed to transcript-level, expression of *Krt13*, we performed additional IHC staining on day 10 organoids and identified discrete foci of similarly layer populations that were either *Krt13^+^/Krt14^+^*(basal hillock) or *Krt13^+^/Scgb1a1^+^* (luminal hillock) (**Fig. 3J-O**). All hillock-like cells in 3D culture simultaneously expressed *Sprr1a*, a member of the small proline-rich (*Sprr*) protein family, alongside *Krt13*. We highlight this finding as *Sprr1a* has been implicated in terminal squamous differentiation and the formation of the cornified envelope in keratinocytes (<u>Zimmerman et al., 2004; Zheng et al., 2008; Strunz et al., 2020</u>). Therefore, the co-expression of *Krt13* and *Sprr1a* by these cells supports a model in which they are engaging in mature squamous differentiation program.

**Figure 3.**
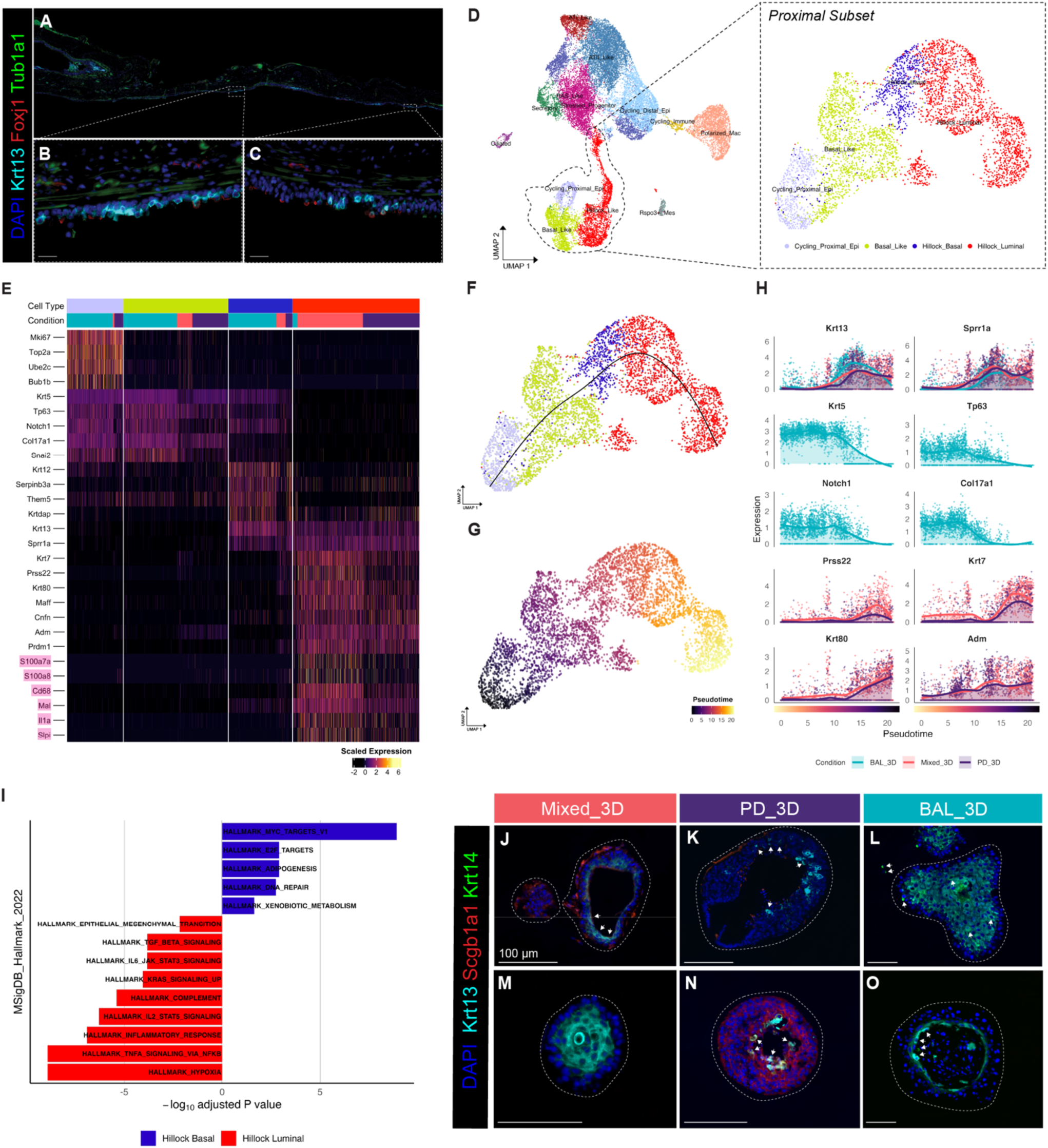
Hillock cells emerge as a polarized epithelial state with distinct differentiation trajectories and immune-regulatory features. **A-C)** lmmunofluorescence staining of native rat trachea shows clusters of Krt13+ squamous cells (cyan) with Tuba1a (green) and Foxj1ciliated cells (red). These stratified structures resemble hillock domains described in mouse and human airways. Scale bars, 50 µm. **D)** UMAP of the integrated Day 10 single-cell dataset, combining all organoid sequencing data. The right panel shows the subsetted proximal epithelial population, used for downstream trajectory inference and marker analysis. Hillock cells are subdivided into Hil1ock_Basal (Krt13+/Krt5+) and Hillock_Luminat (Krt13+/Krt5-) populations. **E)** Heatmap of marker gene expression across hillock subtypes, including squamous markers (Krt13, Sprr1a), basal markers (Krt5, Krt14), and immune-associated genes highlighted in pink. **F-G)** Slingshot pseudotime analysis of the proximal epithelial subset reveals a trajectory from cycling progenitors through Basal_Like, Hillock_Basal, and ultimately to Hillock_Luminal cells. **H)** Gene expression trends across pseudotime show stepwise activation of differentiation and immune programs. **I)** MSigOB Hallmark enrichment analysis comparing Hillock_Luminat and Hillock_Basal populations. **J-O)** lmmunofluorescence staining of day 10 organoids shows condition-specific spatial organization of hillock cells. In Mixed_30 **(J, M)** and P0_30 **(K, N)**, Krt13+/Krt14-luminal cells form flattened, apical epithelial layers. In BAL_30 (**L, 0**), Krt13+/Krt14+ basal-like cells assemble into multilayered squamous structures (white arrowheads). A subset of hillock luminal cells co-express Scgb1a1 (white arrowheads), supporting a luminal hillock identity. Scale bars, 100 µm.

Sub-clustering analysis our sequencing data showed that these *Krt13*⁺ cells clustered distinctly within the proximal epithelial subset (**Fig. 3D**) and could be further subdivided into two transcriptionally polarized populations: hillock basal (*Krt13⁺/Krt5⁺/Sprr1a⁺*) and hillock luminal (*Krt13⁺/Sprr1a⁺/Krt5^−^*), in line with previously characterized hillock subtypes in the murine and human lung (<u>Lin et al., 2024; Montoro et al., 2018; Otto et al., 2025</u> ). Interestingly, we found that the relative proportions of each subtype were condition-specifically polarized: BAL_3D cultures derived from am immune-rich, mesenchyme-poor starting population exhibited an abundance of hillock basal cells whereas Mixed_3D and PD_3D conditions, derived from a more compositionally diverse starting population, favored the hillock luminal phenotype (**Fig. 3D**). Therefore, we propose that the immune-rich context of BAL_3D favors a basal-skewed hillock state whereas epithelial-mesenchymal heterogeneity promotes luminal differentiation (**Fig. 3E**). Given that the initial epithelial compartment in BAL largely mirrors that of PD, albeit at a much smaller scale (**Supplemental Fig. 1b, d; Supplemental Fig. 2A, D**), these results suggest that microenvironmental cues, rather than epithelial-intrinsic differences, are major drivers of hillock subtype polarization.

To understand how these cells differentiate, we used Slingshot pseudotime analysis. From this analysis, we found these cells progress along a well-defined trajectory from proximal cycling epithelial cells through canonical basal cells (*Krt5^+^/Sprr1a^+^/Krt13^−^*) to basal-hillock-like (*Krt5^+^/Sprr1a^+^/Krt13^+^*), ultimately terminating in the luminal hillock state (*Krt5^−^/Sprr1a^+^/Krt13^+^*) (**Fig. 3F-G**). Earlier findings confirm this trajectory as they suggest that the more squamous, hillock luminal cells may be derived from hillock basal cells and thus, the luminal cells represent a more differentiated or specialized form of the hillock lineage (<u>Lin et al., 2024</u>).

In parallel with this trajectory, we found that hillock luminal cells also upregulate a distinct suite of immune-associated genes, including *Slpi, Mal, S100a7, Il1a, and S100a8* (**Fig. 3e**). The aforementioned genes are classically linked to antimicrobial defense, damage sensing, and inflammatory signaling at barrier surfaces. These functions are increasingly attributed to epithelial cells in injury and regenerative contexts (<u>Hewitt & Lloyd, 2021</u>).

Given evidence suggestive of the immunological roles of epithelial cells in repair contexts, examining the functions of the immune-associated genes expressed by the hillock-like cells offers insight into their potential contributions in our systems and disease contexts. *Slpi*, expressed by luminal hillock cells in the Mixed_3D and PD_3D systems, is a multifunctional protein with anti-protease, anti-inflammatory and antimicrobial properties (<u>Brown et al., 2024; Osbourne et al., 2022; Nishimura et al., 2008</u>). As a serine-protease inhibitor, *Slpi* blocks the activity of proteases in an effort to protect the airway epithelium from the proteases released by neutrophils during inflammation (<u>Brown et al., 2024; Camper et al., 2015</u>). *Il1a*, a pro-inflammatory cytokine, is rapidly produced by epithelial cells in response to stress, necrosis, or damage, where it acts as an alarmin to recruit and activate immune cells (<u>Suwara et al., 2014</u>). In addition, *Mal*, a proteolipid involved in mediating apical sorting among polarized epithelial cells, ensuring the proper delivery of proteins to the apical membrane, as well as facilitating Lck trafficking in T cells for TCR-immune activation (<u>Cao et al., 2012; Anton et al., 2008</u>).

*S100a7* (psoriasin) and *S100a8*, both calcium-binding proteins, play a similar role as *Il1a* in that they are considered alarmins themselves. S100a8 has been identified to induce the production of *Muc5ac* by airway epithelial cells in COPD as well as activating alveolar epithelial cells via TLR-4 to prompt the release of various chemokines and cytokines (<u>Kang et al., 2014; Chakraborty et al., 2017</u>). *S100a7*, on the other hand, is constitutively expressed in bronchial epithelial cells, however, is upregulated in response to bacterial infection (<u>Andresen et al., 2011</u>). The expression of such genes by luminal hillock suggests that these polarized epithelial cells may uniquely contribute to epithelial-immune communication, serving not merely as transitional intermediates but as immunomodulatory effectors within the remodeling airway epithelium (<u>Katsura et al., 2019</u>).

We further explored the functional specialization within the hillock population by comparing luminal hillock cells to basal hillock cells via MSigDB Hallmark pathway enrichment analysis (<u>Liberzon et al., 2015</u>). Luminal hillock cells were significant enriched for pathways related to inflammatory signaling and immune communication (TNFα signaling via NF-κB, IL2/STAT5 signaling), as well as stress adaptation (complement, TGFβ signaling, and hypoxia) (**Fig. 3I**). In contrast, basal hillock cells were enriched for processes related to proliferation (MYC targets, E2F targets, and G2M checkpoint), consistent with a more canonical basal epithelial state (**Fig. 3I**) (<u>Dong et al., 2014</u>).

The signaling profile of hillock luminal cells, characterized by high expression of cytokines and alarmins along with activation of inflammatory pathways, suggests that they may adopt a regulatory epithelial state. Drawing on the framework proposed by Medzhitov (<u>2021</u>), this pattern is consistent with an epithelial response to loss of regulatory control, in which classical homeostatic mechanisms are insufficient, and inflammation is repurposed to enforce system-level coordination.

To validate these transcriptional and pathway-based findings, we also more deeply studied the spatial distribution of hillocks using immunofluorescence staining of day 10 organoids across BAL_3D, PD_3D, and Mixed_3D conditions. We found that in BAL_3D, *Krt13^+^* cells were of a multilayered squamous appearance and were localized throughout the apical (**Fig. 3I**). Alternatively, *Krt13^+^* cells in Mixed_3D and PD_3D exhibited a flattened, apically oriented morphology with less extensive coverage of the epithelial surface (**Fig. 3J, K, N**). Of note, in both PD_3D and Mixed_3D organoids, some luminal cells also co-expressed *Scgb1a1*, consistent with their transcriptional signature and suggesting that *Krt13^+^*/*Scgb1a1^+^* cells represent a differentiated luminal hillock state (**Fig. 3J, N**).

Collectively, these results demonstrate that hillock-like cells spontaneously emerge from native lung-derived organoid cultures and are either stalled in a more basal-hillock progenitor like state or become more differentiated, taking on a luminal-hillock signature. We propose that these differentiation events are associated with the cellular composition of the starting population. These cells also exhibit distinct spatial patterning, differentiation trajectories, and immune-associated gene expression consistent with their proposed dual role in epithelial remodeling and immune coordination. Therefore, hillock cells are not only an example of epithelial plasticity but also provide insight into the multicellular dynamics that underlie airway regeneration in response to inflammation.

### Culture Composition Dictates Divergent Macrophage Activation Programs

To better understand how the immune compartment changed over the course of culture, we next examined the persistence and spatial dynamics of immune cells across conditions. While most immune populations were completely absent in our organoid cultures by day 10 of culture, a population of activated macrophages represented an exception. These macrophages were maintained in all three engineered conditions, with the most consistent maintenance in the BAL_3D and Mixed_3D cultures where the initial (Day 0) ratio of immune cells to all other cells was greatest (**Fig. 2aa; Supplemental Fig. 1A; Supplemental Fig. 2A**). Spatially, the macrophages remained dispersed in the extracellular matrix rather than embedded within organoid structures or contain within luminal spaces (**Fig. 4E**). They were spatially segregated from epithelial cells, indicating that their effects likely were facilitated via paracrine signaling rather than cell-cell contact (**Fig. 4E**).

**Figure 4.**
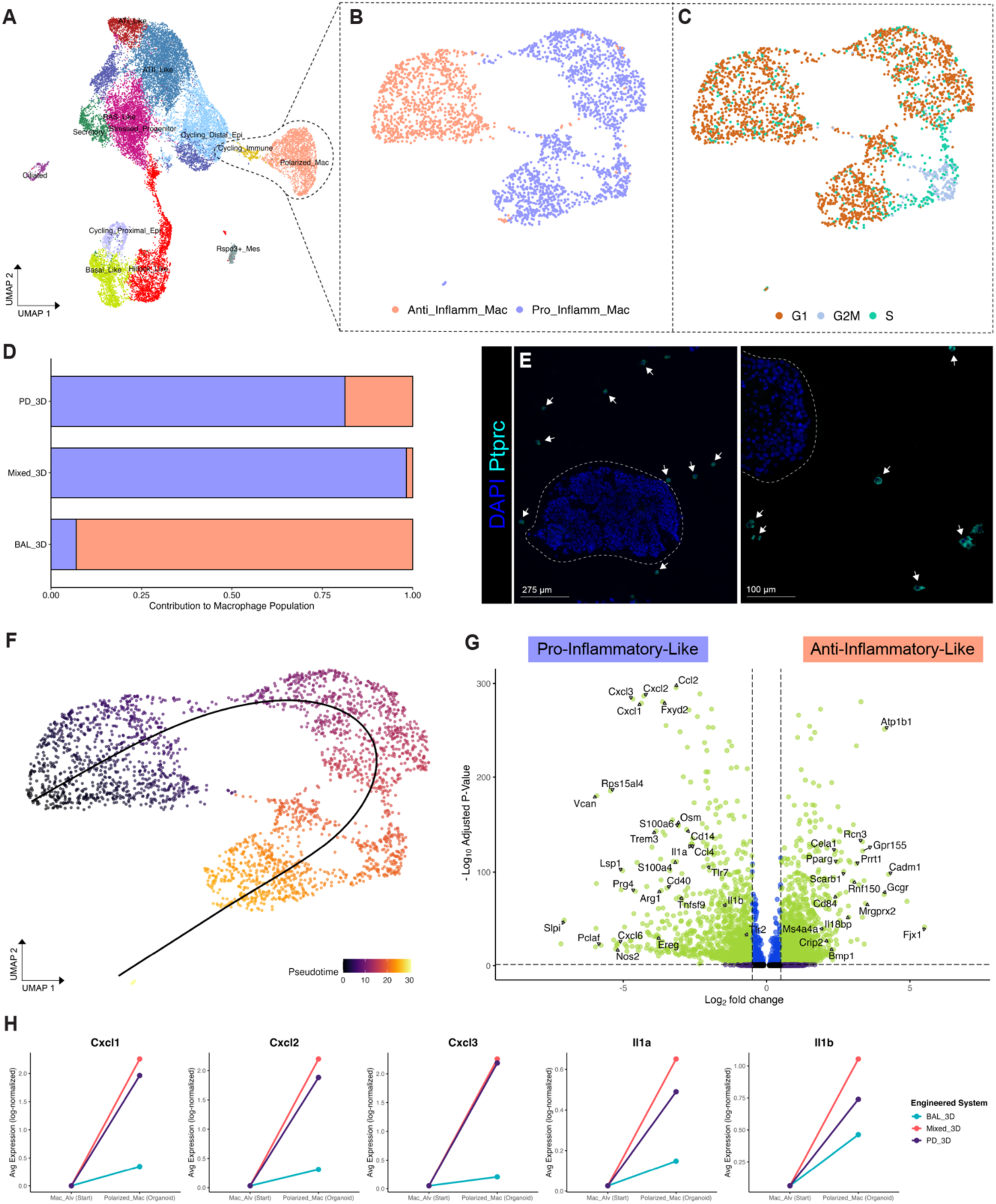
Macrophage polarization trajectories reveal condition-specific immune modulation. **(A)** UMAP of the integrated day 10 organoid dataset highlighting macrophage populations, which cluster as a transcriptionally distinct immune compartment within the broader dataset. **(B)** UMAP of macrophage subset, annotated into two transcriptionally distinct states: anti-infiammatory-like (salmon) and pro-inflammatory-like (lavender). **(C)** Cell cycle phase scoring of macrophages reveals that while most cells are in G1, a subset of pro-inflammatory-like macrophages are cycling, with enrichment in Sand G2/M phases. **(D)** Stacked bar plot showing condition-specific contributions to the macrophage population. BAL_3D cultures are enriched for anti-infiammatory-like macrophages, whereas PD_3D and Mixed_3D conditions contain a higher proportion of pro-inflammatory-like macrophages. **(E)** lmmunofluorescence staining of day 10 BAL_3D cultures show Ptprc+ alveolar macrophages (cyan) dispersed in the surrounding matrix, often adjacent to but spatially distinct from epithelial organoids (outlined by dashed lines). **(F)** Pseudotime trajectory inferred using Slingshot reveals a transcriptional continuum from anti-inflammatory-like to pro-inflammatory-like macrophage states. (G) Volcano plot showing differentially expressed genes between pro- and anti-inflammatory-like macrophages. Pro-inflammatory-like macrophages express elevated levels of inflammatory and alarmin genes (e.g., Cxcl2, Trem1, 111a, S100a8), while anti-inflammatory-like macrophages upregulate genes associated with tissue remodeling and resolution (e.g., Pparg, Ms4a4a, Cela1). **(H)** Changes in average gene expression (log-normalized) between alveolar macrophages in the starting populations and polarized macrophages in the three organoid systems.

To further characterize macrophage heterogeneity, we subclustered the immune compartment and identified three transcriptionally distinct macrophage clusters (**Fig. 4A**). To investigate whether the macrophages that persisted across conditions adopted distinct activation states, we performed differential gene expression analysis on this subset, comparing cluster 0 (enriched in BAL_3D) to clusters 1 and 2 (enriched in Mixed_3D and PD_3D). The resulting transcriptional programs did not fully align with classical M1/M2 macrophage paradigms but instead resembled anti-inflammatory-like and pro-inflammatory-like states (**Fig. 4A-B**) (<u>Guan et al., 2023</u>).

Anti-inflammatory-like macrophages, which predominated in BAL_3D, expressed a coordinated program of immune regulatory and tissue-adaptive genes, including *Pparg*, *Cd84*, *Cadm1*, and *Cela1* (**Fig. 4G**). *Pparg*, a nuclear receptor known to repress inflammatory gene expression, likely acts upstream to promote alternative activation and lipid handling (<u>Nolte et al., 1998; Neri et al., 2011</u>). *Cd84*, a SLAM family receptor, may reinforce this anti-inflammatory response by modulating NF-κB signaling during inflammation (<u>Sintes et al., 2010)</u>. *Cadm1* and *Cela1* further point to tissue adaptation: *Cadm1* has been implicated in macrophage-epithelial interactions, while *Cela1*, although canonically epithelial, contributes to extracellular matrix remodeling and has been detected in macrophages during lung development (<u>Ramos et al., 2022; Joshi et al., 2018</u>). Taken together, these data suggest that macrophages in BAL_3D, lacking mesenchymal signals, not only play immunoregulatory roles but may be involved in restructuring the physical properties of the engineered niche.

Pro-inflammatory macrophages, which were enriched in Mixed_3D and to a lesser extent PD_3D, upregulated genes associated with acute inflammatory signaling, leukocyte recruitment, and matrix remodeling (**Fig. 4G**). These cells showed a marked increase expression of several potent neutrophil chemoattractants such as *Cxcl1*, *Cxcl2*, and *Cxcl3*. These chemokines are known to signal through *Cxcr2*, which interestingly, was not expressed by any of cells in our global engineered dataset (<u>Thomas et al., 2025; Russo et al., 2008; Belperio et al., 2002</u>). However, more recent findings propose that when *Cxcr2* is absent, Cxcl1 and Cxcl2 can operate through an alternative pathway, involving the p38 MAPK pathway and operating through *Cxcr1*, albeit with a lower binding affinity compared to *Cxcr2* (<u>Boon et al., 2025</u>). Unlike *Cxcr2*, *Cxcr1* was expressed by a smaller subset within our activated macrophage population, thereby suggesting that the pro-inflammatory macrophage upregulating *Cxcl1/2/3* were operating via an autocrine loop to elicit their pro-inflammatory effects.

Concurrent expression of *Il1a* and *Nos2* by the pro-inflammatory macrophage subset reflects classical M1-like features: *Il1a* being a master cytokine for amplifying local inflammation, and *Nos2* catalyzing nitric oxide production, contributing to microbicidal activity and tissue damage (**Fig. 4G**) (<u>Raja et al., 2024; Palmieri et al., 2020</u>).

Supporting our findings that the macrophages in PD_3D and Mixed_3D assumed a pro-inflammatory signature, a subset of these macrophages were inferred to be in the G2M and S phases, according to cell cycle scoring performed in Seurat (**Fig. 4C**). This observation suggests that these cells were undergoing continuous cell cycle progression, which is a feature that has previously been associated with activated-induced expansion of inflammatory macrophage populations (<u>Micochova et al., 2024)</u>.

This pro-inflammatory program was more common in the systems that maintained mesenchymal cells (Mixed_3D, and to a lesser degree, PD_3D, but not BAL_3D) (**Fig. 2aa**; **Fig. 4D**), suggesting that mesenchymal-derived cues could drive macrophage inflammatory polarization, even under otherwise similar culture conditions.

Building on our previous findings that macrophage states varied across organoid systems, we performed pseudotime analysis on the immune subset to further characterize the relationship between these states and their system-level specificity (**Fig. 4F**). The analysis revealed a continuous trajectory from the anti-inflammatory cluster to the pro-inflammatory cluster. Again, given our findings that the anti-inflammatory macrophages were unique to the BAL_3D system that lacked mesenchyme, pseudotime analysis reinforces our interpretation that mesenchyme-derived systems drive differentiation (**Fig. 4D**). Like the transcriptomic polarization patterns we observed in epithelial hillock cells, the less-differentiated basal-hillock cells predominated in BAL_3D, as well, while the more differentiated luminal hillock cells were enriched in the mesenchyme-containing systems, PD_3D and Mixed_3D. The continuous trajectory in pseudotime, combined with the system-specific distribution of cell states, suggests that these populations may not represent fixed or terminal identities but instead exist along an activation spectrum and that mesenchymal signals are critical in driving such activation. Without mesenchymal signals, we hypothesize that macrophages in our system remain in an anti-inflammatory state.

### Sox9⁺ Transitional States and RAS-Like Cells Define Alternative Regenerative Programs

Injury-associated epithelial progenitor states have been identified as important regulators of distal lung regeneration and when disrupted, are major mediators of disease (<u>Alysandratos et al., 2021</u>). In conditions such as idiopathic pulmonary fibrosis (IPF) and chronic obstructive pulmonary disease (COPD), *Sox9^+^* progenitors and secretory-biased intermediates deviate from canonical alveolar differentiation pathways to participate in distal epithelial repair and facilitate alveolar remodeling (<u>Cai et al., 2024; Basil et al., 2022</u>). Although *Sox9⁺* progenitor cells are present at low levels in the healthy lung, their expansion is associated with alveolar injury, aberrant regeneration, and impaired lineage resolution (<u>Sun et al., 2022; Cai et al., 2024</u>).

In human distal lung biology, two transitional populations bridging canonical secretory and alveolar identities have been recognized: respiratory airway secretory (RAS) cells (<u>Basil et al., 2022</u>) and alveolar type 0 (AT0) cells (<u>Murthy et al. (2022)</u>. RAS cells have been defined as a unique secretory population that exist in the respiratory bronchioles of humans and ferrets and co-express *SCGB3A2* and *SCGB1A1* (<u>Basil et al., 2022</u>). Upon injury, these cells become activated and express *SFTPC*, in addition to *SCGB3A2* and *SCGB1A1*, demonstrating their transcriptional proximity to ATII cells as well as canonical secretory populations (<u>Basil et al., 2022</u>). Upon activation by injury or in organoid models, <u>Basil et al., 2022</u> observed RAS cells to quickly upregulate *SFTPC* while simultaneously downregulating *SCGB3A2*, indicating their differentiation toward a more canonical ATII fate.

AT0 cells, described by <u>Murthy et al., 2022</u>, are similar to RAS cells, but do demonstrate distinct differences worth describing. While RAS cells have been identified in human and ferret lungs, AT0s have been found in human and primate (rhesus macaques) lungs. What makes AT0s different from RAS cells is that AT0s are thought to be derived from ATII cells upon injury whereas RAS cells are positioned as a specialized type or secretory cell. When injury occurs, <u>Murthy et al., 2022</u> propose that ATII cells shift to the AT0 state where they then can differentiate into one of two cell types: ATI or terminal respiratory bronchiole secretory cells (TRB-SCs). Additionally, AT0s are defined by their co-expression of *SCGB3A2*, *SCGB1A1*, and *SFTPB*, not *SFTPC* (<u>Murthy et al., 2022</u>). In both cases, however, both RAS and AT0 cells are associated with sites of active epithelial remodeling (<u>Murthy et al., 2022; Basil et al., 2022</u>).

Prior to the identification of RAS and AT0 cells, the bronchioalveolar stem cell (BASC), discovered by <u>Kim et al. (2005)</u>, was thought to be the primary epithelial progenitor population in the distal lung—first characterized as a bipotent progenitor located at the bronchioalveolar duct junction (BADJ) in the mouse lung and express both *Scgb1a1* and *Sftpc*. The physiological location and gene expression profile of these rare cells were indicative of their ability to give rise to both bronchiolar (secretory) and alveolar (ATII) lineages (<u>Kim et al., 2005</u>). With further study, these cells have been shown to be resistant to epithelial injury, proliferative following damage, and capable of self-renewal and multilineage differentiation in vitro and in vivo (<u>Kim et al., 2005; Salwig et al., 2019</u>). However, their human orthologues are still unknown, and evidence suggests that the more recently discovered progenitor intermediates such as RAS and AT0 cells may be functionally equivalent to or even replace BASCs during regeneration and disease (<u>Hoffman et al., 2022</u>). Most importantly, findings by <u>Martins et al., 2024</u> imply that a single transitional epithelial subtype does not single-handedly drive lung regeneration or take over the role of the BASC. Rather, it is more likely that the multiple intermediate epithelial cell types can become activated upon injury and when this occurs, together, they play complementary roles to facilitate repair (<u>Martins et al., 2024</u>). These insights highlight how different regenerative strategies are used by the distal epithelium depending on the exact contexts of injury.

Consistent with results reported by others, we identified *Sox9*⁺ cells in the native adult rat lung near the BADJ and occasionally co-expressing *Scgb1a1*, as expected of this bipotent progenitor population (**Fig. 5A**). We also identified *Scgb3a2*⁺ secretory cells concentrated in the conducting airways, mirroring the spatial distributions similar to those reported in human and ferret respiratory bronchioles (**Fig. 5B**). In native adult rat lungs not exposed to injury cues, the *Scgb3a2*⁺ cells were not observed to express Sftpc, as to be expected (**Fig. 5B**). Rather, Sftpc expression appears restricted to canonical ATII cells in the alveolar sacs (**Fig. 5B**).

**Figure 5.**
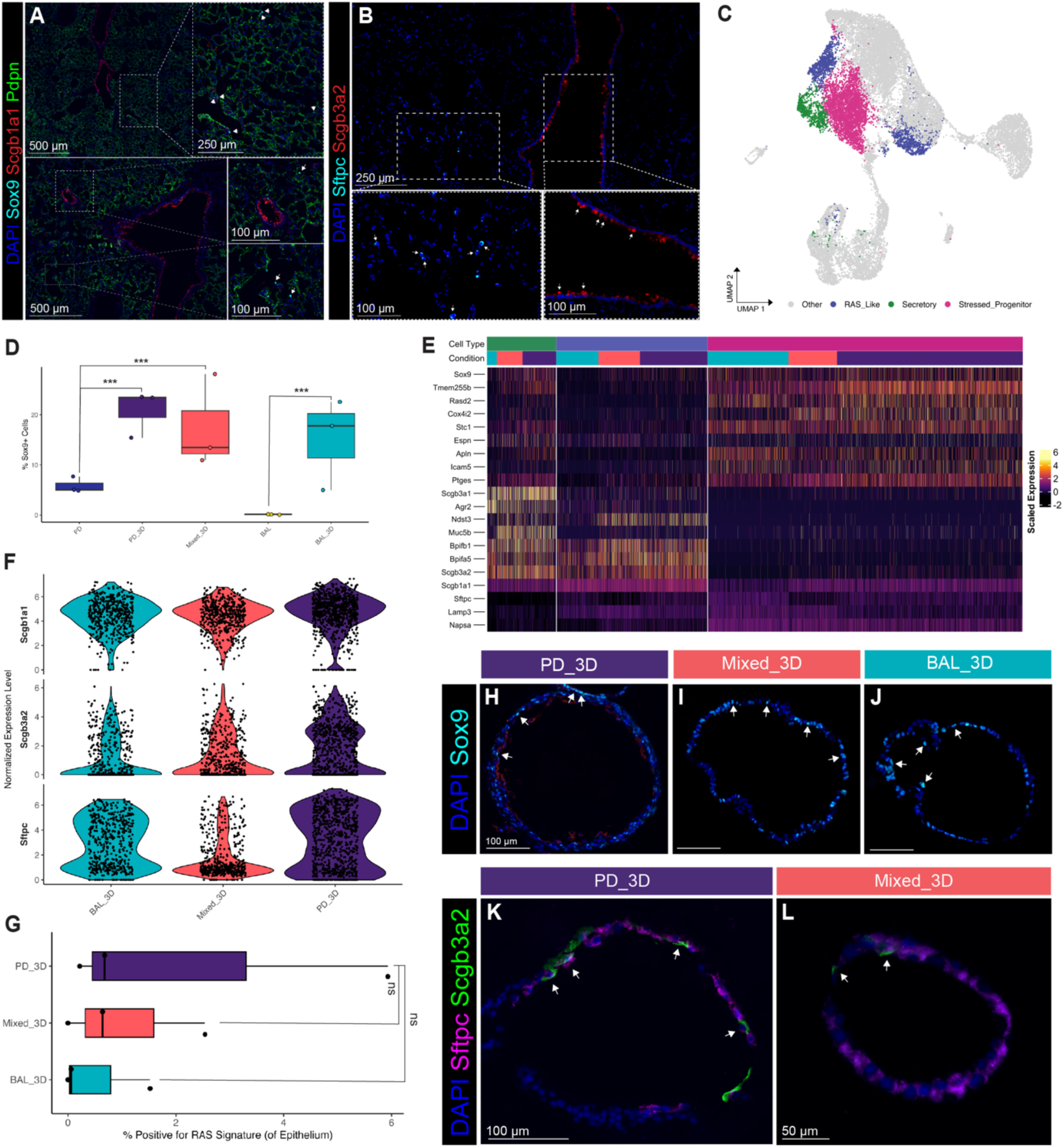
Mesenchyme-rich conditions promote the emergence of secretory-alveolar intermediates, while Sox9+ stressed progenitors emerge across all engineered systems. **(A-B)** lmmunofluorescence staining of native adult rat lung reveals rare epithelial progenitor and secretory populations. **(A)** Sox9+ cells (cyan) are observed near the bronchoalveolar duct junction (BADJ), occasionally co-expressing Scgb1a1 (red). Pdpn (green) marks alveolar type I (ATI) cells. **(B)** Scgb3a2+ secretory cells are localized to the conducting airways. Arrowheads highlight individual positive cells. (C) UMAP of the integrated day 10 organoid dataset highlights transcriptionally distinct transitional populations, including aRASIAT0-like, stressed progenitor (Sox9+), and canonical secretory subsets. **(D)** Boxplot quantifying the percentage of Sox9+ cells (threshold: Sox9 expression > 0.1) across five conditions (PD, BAL, PD_3D, Mixed_3D, BAL_3D). Sox9+ cells are rare in the starting populations but expand markedly after 10 days of culture in all conditions. Each point represents an individual biological replicate. Statistical comparisons were performed using two-proportion Z-tests on raw Sox9+ counts, corrected using the Benjamini-Hochberg method (***p < 0.001). **(E)** Scaled heatmap showing differential gene expression across transitional epithelial subtypes. aRASIAT0-like cells (Scgb3a2+/Scgb1a1+/Sftpc+) are distinct from stressed progenitors, which express stress-associated markers (Cox4i2, Ptges, Stc1) alongside the progenitor marker Sox9. **(F)** Violin plots showing expression of signature markers distinguishing canonical secretory and activated RAS-like epithelial populations, split by condition. (G) Boxplot showing the percentage of epithelial cells per replicate positive for a RAS-like signature (Scgb1a1+/Scgb3a2+/Sftpc+) across engineered conditions. Signature positivity was determined using scaled expression thresholds: Scgb1a1 > 2, Scgb3a2 > -2.5, and Sftpc >-1.5. Triple-positive cells were observed across all three conditions. Statistical comparison via a Kruskal-Wallis test revealed no significant differences in overall prominence; however, the PD_3D system exhibited greater replicate variability, as demonstrated by its wider interquartile range. **(H.J)** lmmunofluorescence staining of day 10 organoids reveal Sox9+ epithelial cells (cyan) variably distributed across all conditions. Scale bars: 100 µm. **(K-L)** lmmunofluorescence staining of day 10 organoids (PD_3D and Mixed_3D) showing co-expression of Scgb3a2 (green) and Sftpc (magenta) (RAS-like cells). Scale bars: 100 µm (K), 50 µm (L).

Following 10 days in culture, we observed an *Scgb1a1^+^/Sftpc^+^/Sox9^+^*-expressing progenitor-like population in all three organoid systems (**Fig. 5D, H-J**). While these cells were present in our PD and BAL starting populations, they were not nearly as abundant (relative fraction of sample) (**Fig. 1B-C**). Interestingly, after 10 days in organoid culture, adopted the expression of stress-associated genes including *Cox4i2*, *Ptges*, and *Stc1,* in addition to the already present expression of BASC signature genes (**Fig. 5E**). The co-expression of BASC signature markers and stress-associated genes suggests that these Sox9^+^ transitional cells may be an injury-responsive intermediate state reflecting the intrinsic plasticity and extrinsic cues within our controlled culture conditions (**Fig. 5C, E**).

While *Sox9* reportedly marks adult epithelial progenitors that regenerate alveoli, it’s expression is also localized to distal tip progenitors in the fetal lung (<u>Sun et al., 2022; Rockich et al., 2013; Cai et al., 2024</u>). Conversely, *Cox4i2*, a nuclear subunit of cytochrome c oxidase (complex IV of the electron transport chain), is present in healthy lung tissue as it is critical for maintaining respiratory function and energy metabolism. However, epithelial cells in hypoxic conditions have been shown to increase their expression of *Cox4i2* (<u>Aras et al., 2013; Hüttemann et al., 2012</u>). Therefore, we propose that these progenitors experienced cell stress (hence their naming, ‘Stressed_Progenitors’) and likely underwent an initial expansion to repair sense damage in organoid culture but then needed to undergo a metabolic shift for survival. In this context, the upregulation of *Cox4i2* may facilitate an alternative path for ATP generation that is more efficient under the unique conditions created by the organoid microenvironment (<u>Aras et al., 2013; Rockich et al., 2013</u>).

In parallel, the upregulation of *Ptges* and *Stc1* may further indicate that such stressed *Sox9^+^* progenitors are not only expanding to serve as a reserve pool of multipotent progenitors but actively contributing to the modulating repair of the system. *Ptges*, involved in PGE2 synthesis, has strong links to modulating inflammation while *Stc1*, a secreted glycoprotein hormone, has been shown to activate anti-apoptotic signaling in injury contexts (<u>Block et al., 2009; Liu et al., 2012</u>).

We identified an activated RAS-like epithelial state, as defined by the co-expression of *Scgb1a1*, *Scgb3a2*, and *Sftpc*, across all three organoid systems, but the fraction of the epithelium that they represented varied significantly. These cells were most abundant in the PD_3D and Mixed_3D conditions where mesenchyme was preserved (as demonstrated by both RAS signature scoring and expression of individual genes, **Fig. 5F-G, Supplemental Fig. 7D, H, I**). In addition to the RAS-like transitional state, we also identified canonical secretory cells (*Scgb1a1⁺*/*Scgb3a2⁺/Sftpc⁻*) across all three systems, but unlike the RAS-like cells, these cells did not express alveolar-associated genes (e.g. *Sftpc, Napsa, Lamp3*). Organoid-inherent cues aside, the RAS-like intermediate population was not starting population specific nor did it require exogenous injury to emerge, suggesting that epithelial-mesenchymal-immune interactions affect the balance between transitional and differentiated epithelial states under homeostatic culture conditions.

### Intra-Lineage Signaling Coordinates Epithelial, Immune, and Mesenchymal Regenerative Response

Understanding how diverse cellular inputs shape epithelial and immune behavior requires not only classifying cell states but also mapping the signaling cues that guide them. Although our previous analyses suggested a condition-dependent emergence of specific macrophage and hillock phenotypes, the mechanisms coordinating these shifts remained difficult to elucidate at the transcriptomic level. With an understanding that mesenchymal cells facilitate paracrine signaling, producing signals that affect both epithelial differentiation and immune polarization, we hypothesized that variation in the presence of mesenchyme modulates these outcomes by regulating context-specific signaling programs (<u>Enomoto et al., 2023; Manneken & Currie, 2023; Negretti et al.,</u> 2025).

We therefore employed NICHES (*Niche Interactions and Communication Heterogeneity in Extracellular Signaling*), an algorithm that uses single-cell transcriptomic data to quantify the ligand-receptor signaling potential between individual cells at the pairwise level (Raredon et al., 2023). Through this analysis, we were able to resolve how communication within and across epithelial, mesenchymal, and immune compartments changes under different engineered conditions (**Fig. 6A**). We identified signaling axes upregulated in mesenchyme-rich (Mixed_3D, PD_3D) versus mesenchyme-poor (BAL_3D) contexts, including many involving pro-regenerative, pro-inflammatory, and immunomodulatory ligands (**Fig. 6C**).

**Figure 6.**
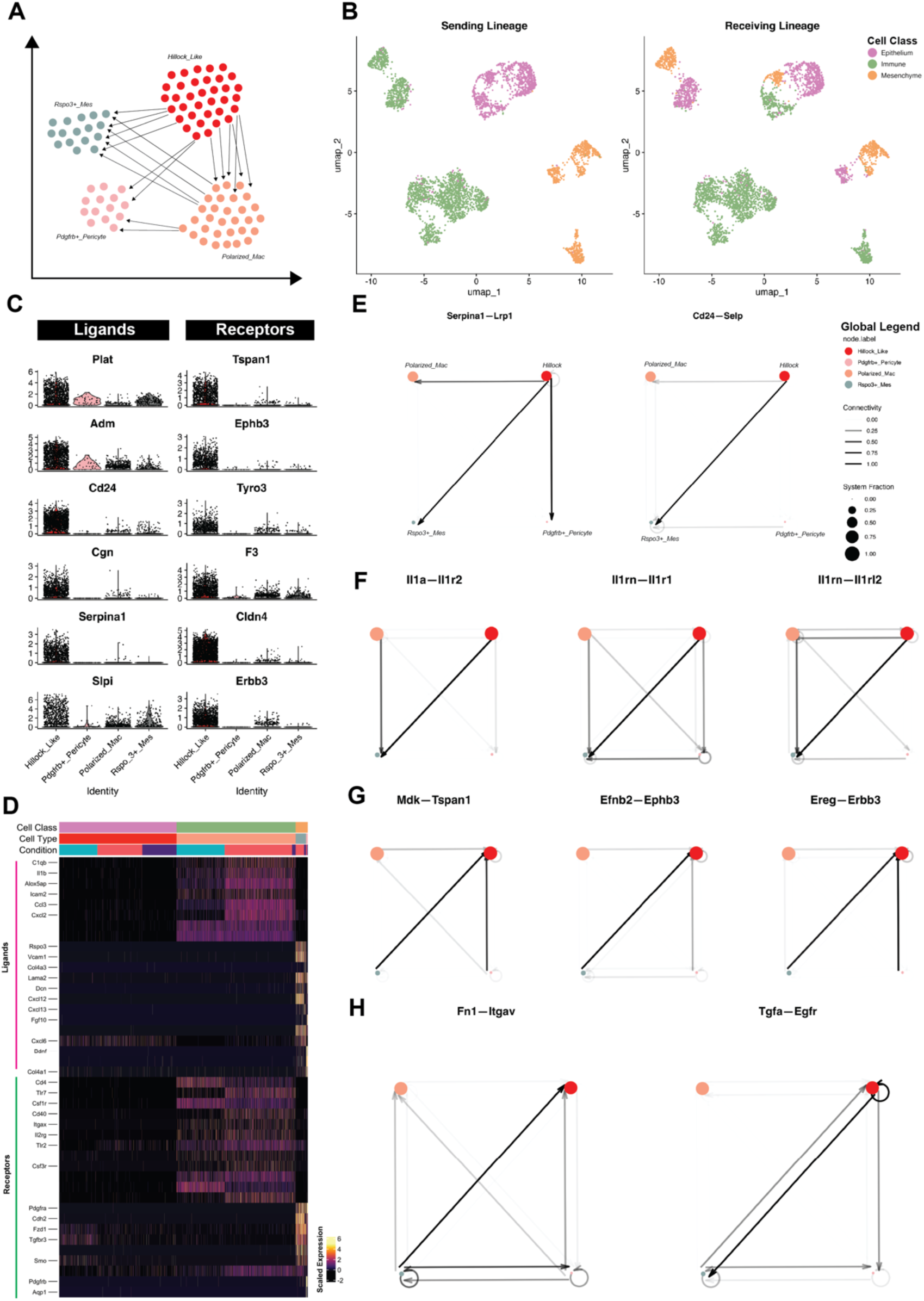
Hillock cells participate in reciprocal signaling circuits that coordinate mesenchymal and immune inputs. **A)** Schematic summary of ligand-receptor interactions between cell populations of interest, inferred using NICHES. Arrows represent directional communication among Polarized_Mac, Pdgfrb+_Pericyte, Rspo3+_Mes, and Hillock_Uke epithelial cells. Hillock cells act as central orchestrators in these networks, orchestrating and responding to diverse mesenchymal and immune-derived signals. **B)** UMAPs showing sending (left) and receiving (right) lineages used for analysis of cell-cell connectivity (NICHES output). **C)** Violin plots showing expression of candidate ligands and receptors enriched in hillock cells. **D)** Scaled heatmap of ligand and receptor expression across cell types and conditions, highlighting transcriptional diversity of connectomic inputs. **E-H)** Circuit plots illustrating inferred cell-cell signaling across selected ligand-receptor axes. Each plot shows directional connectivity betwen sending and receiving populations based on NICHES scoring. Line thickness represents scaled communication strength; node size reflects the relative abundance of each cell population. **E)** Circuits for Serpina1-Lrp1 and Cd24-Selp axes. **F)** Circuits for II1a-II1r2, II1rn-II1r1, and 111m-II1r2, highlighting inflammatory and regulatory signaling pathways. **G)** Circuits for developmental remodeling signals including Mdk-Tspan1. Efnb2-Ephb3, and Ereg-Erbb3. **H)** Circuits for matrix-associated and EGFR signaling including Fn1-ltgav and Tgfa-Egfr.

To simplify the interpretation of cross-lineage signaling, we first grouped the hillock (basal and luminal) and macrophage (pro-inflammatory and anti-inflammatory) subsets into unified populations, called ‘Hillock_Like’ and ‘Polarized_Mac,’ respectively. We then created a subset object that only included populations of interest for this analysis. These populations included: ‘Hillock_Like,’ ‘Polarized_Mac’, ‘Rspo3+_Mes,’ and “Pdgfrb+_Pericytes’ (**Fig. 6B**). Using unsupervised clustering on the NICHES CellToCell matrix allowed for the grouping of ligand-receptor interactions into discrete signaling modules. We then were able to determine the dominant (highly expressed) sender-receiver pairings within each module which revealed a variation in epithelium-to-immune and mesenchyme-to-epithelium signaling routes that varied across conditions. This variation reveals how lineage-specific signaling shifts in different cellular milieus that possess different cellular ratios.

With knowledge of the presence of distinct epithelial and immune cell states in our three different systems, we hypothesized that context-specific cell states are orchestrated by the differential expression of ligand-receptor pairs mediating cross-compartmental signaling. To test this, we used known ligand-receptor interactions from the FANTOM5 database to identify genes expressed in our dataset and assembled high-confidence sender-receiver pairs (<u>Ramilowski et al., 2015</u>). We then performed differential expression analysis within each cell subset (log_2_ fold-change > 0.25, expressed in ≥25% of cells) and calculated a power metric (expression ratio * avg. log_2_FC). Ranking these interactions by power and adjusted p-value allowed for the identification of signaling interactions that were both differentially expressed and population-specific.

This analysis validated our hypothesis, demonstrating a clear signaling axis between hillock-like cells and activated macrophages. Broadly, this network analysis revealed that all lineages included were communicating with via both autocrine and paracrine signaling (**Fig. 6B**). The ligands *Serpina1*, *Cd24*, *Ill1rn*, and *Il1a* were expressed by both hillock and macrophage populations and targeted receptors on mesenchymal and immune cells (**Fig. 6C-D**). For example, the upregulation of the *Serpina1—Lrp1* and *Cd24—Selp* signaling axes point to the possible role hillock cells play in regulating macrophage adhesion and activation (**Fig. 6E**) (<u>Morrow et al., 2024; Garcia-Arcos, 2022</u>). By comparison, *Il1rn—Il1r1, Il1rn—Il1r2*, and *Il1a—Il1r2* signaling converged on the *Rspo3^+^* mesenchyme node, indicating that epithelial and immune sources may work in tandem to influence mesenchymal behavior (**Fig. 6F**). The frequency of the IL-1 family interactions, as well as their specificity, made them stand out as a potential core organizing principle within the hillock-mesenchyme-macrophage circuit created. The juxtaposition of *Il1a*, a pro-inflammatory cytokine, with *Il1rn*, a natural antagonist that dampens IL-1 signaling suggests not only the delivery of signal, but also a mechanism to balance activation and resolution, almost like a built-in regulatory logic gate (<u>Willart et al., 2012; Eislymayr et al., 2022</u>). Epithelial and myeloid cells do not directly activate mesenchyme engagement in a unidirectional, “fibroinflammatory” response, but instead initiate and modulate mesenchymal engagement to ensure the amplitude and duration of signaling is appropriate for maintaining tissue structure and avoiding excessive remodeling (<u>Minagawa et al., 2014</u>).

We also identified reverse signaling from mesenchymal populations to hillock cells, including *Mdk—Tspan1, Efnb2—Ephb3*, and *Ereg—Erbb3* (**Fig. 6g**) These axes may imply that mesenchymal cells contribute actively to epithelial regulation, delivering cues that support survival, polarity, and lineage plasticity. Midkine (*Mdk*) signaling through *Tspan1* may help maintain epithelial cells in a reparative and plastic state capable of responding to environmental cues when necessary (<u>Zhang et al., 2023</u>). *Tspan1* has been identified as a regulator of epithelial-to-mesenchymal transition (EMT) in alveolar epithelial cells in the context of idiopathic pulmonary fibrosis (IPF) whereby it’s expression downregulates Smad2/3 and beta-catenin signaling (<u>Liu et al., 2019</u>). This activity helps preserve epithelial identity and prevents the transition toward a mesenchymal state (<u>Liu et al., 2019</u>).

*Efnb2—Ephb3* interactions may contribute to the spatial containment of the hillock-like epithelium. Here, *Efnb2*, secreted by *Rspo3*^+^ mesenchyme, acts on the *Ephb3* receptor of the hillock-like population. The upregulation of this signaling axis between these two cell populations likely reinforces compartment boundaries and thus prevents hillock cell expansion into surrounding alveolar or transitional territories (<u>Li et al., 2012; Lewis et al., 2022</u>). This spatial guidance allows hillock-like cells to maintain their identity by preserving their stacked-squamous-like architecture and required spatial insulation. *Ereg—Erbb3*, meanwhile, may promote hillock survival and stress adaptation, consistent with the findings that *Erbb3* protects against injury-induced cell death and facilitates epithelial repair via PI3K/AKT activation (<u>Matsuda et al., 2023; Zhang et al., 2012; Deguchi et al., 2024</u>).

Simultaneously, we observed broader multi-nodal pathways such as the *Tgfa—Egfr* axis, which exhibited robust signaling from hillock cells to multiple compartments, including *Rspo3^+^* mesenchyme, Pdgfrb^+^ pericytes, themselves (**Fig. 6h**). *Tgfa*, a potent *Egfr* ligand involved in epithelial proliferation and morphogenesis under stress (<u>Korfhagen et al., 1994; Hardie et al., 2004; Buckley et al., 2008</u>), was broadly expressed in hillock cells, while *Egfr* was distributed across hillock-like and mesenchymal populations. This specific network implies that Tgfa-producing hillock cells may serve as central broadcasting effector cells to coordinate multicellular responses (sensor cells) via autocrine and paracrine loops. While *Tgfa—Egfr* signaling reinforces epithelial identity, it may also modulate mesenchymal responsiveness or act as a feedback hub. Other cross-compartmental axes, such as *Fn1—Itgav*, flowed predominantly from mesenchyme to epithelium, indicating the support of matrix remodeling and epithelial repair (**Fig. 6h**) (<u>Wang et al., 2022).</u>

Collectively, these patterns describe a multi-lineage signaling circuit that includes epithelial, immune, and mesenchymal populations, each playing a specific, but coordinated role. In this circuit, hillock cells act as central nodes capable of broadcasting signals to modulate immune and mesenchyme driven processes while also responding to instructive cues from themselves. This conclusion is reflected by the existence of shared ligands *Il1a*, *Il1rn*, and *Serpina1*, along with reciprocal mesenchymal cues facilitated via *Mdk*, *Efnb2*, and *Ereg* that suggest a signaling network that is not rigidly regenerative or inflammatory but responsive to tissue context in an adaptive fashion.

Taken together, this pattern of interactions implies a model in which transitional epithelial states are guided by an emergent cross-compartmental logic, one that integrates pro-repair, pro-inflammatory, and regulatory signals across the epithelial, immune, and mesenchymal compartments, with distributed signaling feedback rather than top-down hierarchy, such that tissues can flexibly recalibrate their composition and signaling dynamics in response to perturbation, but not necessarily return to homeostasis. Within this framework, mesenchyme is an active participant that interprets and integrates immune and epithelial inputs to create the structural and signaling environment within which repair occurs.

## Discussion

Here, we report on a lung organoid model defined by compositional inputs that enables multicellular self-organization, lineage-specific fate emergence, and microenvironmental signaling analysis. By varying the relative contribution of immune- and mesenchyme-containing inputs under uniform matrix, media, and environmental conditions, we established a platform to investigate how initial cellular composition shapes epithelial and immune dynamics. The ability to maintain cell-type diversity and track context-dependent behavior under standardized conditions enables a degree of experimental control rarely achievable in vivo and allows for interrogation of regenerative signaling at both cellular and network levels.

While recent work has used fluorescence-activated cell sorting (FACS) to expand BAL-derived progenitors (<u>Liu et al., 2024</u>), our BAL_3D organoid model exhibits substantial epithelial expansion from rat BAL samples that had initially comprised only a minor population of epithelium and no mesenchyme. These cultures, which were predominantly immune in composition and devoid of any deliberate injury cues or progenitor enrichment, reliably produced a wide range of epithelial cell types, including the transitional and secretory subtypes of increasing interest to the pulmonary biology community. Additionally, this system, as well as the PD_3D and Mixed_3D systems, allowed for the expansion of both proximal and distal populations without requiring specialized “proximal” or “distal” growth media.

The ability to generate complex epithelial structures from a cell suspension with trace epithelial input under defined culture conditions recognizes our BAL_3D system as having significant translational potential for regenerative medicine and cell therapy development.

Transitional epithelial states (i.e., *Sox9⁺* stressed progenitors, hillock cells, and secretory-alveolar intermediates) emerged without deliberate injury cues such as exposure to noxious agents or mechanical agitation, suggesting that matrix and media alone promote epithelial plasticity. Nevertheless, we do recognize that the culture medium utilized was originally designed to expand embryonic lung progenitors (<u>Nichane et al., 2017</u>). Despite this, the relative abundance and character of transitional populations in our systems varied between conditions, indicating that local niche composition, especially regarding immune and mesenchymal cells, can influence epithelial remodeling. Among the transitional states observed, hillock-like cells expanded with striking consistency demonstrating the utility of this system to expand these cells in vivo, thus affording researchers the ability probe their molecular identity, functional role, and potential contributions to tissue homeostasis in greater depth.

Furthermore, hillock-like cells were most prominent in conditions retaining mesenchymal populations, particularly Mixed_3D and PD_3D. Their consistent emergence under defined conditions provides a model to study their role in epithelial repair, immune coordination, and pathologic remodeling. These *Krt13⁺/Sprr1a⁺* cells exhibited stratified organization, context-specific polarization, and expression of immune-modulatory genes, features consistent with barrier-associated remodeling (<u>Holztmann et al., 2002; Stouch et al., 2016; Zhang et al., 2023</u>). Luminal hillock cells expressed cytokines, alarmins, and antimicrobial genes, suggesting a regulatory role at the interface of epithelial and immune compartments. To our knowledge, this is the first study to begin delineating the cell-cell signaling architecture associated with hillock cells. We identified signaling axes linking hillock cells to mesenchymal and immune partners, including *Il1a—Il1r2 and Serpina1—Lrp1*, that position these cells as both sensors and broadcasters of inflammatory and regenerative cues. Rather than passive responders or indicators of dysfunction, hillock cells may serve as active modulators of tissue recovery, helping to calibrate local inflammation and stabilize epithelial integrity (<u>Lin et al., 2024</u>).

As with the more differentiated luminal hillock-like cells generated by our models, the RAS-like intermediates expanded most robustly in systems containing mesenchyme (e.g., Mixed_3D and PD_3D). Therefore, we propose that the presence of mesenchyme, and subsequent presence of mesenchymal-derived ligands like WNT and BMP critical for epithelial patterning and resolution are supportive of transitional epithelial states, providing them with stability to exist at the “ridges” of Waddington’s landscape (<u>Oikonomakos et al., 2024; Frum et al., 2023; Konigshoff & Eickelberg, 2009; Ostrin et al., 2018</u> ). This observation is consistent with the growing body of evidence that epithelial fate is not strictly lineage-encoded but rather dynamically tuned by environmental signals (<u>Hurley et al., 2020; Toth et al., 2023</u>). Additionally, it lends further credence to the hypothesis that maladaptive stabilization of transitional states underlies disease pathogenesis (<u>Jones et al., 2024; Strunz et al., 2020</u>). Our model provides a tractable system to study the regulation and persistence of these alveolar-secretory intermediates, just as it does for the hillock-like cells.

Like the polarization displayed by several epithelial intermediates in our systems and their implications regarding plasticity in response to local cues, the macrophages that were sustained in each 3D system can be thought of in a similar way. The pro-versus anti-inflammatory bias exhibited by the activated macrophages aligns with frameworks that poise macrophage plasticity as environmentally regulated rather than preprogrammed, especially in the context of chronic lung diseases (<u>Eapen et al., 2017; Ge et al., 2024</u>). Macrophages that were maintained in our 3D culture platform in the presence of mesenchymal cues (Mixed_3D and PD_3D) were biased toward a pro-inflammatory state and thus, upregulated canonical inflammatory mediators such as *Il1a*, *Il1b*, *Cxcl1/2/3*, and *Nos2*. These findings are in line with previous reports of mesenchyme guiding a pro-inflammatory state in development and injury processes (<u>Eislmayr et al., 2022; Wang et al., 2022; Giri et al., 2020</u>). When mesenchyme was not present to provide secreted ligands such as *Il6*, *Ccl2* and *Cxcl12* that have been demonstrated to activate macrophages cells to a pro-inflammatory state (<u>Li et al., 2021; Liang et al., 2012; Ayaub et al., 2017</u>), as was the case in the BAL_3D system, macrophages adopted an anti-inflammatory phenotype, amplifying programs associated with immune resolution and epithelial support via the expression of genes such as Pparg, *Cela1*, and *Lpl* (<u>Lee et al., 2024; Babaev et al., 1999; Joshi et al., 2017</u>). While our findings are unable to confidently and accurately define the exact molecular mechanisms driving the shifts observed in macrophage polarization in chronic lung disease, our findings do suggest that the crosstalk driving macrophage polarization is not mediated by a singular axis between mesenchyme and immune or epithelium and immune. Rather, the three lineages (epithelium, immune, and mesenchyme) are in dynamic conversation with one another to define regenerative versus maladaptive trajectories.

Across the various cell types described and our study of their intra-lineage communication, we continuously come back to the framework set forth by <u>Medzhitov (2021)</u> whereby inflammation is not considered a byproduct of damage but a regulatory process that maintains tissue function through adaptive coordination. From this perspective, the emergence of *Sox9^+^* stressed progenitors, hillock-like cells, and RAS-like intermediates ought to be thought of as an anticipatory epithelial response to environmental cues to ensure regenerative preparedness. These transitional states are far from artifacts of culture but may serve as context-sensitive intermediates ready to modulate proliferation, stress response, and signaling plasticity (<u>Warren et al., 2024</u>). The reproducibility of these intermediates, which are of increasing interest to the lung biology community, speaks to the value of our platform in studying successful regeneration as well as its derailment in chronic disease.

### Limitations

Although our organoid model provides a functional and convenient system for exploring epithelial plasticity and intercellular signaling under controlled conditions, several limitations warrant consideration.

First, while we performed single-cell RNA sequencing on the initial PD and BAL isolates that were seeded in various ratios for our three systems, we did not prospectively enrich, deplete, or barcode individual lineages before the start of culture. Therefore, while we were able to characterize the lineages and their respective cell types present at the start of culture in each system, we cannot reliably determine which exact cells differentiated in culture. That being said, our approach did allow us to identify which cell types, more broadly, persisted and which ones emerged spontaneously in 3D culture. If a particular cell type with a specific transcriptional program was not present in either of our starting samples (PD or BAL), but was present in the sequencing data generated from our engineered samples (PD_3D, BAL_3D, and Mixed_3D), it is likely that these transitional cell types emerged as a product of the system in which their origin cell was grown in, highlighting the plasticity and context-dependent emergence of these intermediates. Future experiments employing prospective lineage tracing, genetic barcoding, or FACS-based enrichment protocols would serve useful to resolve the origin and fate of particular cell types within the proposed regenerative circuit.

Second, our analysis was conducted at a single, terminal time point (day 10). Without sampling over time, it remains unclear whether the epithelial populations we observed such as the stressed Sox9+ progenitors, hillock cells, and secretory-alveolar intermediates are stable endpoints of transient endpoints along the path to terminal differentiation. Defining such snapshots in time for these transitional cells warrants time-course transcriptomic profiling in combination with the already utilized trajectory inference and fate-mapping tools.

Third, although we confirmed the presence and localization of the various highlighted epithelial populations of interest via immunohistochemistry, we were not able to effectively validate the spatial relationships between all signaling partners (e.g., macrophage, mesenchymal, hillock cells). These reasoning behind these limitations are two-fold: first, we found that our organoids exhibited such significant structural complexity that when it came time to dissociating them (i.e., digesting the Matrigel), they had remodeled the culture environment to such a large degree that dissociation was quite difficult. Additionally, organoids in the BAL_3D system, specifically, frequently lost structural integrity quickly upon dissociation making them difficult to manipulate for whole-mount staining. Second, there remains to be a limited availability of high-quality rat-reactive antibodies for the detection of the specific molecules of interest in our systems. Future application of spatial transcriptomics or high-resolution in situ hybridization approaches will be crucial to determine whether observed ligand-receptor circuits correspond to physically adjacent cell types and to define how niche topology may constrain or facilitate regenerative signaling.

Fourth, although media conditions were standardized across cultures, the inclusion of small molecule inhibitors and growth factors inherently selects for progenitor maintenance. While this design enabled controlled comparisons across compositions, it may have artificially sustained transitional states that would otherwise resolve under more restrictive conditions. Future iterative perturbation of individual signaling components will be important to define the context-dependence of observed plasticity.

Finally, this model captures many aspects of regenerative signaling and tissue self-organization, but it does not incorporate biomechanical inputs (e.g., stretch, airflow, perfusion), which are known to affect epithelial differentiation, cytoskeletal dynamics, and extracellular matrix remodeling in vivo. For example, cyclic stretch and airflow have been shown to regulate alveolar type I/II fate balance, in addition to modulating surfactant production and progenitor cell activation in the native lung (<u>Sanchez-Esteban, 2001; Goodwin et al., 2023</u>). Mechanical strain has also been implicated in the mediation of mesenchymal-epithelial interactions during branching morphogenesis (<u>Ahmed et al., 2023; Shiraishi et al., 2023; Vining & Mooney, 2017</u>) and reported to bias immune cells toward polarized states via mechanotransduction pathways such as YAP/TAZ and PI3K/AKT/mTOR (<u>Meli et al., 2020; Sapudom et al., 2024; Di-Luoffo et al., 2021</u>). Incorporating biomechanical forces into future versions of this model, whether via microfluidic perfusion, cyclic stretch chambers, or bioreactors, may enhance physiological fidelity and yield new insight into how physical stimuli intersect with lineage composition and intercellular communication.

## Future Directions

Our findings suggest that transitional cells are part of a larger regenerative logic: plastic intermediates that integrate local cues to orchestrate repair, calibrate inflammation, and potentially resist progression to disease.

Based on this work, we propose several questions for future study:

- What determines whether transitional states resolve or persist? Can Sox9⁺ or hillock-like cells revert, differentiate, or become pathologically stabilized depending on microenvironmental context?
- Is the hillock niche an inducible, facultative module for immune-epithelial coordination in the proximal lung or a universal regenerative structure across airway compartments? How does its formation relate to chronic remodeling in disease?
- How tightly couples are macrophage polarization and epithelial transitions? What additional signaling axes beyond the IL-1 family and Serpina1 axes may govern their coupling or uncoupling?
- Can lineage manipulation at the start of culture (i.e., day 0) redirect the subsequent regenerative trajectories we observed, and how do immune or mesenchymal biases (imposed at seeding) influence the fate spectrum or propensity for repair versus maladaptive remodeling?
- What role do biomechanical forces play in driving or stabilizing transitional states? As epithelial fate decisions may remain plastic during static culture conditions, it is possible that mechanical inputs such as cyclic strain or shear stress could guide the trajectory of transitional cell states in culture. Incorporating these elements into future models might help us illuminate how tissue mechanics intersect with regenerative patterning.

We present these experiments and findings to the community to advance understanding of how compositional and signaling cues guide epithelial state transitions. This knowledge may open new therapeutic avenues for restoring tissue integrity in chronic lung disease.

## Materials & Methods

### Animal Handling and Lung Dissociation

8-10-week-old male Sprague Dawley rats (n = 6; 250 g ± 25) were sacrificed for cell isolation. Rats were sedated in an induction chamber containing Isoflurane-soaked (20% w/v) gauze, followed by intraperitoneal administration of 0.25 mL Ketamine-Xylazine solution (K: 75 mg/mL; X: 5 mg/mL) and 0.15 mL heparin (1,000 U/mL). Once fully sedated, animals were sterilized with 70% ethanol and povidone-iodine prep pads (Dynarex). Through the thoracic cavity, the lungs were allowed to deflate, and the thymus was removed. The clavicle was dissected to expose the trachea, which was cannulated using a barbed Y 1/16” connector. A second barbed Y 1/16” connector was used to cannulate the pulmonary artery. The cardiac apex was excised to allow fluid flow, and the lung tissue was perfused at 50 mL/min with heparin (100 U/mL) and sodium nitroprusside (SNP; 0.1 mg/mL). The lungs were then excised and placed in a petri dish for perfusion with dissociation buffer (DMEM HG (Gibco), 1 mg/mL Collagenase/Dispase (Roche), 3 U/mL Elastase (Worthington), 20 U/mL DNase) through both the airway and vasculature via gravity. 10 mL of dissociation buffer was used to inflate the lungs three times. The lungs were transferred to conical tubes containing dissociation buffer and incubated on a rocker at 37°C for 25 minutes. All animal procedures were conducted in accordance with Yale Institutional Animal Care and Use Committee (IACUC).

### Pulmonary Dissociation Cell Isolation and Processing

Cells were dissociated from native rat lung tissue following enzymatic digestion (described above) and then mechanically disrupted using a spatula using a protocol previously reported by our lab (<u>Leiby et al., 2023; Greaney et al., 2020; Raredon et al., 2019</u>). Large collagenous structures were allowed to pass through a strainer. The strainer was then washed with 20 mL of quenching medium (DMEM HG (Gibco, 11965092), 10% FBS, 1% P/S (Worthington), 0.1% gentamycin (Worthington), 1% ampicillin (Worthington)), and the suspension was collected. The suspension was centrifuged at 300 x g for 5 minutes at 4°C to pellet cells and supernatant was then discarded. The pellet was resuspended in an equal volume of ACK Lysing Buffer (Gibco, A1049201) and gently agitated by tapping for 2 minutes at room temperature to lyse red blood cells. This suspension was then diluted with 10 mL of MACS buffer (0.1% BSA (Gemini) in PBS (Gibco), filtered), and then centrifugation at 300 x g for 5 minutes at 4°C to pellet. Supernatant was aspirated and the pellet resuspended in 5 mL of MACS buffer before being passed through a 70 μm strainer (Falcon, 352350) and then a 40 μm strainer (Falcon, 352340). The filtered cell suspension was then centrifuged again at 300 x g for 5 minutes at 4°C to pellet cells, supernatant was aspirated, and the pellet resuspend in 5 mL of MACS buffer. To ensure a single-cell suspension, the suspension was passed through a 40 μm strainer twice more. Cells were then counted and viability evaluated using 0.4% Trypan Blue solution (Gibco, 15250061). Cells were then resuspended in MACS buffer at 1 mL per 10⁷ total cells before proceeding to cell selection using *Dclk1^+^* tagged beads.

Following enzymatic digestion, cells were dissociated from the lung tissue using a spatula and passed through a strainer, leaving behind larger collagenous structures. The strainer was rinsed with 20 mL of quenching medium (DMEM HG (Gibco, 11965092), 10% FBS, 1% P/S (Worthington), 0.1% gentamycin (Worthington), 1% ampicillin (Worthington)), and the resulting cell suspension was collected. The suspension was centrifuged at 300 x g for 5 minutes at 4°C, and the supernatant was discarded. The cell pellet was resuspended in an equal volume of ACK Lysing Buffer (Gibco, A1049201) and agitated manually with gentle tapping for 2 minutes at room temperature to lyse red blood cells. The cell suspension was then diluted with 10 mL of MACS buffer (0.1% BSA (Gemini) in PBS (Gibco), filtered), followed by centrifugation at 300 x g for 5 minutes at 4°C. Supernatant was aspirated, and the pellet was resuspended in 5 mL of MACS buffer. The suspension was sequentially filtered through 70 μm and 40 μm strainers (Falcon, 352350, 352340) to remove debris. Cells were pelleted by centrifugation at 300 x g for 5 minutes at 4°C, and the supernatant was removed. The pellet was resuspended in 5 mL of MACS buffer, filtered twice through a 40 μm strainer, and counted, as well as evaluated for viability using 0.4% Trypan Blue solution (Gibco, 15250061). Cells were resuspended in the 1 mL of MACS buffer per 10⁷ total cells before proceeding to cell selection with *Dclk1*^+^ tagged beads.

#### Preparation of Anti-Dclk1-Tagged Magnetic Beads

Magnetic beads were prepared using 1 mg (100 μL) of M-280 Sheep anti-Rabbit IgG Dynabeads (ThermoFisher, 11203D) suspended in 500 μL of MACS buffer (0.1% BSA (GEMINI, 700-100P) in PBS (Gibco, 14190144)). Bead suspension was placed in a DynaMag™-5 Magnet (ThermoFisher, 12303D) for 1 minute to separate the beads, and the supernatant was aspirated. Beads were washed three times with 500 μL of MACS at each wash. After the final wash, Dynabeads were resuspended in MACS buffer, and 10 μg of anti-*Dclk1* antibody (Abcam, ab31704) was added to achieve a final volume of 100 μL. The volume of the antibody solution was adjusted based on the supplied concentration. The bead-antibody mixture was incubated at room temperature for 30 minutes with gentle tilting and rotation on a standard platform shaker (VWR). Following incubation, 500 μL of MACS buffer was added, and the suspension was pipetted up and down several times. The beads were washed three additional times with MACS buffer using the DynaMag™-5 Magnet. The final bead suspension was resuspended in 100 μL of MACS buffer and stored at 4°C for no more than 24 hours before use.

#### Cell Type Enrichment with Dclk1^+^ Tagged Beads

Cells dissociated from lungs were strained through a 40 μm cell strainer (Falcon, 352340) to break up clumps of cells, and then incubated at 4°C with Dynabeads prepared with anti-*Dclk1* antibody (Abcam, ab31704) (25 µL per 10⁷ total cells) for 30 minutes with tilting and rotation using a tube tilter to prevent bead settling. The suspension was diluted two-fold with MACS buffer to reduce cell adhesion, and then the tubes were placed in the DynaMag™-5 Magnet under aseptic conditions for 2 minutes and the supernatant was aspirated. Cells were washed with 1 mL of MACS buffer and the magnet was used again for an additional 1 min and this step was repeated four times. The final cell pellet was resuspended in MACS buffer to remove any cells that had stuck to the tube walls, centrifuged at 300 g for 5 minutes at 4°C, supernatant removed, and cells were resuspended in LPM-3D medium to achieve a concentration of organoid plating.

### Bronchoalveolar Lavage (BAL) Cell Collection

BAL-derived (immune and epithelium) cells were harvested from 8-10-week-old male Sprague Dawley rats (n = 3; 250 g ± 25), as described above in *Animal Handling and Lung Dissociation* methods. Lungs were placed into a conical tube with 40 mL of ice-cold saline solution (PBS, 1% P/S) on ice for 60 minutes. The lungs were then maximally inflated by perfusing the trachea with sterile saline solution, gently massaging the tissue. Negative pressure was applied at the open end of the tracheal cannula by covering it with one finger and the PBS containing cells was aspirated using a 10 mL syringe. This process was repeated 5-7 times, with all the aspirated fluid being collected into a sterile 50 mL conical tube. The BAL cells were centrifuged (500 x g for 10 minutes at 4°C), supernatant aspirated, and resuspended in an equal volume of ACK lysing buffer (Gibco, A1049201) (1:1 ratio). The pellet was then agitated manually with gentle tapping for 2 minutes at room temperature before dilution with 2 mL DMEM supplemented with 10% FBS. The cells were counted and resuspended to the desired concentration for organoid seeding.

### Organoid Culture

Organoid systems were generated in 0.4 μm, 24-well cell culture inserts (Falcon) using a 1:1 (v/v) ratio of growth factor-reduced Matrigel (Corning, Matrigel GFR, Phenol Red-Free, #356231) to cell suspension. Each transwell insert contained a total volume of 90 μL, consisting of 45 μL Matrigel and 45 μL cell suspension. All organoid suspensions (Matrigel + cells) were prepared in LPM-3D medium and maintained at 37°C. The following cell seeding densities were used: PD_3D (0.1 × 10⁶ cells from PD), Mixed_3D (5 × 10⁴ PD cells and 5 × 10⁴ BAL cells), and BAL_3D (0.1 × 10⁶ cells from BAL).

Organoid suspensions (Matrigel + cell suspension) were prepared on dry ice, and chilled. Wide-bore pipette tips were used for sample distribution to ensure consistent handling and to avoid further physical agitation of the cells. Large bubbles in the Matrigel were removed using 27 G x ½ (0.4 mm x 13 mm; BD, 305109) sterile needles before Matrigel was allowed to solidify. Following a 30-minute incubation at 37°C, 500 μL of culture medium was added to the basolateral compartment of the Transwell system. Cell culture medium was changed every 48 hours across all conditions and media samples were saved for potential future experiments. Biological replicates were prepared for each condition (n=18, with n=6 per experiment). Organoids were maintained in culture for 10 days at 5% CO_2_ at 37°C. Organoids sent for single-cell RNA sequencing (scRNAseq) were dissociated on day 10 following the same methods outlined in methods section *Organoid Dissociation for scRNAseq and Other Downstream Processes*.

### Organoid and Tissue Fixation

#### Native Rat Lung

Native rat lungs fixed via perfusion with 4% paraformaldehyde (PFA) at a flow rate of 60 mL/min for a minimum of 12 hours. After fixation, the lungs were transferred to a sterile petri dish for dissection. Anatomical dissection was performed to ensure that each of the five lobes and the trachea were embedded individually. Trachea and lobes were sectioned longitudinally. Tissue embedding and sectioning were performed by Yale Pathology Tissue Services (YPTS).

#### Organoids

Organoids were fixed using one of three methods, depending on the intended downstream application. For organoids embedded and sectioned with the Transwell membrane, cell culture media was removed, and 0.5 mL of 4% PFA was added to the basolateral compartment of the Transwell insert. The plate was placed on a rocker for 16 hours without prior Matrigel digestion. After 12 hours of incubation, the Transwell membrane containing the Matrigel-embedded organoids was excised using a scalpel. The excised membrane was embedded in HistoGel (ThermoFisher, HG-4000-012) before paraffin embedding and sectioning by Yale Pathology Tissue Services (YPTS).

For organoids embedded in paraffin without Matrigel, samples were washed with ice-cold PBS and placed on an orbital shaker at 4°C for 60 minutes to digest as much of the ECM as possible without the use of enzymes. Organoids were collected in PBS using a P1000 wide-bore pipette coated with BSA to prevent adherence to the pipette tip. The collected organoids were centrifuged in a 15 mL conical tube coated with BSA at 200 x g for 3 minutes at 4°C. The supernatant was carefully aspirated to avoid loss of organoids, and the pellet was resuspended in 2 mL of 4% PFA. Organoids in 4% PFA were then placed on a rocker for a minimum of 4 hours at room temperature. Following fixation, PFA was aspirated, and the organoids were embedded in a 100-200 μL droplet of HistoGel (ThermoFisher, HG-4000-012) before paraffin embedding and sectioning by Yale Pathology Tissue Services (YPTS).

For organoids undergoing whole-mount staining, the same preparation steps as those used for organoids embedded without Matrigel were followed. However, instead of HistoGel embedding, organoids were resuspended in PBS after fixation with 4% PFA and transferred to 4-well chamber slides (ThermoFisher, Nunc Lab-Tek II) coated with 0.4% BSA in PBS, to prevent adherence. Staining and imaging was then carried out with the organoids in the chamber slides. For information on whole-mount staining, see *Methods: Immunohistochemistry of Whole-Mount Organoids*.

### LPM-3D Media Preparation

Epithelial expansion medium was prepared as previously described, with slight modifications to make the media appropriate for rat-derived cells (<u>Nichane et al., 2017</u>). Briefly, Advanced DMEM/F12 basal medium (ThermoFisher, 11320033) was filtered and supplemented with the following rat growth factors: FGF10 (50 ng/mL; R&D Systems, 7804-FG), FGF9 (50 ng/mL; R&D Systems, 273-F9), and EGF (50 ng/mL; R&D Systems, 3214-EG) (**Supplementary Table 1 (S1)**). Additionally, the medium was supplemented with small molecule inhibitors, including CHIR99021 (3 μM; GSK-3 inhibitor/Wnt activator; Cayman Chemical, 13122), BIRB796 (1 μM; p38-MAPK inhibitor; Cayman Chemical), Y27632 (10 μM; ROCK inhibitor; Cayman Chemical, 10460), and A8301 (1 μM; Activin/NODAL/TGF-β pathway inhibitor; Cayman Chemical, 9001799).

Further supplements included heparin (5 μg/mL; Sigma Aldrich, 9041-08-1), insulin (10 μg/mL; Roche), and transferrin (15 μg/mL; Roche). The prepared medium was pre-warmed to 37°C before given to organoid systems at feeding. Antibiotics were added to prevent at concentrations of 1% penicillin-streptomycin (P/S) and 0.1% gentamicin to prevent infection.

### Immunohistochemistry of Paraffin Embedded Sections

Paraffin-embedded sections of perfused native rat lung, as well as trachea and organoid systems were immunostained to better understand protein expression localization patterns. Samples were first deparaffinized by heating at 65°C for 30 minutes, followed by sequential rinsing in a xylene and ethanol gradient (100%, 95%, and 70% ethanol).

Antigen retrieval was performed by immersing the slides in antigen retrieval buffer (0.1 M citric acid, 0.05% Tween 20, pH 6) and placing them in a water bath at 75°C for 20 minutes. Slides were then cooled to room temperature for a minimum of 15 minutes.

Following antigen retrieval, slides were placed in a slide box containing PBS for 5 minutes. Tissue sections on the slides were then outlined using a hydrophobic marker (ReadyProbes™ Hydrophobic Barrier Pap Pen; ThermoFisher). Tissue sections were then rinsed 3-5 times with PBS. Two different permeabilization buffers were employed depending on the proteins being stained for. For nuclear stains (e.g. transcription factors such as *Sox9*), tissue sections were permeabilized using PBS supplemented with 0.2% Triton-X (Invitrogen, HFH10) for 15 minutes. After permeabilization, samples were blocked with blocking buffer (0.75% glycine, 5% Donkey Serum (Gibco, PCN5000) in PBS) for 1 hour to minimize nonspecific binding. For cell surface and cytoplasmic markers, tissue sections were permeabilized with 0.5% Tween-20 (Millipore, P1379) in PBS for 15 minutes, followed by the same blocking buffer step described previously.

Primary antibodies, prepared in the same blocking buffer, were applied to the samples at their specified concentrations (*See Supplemental Table 2 (S2)*) and incubated overnight at 4°C. Following incubation, slides were washed 3-5 times with PBS before the application of secondary antibodies (concentration: 1:500) for 1 hour at room temperature. Slides were washed again with PBS and counterstained with DAPI (ThermoFisher, 62247) (1:1000 in PBS) for 1 minute to visualize nuclei. Finally, sections were mounted using PVA-DABCO (Sigma Aldrich,10981). Details for all primary and secondary antibodies used are provided in **Supplementary Tables 2 (S2) and 3 (S3),** respectively. All samples were imaged using either the EVOS Auto FL 2 imaging system or Stellaris 8 confocal microscope.

### Immunohistochemistry of Whole-Mount Organoids

Organoids digested from Matrigel, as described above (*See Methods: Organoid and Tissue Fixation (Organoids)*), were transferred to a 4-well chamber slide (ThermoFisher, Nunc Lab-Tek II) coated with 0.4% BSA in PBS (to prevent adherence). Multi-well chamber slides allowed for the clear separation of condition-specific organoids. Roughly 8-12 organoids, depending on size, were transferred to each chamber using a P1000 wide-bore pipette tip coated with 0.4% BSA in PBS. The same protein-dependent permeabilization steps were performed as previously described (*See Methods; Immunohistochemistry of Paraffin Embedded Sections*). Organoids were then blocked with 500 μL of blocking buffer (0.75% glycine, 5% Donkey Serum (Gibco, PCN5000) in PBS) for 2 hours at 37°C and placed on an orbital shaker rotating at 280 rpm. Blocking buffer was then carefully removed, ensuring not to disrupt organoids, and blocking buffer containing primary antibodies (at respective concentrations) were added to the chambers. Organoids with primary antibodies were incubated overnight at 4°C with gentle shaking on an orbital shaker (280 rpm). The next day, the primary antibody solution was removed carefully, and the samples were washed with 500 μL of PBS 3 times over, with a 10-minute incubation period between each wash. The samples were then incubated with 500 μL of secondary antibody (each at a concentration of 1:500) in blocking buffer for 2 hours at 37°C, with gentle rocking using the orbital shaker.

Secondary antibody was then carefully aspirated, and samples were incubated with DAPI (ThermoFisher, 62247) solution (1:1000 in PBS) for 1 minute at 37°C. DAPI solution was then removed, and 250 μL of PVA-DABCO (Sigma Aldrich,10981) was added to each chamber before imaging on either the EVOS Auto FL 2 imaging system or Stellaris 8 confocal microscope.

### Immunofluorescence of Starting Population Cytospins

Cytospins were generated for all three starting populations (BAL, PD, and a 1:1 mix of the two that subsequently became Mixed_3D). Slides were securely placed in the slide holder, ensuring proper orientation with the frosted side of the slide facing upwards. A filter card, with its absorbent side against the slide, was positioned alongside the cytofunnel, ensuring proper alignment of all components before securing the holder.

Each assembled holder was then placed in the Cytospin centrifuge (Cytospin 4 Centrifuge, ThermoScientific). Each cell suspension, prepared at a concentration of 0.5 × 10⁶ cells/mL in DMEM + 10% FBS, was pipetted into each cytofunnel as a 200 µL volume. The slides were centrifuged at 1000 × g for 5 minutes. Following centrifugation, slides were removed from the centrifuge and taken out of the holders. Slides were placed cell-side up and left to air-dry for 1 hour. Spots containing cell suspensions were outlined with a hydrophobic marker (ReadyProbes™ Hydrophobic Barrier Pap Pen; ThermoFisher) before being fixed with 4% paraformaldehyde in PBS for 10 minutes at room temperature. After fixation, slides were rinsed 3 times with PBS and stored in PBS at 4°C until needed for staining (up to one week). Slides were processed as described in the previous section from the antigen retrieval step onward (*See Methods: Immunohistochemistry of Paraffin Embedded Sections*). While BAL starting populations were stained and imaged, PD (for PD_3D) and BAL+PD (for Mixed_3D) were stained, but did not yield valuable data as the beads used for selection made it difficult to visualize cells.

### Organoid Image Analysis & Quantification

Time course brightfield images of each condition (n=3) were taken at days 0, 3, 5, 7, 9, and 10 using the EVOS FL Auto 2 microscope. Stitched images of each well were generated. After image acquisition, images were analyzed using QUPath (v.0.5.1).

Organoid areas in stitched images at each time point were recorded using both the ellipse and brush tools in QUPath. To facilitate accurate organoid area quantification (µm^2^), image scale was calibrated in QuPath using the embedded 1 mm scale bar present in each stitched image, which was measured by counting pixels spanning the scale bar using Fiji (ImageJ) and entered manually into QUPath image properties to ensure that all organoid area measurements reflected physical dimensions. Organoid count and area values were averaged across replicates per condition and timepoint. Time was converted to hours and line plots generated showing mean values ± standard deviation in R. Welch’s t-tests performed comparing conditions at each timepoint for both organoid count and area without assuming equal variance. Resulting p-values, adjusted using the Benjamini-Hochberg (BH) method to control the false discovery rate, are reported as follows: p ≤ 0.05 (*), ≤ 0.01 (**), ≤ 0.001(***).

### Organoid Dissociation for scRNAseq and Other Downstream Processes

Organoids were dissociated from Matrigel ECM using an adapted enzymatic digestion protocol previously described above (*Methods: Pulmonary Dissociation Cell Isolation and Processing*). The culture medium was aspirated, and 1 mL of ice-cold DPBS (Gibco, 14190144) was added to each well; the growth factor-reduced Matrigel (Corning, Matrigel GFR, Phenol Red-Free, #356231) matrix was disrupted by gently pipetting the suspension up and down approximately 20 times using a P1000 wide-bore tip coated with 0.1% BSA in PBS to prevent organoids from adhering to the walls of the pipette tip. The organoid suspension was transferred into a 15 mL conical tube pre-coated with anti-adherence solution or 0.1% BSA in PBS and centrifuged at 300 x g for 5 minutes at 4°C to pellet the organoid-cell suspension. Supernatant was aspirated, and the pellet washed with 0.5 mL of ice-cold PBS to remove as much of the residual Matrigel as possible before enzymatic dissociation. The pelleted organoids were resuspended in 1 mL of dissociation enzyme solution (DMEM HG (Gibco, 11965092), 1 mg/mL Collagenase/Dispase (Roche, 11097113001), 3 U/mL Elastase (Worthington, LS002292), and 20 U/mL DNase) and incubated at 37°C on an orbital shaker to provide gentle agitation and facilitate dissociation while minimizing mechanical stress. After 20-25 minutes of incubation, the suspension was gently pipetted up and down 5-10 times using a wide-bore tip before proceeding to filtration.

To terminate enzymatic activity, 10 mL of Advanced DMEM/F12 medium (ThermoFisher, 12634010) was added to the cell suspension-enzyme mixture. The cells were then filtered through a 100 μm cell strainer (Corning, 431752) and the rubber head of a 3 mL syringe plunger (BD, 309657) was used to gently push any remaining cell aggregates through the strainer. The suspension was again centrifuged at 300 x g for 5 minutes at 4°C, and the supernatant was discarded. The pellet was again resuspended in 5 mL of LPM-3D medium and filtered through a 70 μm cell strainer (Corning, 431751), using a clean syringe plunger to push remaining aggregates through. The filtered suspension was centrifuged at 300 x g for 5 minutes at 4°C one last time before being filtered through a 40 μm cell strainer (Corning, 431750) to ensure a single-cell suspension. Cells were then counted using a hemocytometer to determine concentration and viability was assessed using 0.4% Trypan Blue solution (Gibco, 15250061) before being processed for downstream applications—either scRNAseq or passaging or cryopreservation. Samples dissociated for single-cell RNA sequencing were resuspended at 1 million cells/mL in 0.1% BSA in PBS. Cells to be cryopreserved were resuspended in the desired volume of freezing medium (90% FBS (Hyclone, SH30071), 10% DMSO (Sigma, C6295)) before being transferred to -80 °C for no more than 24 hours and then the cryogenic dewar for longer-term storage.

### Single-Cell RNA Sequencing Library Preparation, Sequencing, and Alignment

Dissociated organoid and starting population cell suspensions were prepared for single-cell RNA sequencing using the Chromium Next GEM Single Cell 3’ Reagent Kits v3.1, according to the manufacturer’s instructions (10x Genomics, Pleasanton, CA). After dissociation, cells were counted, assessed for viability using 0.4% Trypan Blue solution, and then resuspended at a concentration of 1 million cells/mL in 0.1% BSA in PBS for a targeted cell recover of 2,000-10,000 cells per sample. Libraries were sequenced by the Yale Center for Genomic Analysis (YCGA) at a target depth of 50,000 reads per cell using the Illumina NovaSeq 6000 platform. Alignment was performed using Cell Ranger (v8.0.1) (10 Genomics) and the reference transcriptome Rattus norvegicus.Rnor_6.0-95. Alignment was performed with the “-include - introns” option enabled, allowing both exonic and intronic reads to be considered for gene expression quantification.

### Single-Cell Data Processing and Analysis

Single-cell RNA sequencing (scRNA-seq) data were processed individually for each sample using the Seurat package (v5.2.0), following best practices (<u>Luecken & Theis, 2019</u>). Raw sequencing data files were read into R using the Read10X function, and Seurat objects were created for each sample with a minimum of 3 cells and 50 features. Quality control metrics were calculated, including the percentage of mitochondrial gene content (percent.mt), total unique molecular identifier (UMI) counts (nCount_RNA), and detected features (nFeature_RNA). Sample-specific filtering thresholds for nCount_RNA and nFeature_RNA were applied as detailed in **Supplemental Table 4 (S4).**

Initial quality control (QC) was performed to remove low-quality cells based on mitochondrial gene expression and unique feature counts. Cells with high mitochondrial percentages or low feature counts were excluded from further analysis. To ensure accurate identification and removal of low-information cells, an initial clustering step was performed at a high resolution (e.g., res = 5.0) to finely delineate clusters. This allowed us to identify and remove clusters with low gene expression variability or high proportions of ambient RNA contamination. Once low-information cells were filtered out, we re-clustered the remaining cells at an optimized resolution to obtain biologically meaningful clusters. QC plots for the starting BAL population can be found in **Supplemental Fig. 5**. QC plots for the starting PD population can be found in **Supplemental Fig. 6**. QC plots for the organoid sequencing data (merged) can be found in **Supplemental Fig. 4**.

Dimensionality reduction was conducted using principal component analysis (PCA), followed by Uniform Manifold Approximation and Projection (UMAP) for visualization. Clustering was performed using the Louvain algorithm implemented in Seurat. UMAP dimensionality reduction was performed using RunUMAP with the selected principal components, followed by construction of a shared nearest neighbor (SNN) graph using FindNeighbors(). Clustering was conducted using the FindClusters() function, a Louvain algorithm implemented in Seurat, with a range of resolution values to achieve optimal clustering granularity.

#### Differential Expression Analysis

To identify marker genes for each cluster, differential expression analysis was performed using FindAllMarkers() with a minimum percentage threshold of 0.1 and log-fold change threshold of 0.1. Top markers were selected based on power metrics (ratio * log-fold change), and heatmaps, as well as volcano plots were generated to visualize cluster-specific expression patterns. Heatmaps and volcano plots included in manuscript figures were generated using the ComplexHeatmap (v2.24.0) (<u>Gu et al., 2016)</u> and EnhancedVolcano (v1.26.0) (<u>Blighe et al., 2025</u>) packages.

#### Cell Type Annotation

Cell type annotation was performed iteratively. To ensure confidence in our final cell type annotations, we consulted multiple reference datasets when assessing the most highly expressed genes of each cluster. However, prior to cell type annotation, clusters were grouped by cell class, using lineage-specific markers that have proven to be most consistent when studying rat lung biology (<u>Greaney et al., 2025</u>). We used the following cell class markers: *Epcam* (epithelium), *Col1a1* (mesenchyme), *Ptprc* (immune), and *Cdh5* (endothelium). Following this, cell type annotations were informed by an in-house adult rat lung atlas, created via integrated scRNAseq data from 14 different rat lung samples of both sexes (<u>Obata et al., 2024</u>). Additionally, we leveraged publically-available single-cell transcriptomic atlases, including LungMAP (<u>Ardini-Poleske et al., 2017</u>) and PanglaoDB (<u>Franzén et al., 2019</u>). These atlases provide reference datasets for both pulmonary (LungMAP) and other tissue-specific cell populations (PanglaoDB). Specifically, we utilized the LungMAP repository to cross-reference transcriptional signatures of pulmonary epithelial cell types such as ATI, ATII, basal, and secretory in adult human lung data with those in rat. While some signature genes were conserved (homologs), not all were. This is to be expected given knowledge that humans and rats evolved along different paths and some genes have evolved to have species-specific functions (<u>Church et al., 2024</u>). Similarly, PanglaoDB, served as a useful reference repository for confirming the distinction between immune and endothelial subtypes, beyond the reliance on literature precedent. Justifications for cell type annotations (by gene expression) for our three main single-cell objects can be found in **Supplemental Figs. 1** (BAL), **2** (PD), and **7** (Organoids: BAL_3D, Mixed_3D, PD_3D).

Specific cell type populations, such as the ‘Stressed_Progenitors’, were identified by also using gene set enrichment analysis (GSEA) of upregulated genes in that cluster of cells. This method was performed using the clusterProfiler package (v4.16.0) in R (<u>Yu et al., 2012; Yu, 2024</u>) Final annotations were validated using feature plots and differential gene expression analysis using our global (merged) Seurat object of all three organoid conditions (n=3 for each condition). R markdown files walking through the cleaning, clustering, and annotation process for each sample can be found on our GitHub repository: https://github.com/RaredonLab/Edelstein2025.

### Gene Set Enrichment Analysis (GSEA) with the Molecular Signatures Database (MSigDB)

To better characterize transcriptional programs distinguishing basal hillock cells from luminal hillock cells, we performed gene set enrichment analysis (GSEA; Broad 2022 release, v7.5.1) using the Hallmark gene sets from MSigDB (Liberzon et al., 2015). The pseudobulk BAL_3D and PD_3D/Mixed_3D comparisons were compared using differential expression analysis and genes were ranked by log_2_ fold-change (avg_log2FC). This marker list was then cleaned, ensuring gene symbols were included as an independent column in the data-frame and any row with a missing value (NA) was removed. The resulting data-frame (marker list) was passed through the GSEA() function in the clusterProfiler package (v4.16.0), using the Hallmark gene sets for Rattus norvegicus (msigdbr) pulled from MSigDB (<u>Wu et al., 2021</u>). We defined significantly enriched pathways by adjusted p-values < 0.05.

To better interpret the different components of each upregulated pathway and the implications of pathway upregulation within our systems, genes for each significant term were annotated using curated ground truth gene lists. A set of rat-specific ligands were obtained from the NICHES FANTOM5 reference database (<u>Raredon et al., 2023</u>). For matrix proteins, we curated our own list by aggregating the categories “ECM Glycoproteins,” “Collagens,” and “Proteoglycans” from the Matrisome Project (MatrisomeDB) (<u>Shao et al., 2020</u>). A list of transcription factors was compiled from the AnimalTFDB database (<u>Shen et al., 2023</u>). From here, we identified the number of ligands in each pathway (excluding matrix proteins), the number of matrix-associated genes, and the identity of transcription factors present in the leading-edge gene set. We then identified the number of ligands in each pathway (excluding matrix proteins), the number of matrix-associated genes, and the transcription factors present in the leading-edge gene set. In the context of these results, a positive normalized enrichment score (NES) was interpreted as being enriched in basal hillock cells while a negative NES indicated enrichment in luminal hillock cells.

### Pseudotime Analysis

Trajectory analysis and subsequent analyses of differential gene expression across pseudotime were conducted using the Monocle3, Slingshot, and TradeSeq packages. These analyses allowed for the exploration of cellular transitions within emergent populations of interest such as the hillock cells and pro-/anti-inflammatory macrophages. The combination of these methods utilized provided us with a robust way of inferring lineage associations and transcription-level differences across multiple datasets.

#### Monocle3 Analysis

Monocle3 (v1.3.7) was utilized to infer cellular trajectories and identify key transcriptional changes (<u>Cao et al., 2019</u>). Previously processed Seurat objects for each emergent population were used as input. Raw gene expression counts, and associated cell metadata were extracted from the Seurat objects, and a Monocle3 cell data set (CDS) was constructed with gene annotations.

Dimensionality reduction embeddings, including principal component analysis (PCA) and uniform manifold approximation and projection (UMAP), were transferred from the original Seurat object to the Monocle3 CDS to maintain consistency across analyses. Parameters were set based on the Seurat subset being studied.

#### Monocle3 Analysis: Hillock Subset

To minimize short, noisy bifurcation points, cells were clustered using Monocle3’s clustering algorithm with a minimum branch length of 8. A Euclidean distance ratio of 2.1 and a geodesic distance ratio of 2.1 was used to optimize trajectory learning, and the graph pruning function was enabled to remove spurious connections along the inferred trajectory. Cells were ordered along the inferred trajectory to model their progression through biological states, with the defined starting cluster being the cycling proximal epithelium. Pseudotime values were extracted and added back into the original Seurat metadata.

#### Monocle3 Analysis: Polarized Macrophage Subset

Differential gene expression analysis was performed along the pseudotime trajectory using Moran’s I test with a statistical significance threshold set at a false discovery rate (FDR) of < 0.05. Genes were ranked based on their q-values and prioritized using Moran’s I statistic to identify key regulators of cellular transitions. Genes of interest were analyzed for their dynamic expression patterns across pseudotime. Trajectory plots were generated to visualize temporal expression trends. Monocle3 analysis scripts are available in our <u>GitHub repository</u>.

#### Slingshot Analysis

To independently confirm the trajectory results obtained by Monocle3, slingshot (v2.10.0) was applied to fit principal curves to the UMAP-reduced space (Street et al., 2018). Emergent populations were converted from Subset Seurat objects into SingleCellExperiment (SCE) format for compatibility with Slingshot; PCA and UMAP embeddings were carried over, and the cluster labeled “Cycling” was used as the starting cluster for pseudotime ordering (Hillock analysis).

Slingshot inferred lineage relationships based on cluster-based lineage tracing guided by the cell type annotations. Default smoothing splines were applied to fit the trajectory curves and pseudotime values were extracted and included in Seurat metadata for comparisons with Monocle3-derived pseudotime trajectories. Differential gene expression analysis along Slingshot-derived trajectories was conducted using TradeSeq (v1.16.0), retaining genes expressed in at least 1% of cells with counts greater than 5 (Van den Berge et al., 2020). The number of knots for smoothing was set to 15 based on the Akaike Information Criterion (AIC) that we determined by comparing models using different numbers of knots from 3 to 15. A test of association was performed between pseudotime and genes with a significant association (FDR < 0.05), followed by start-vs-end test to evaluate gene expression changes between early vs late states in the pseudotime trajectory. Scripts for Slingshot analysis are available in our GitHub repository.

### Cell—Cell Connectomic Analysis with NICHES

NICHES (v0.2.3) was used to infer ligand-receptor-mediated cell-cell communication across experimental conditions. All analyses were conducted in R (v4.2.2) using Seurat (v4.3.0) and SeuratWrappers (v.0.4.0). Global NICHES analysis was performed on an integrated Seurat object of all engineered cells (PD_3D, Mixed_3D, BAL_3D, 12 samples total; n = 3 per condition). A subset analysis was conducted on a subset of object specific populations of interest to afford a finer analysis of cell-cell connectomics.

#### Global Analysis

Cells were first subset by condition and split by replicate (Orig_ID). For each replicate, cell-cell signaling was inferred using the RunNICHES function with the FANTOM5 ligand-receptor database and the Rnor_6.0 rat reference genome. We enabled only the CellToCell inference mode to focus on direct intercellular signaling. Outputs were merged across replicates and combined using JoinLayers. Cells with low signaling information (nFeature_CellToCell < 60) were removed. The resulting object was normalized, scaled, and run through PCA (100 components). UMAP embedding and unsupervised clustering were performed using the top 18 principal components (resolution = 0.4). To correct for batch effects, RPCA integration was performed by condition (PCs 1-16, 18-19) and by sample (PCs 1-14, 16-18). The final integrated object (eng.CTC.final) served as the basis for downstream signaling analyses. The script for this analysis can be found in our <u>GitHub repository.</u>

#### Focused Subset Analysis

To study regenerative signaling in more detail, a focused analysis was performed on a subset of six cell populations from our condition-level annotations: Hillock_Luminal, Hillock_Basal, Rspo3+_Mes, Pdgfrb+_Pericytes, Anti_Inflamm_Mac, and Pro_Inflamm_Mac. These subsets were extracted from our global, node-aligned object. Following normalization and feature selection, PCA was performed, and PCs 1:5, 7, 9-12, and 14 were used for UMAP and clustering (resolution = 0.4). NICHES was then run per replicate for each condition. Cell types with fewer than two cells per replicate were excluded. Merged CellToCell results were filtered to retain cells with ≥40 features and processed using PCA (PCs 1:10, 13-14, 17-18, 21, 23), UMAP, and clustering. The final object (regen.subset_CTC_byCondition) was used for downstream visualization and differential signaling analysis using a composite “power” metric (avg_log2FC × expression ratio). The script for this analysis can be found in our <u>GitHub repository.</u>

#### Cell Circuit-Level Visualization

We visualized signaling mechanisms of interest as directed multicellular circuits (Fig. 6) using the CircuitPlot() function in the <u>NICHESMethods GitHub repository</u> (<u>Raredon et al., 2023; Greaney et al., 2024</u>). This function uses a custom plotting algorithmn (ggCircuit) that employs trigonometric positioning, with the help of ggplot2 (<u>Wickham, 2010</u>) rendering, to visualize ligand-receptor interactions (e.g., Il1a—Il1r2) as directional arrows connecting cell types defined by a metadata grouping variable (e.g., CellType) and edge aesthetics that reflect connection strength. Layout parameters such as graph.angle, offset, and unity.normalize were occasionally passed to the function, when appropriate, and all circuits were generated from the subset connectomic object (regen.subset_CTC_bySample) using annotations from CellType.regen.spec to ensure consistency in node identity and classification.

## Data Availability

Raw FASTQ files, extracted digital gene expression matrices, and all R objects, containing relevant meta-data, used in this manuscript are available at Gene Expression Omnibus GEO *******. Scripts used for the analysis and figures have been made publicly available on GitHub at https://github.com/RaredonLab/Edelstein2025.

## Acknowledgments

We would like to thank Yale Center for Genome Analysis (YCGA), the Yale Pathology Tissue Services (YPTS), and Yale Center for Research Computing (YCRC) for their efforts in making this work possible. We would also like to acknowledge Dr. Allison Greaney and Dr. Themis Kyriakides for their feedback and support in story craft, single-cell analysis methodology, and overall mentorship. We would like to acknowledge Ako Ndefo-Haven for his assistance with line-editing.

## Author Contributions

Conceptualization: S.E.E., M.S.B.R, S.M., M.S.; Methodology: S.E.E., M.S.B.R., S.M., H.K., N.W.; Software: S.E.E., M.S.B.R., N.W., H.K.; Validation: N.W., H.K., S.M., C.H.; Formal Analysis: S.E.E., N.W., M.S.B.R; Investigation: S.E.E., S.M., M.T.G, C.D.; Resources: M.S.B.R; Writing (Original Draft): S.E.E., M.S.B.R; Writing (Review & Editing): S.E.E., M.S.B.R., M.S., V.L., C.H., C.D.

## Supplemental Figures

**Supplemental Figure 1.**
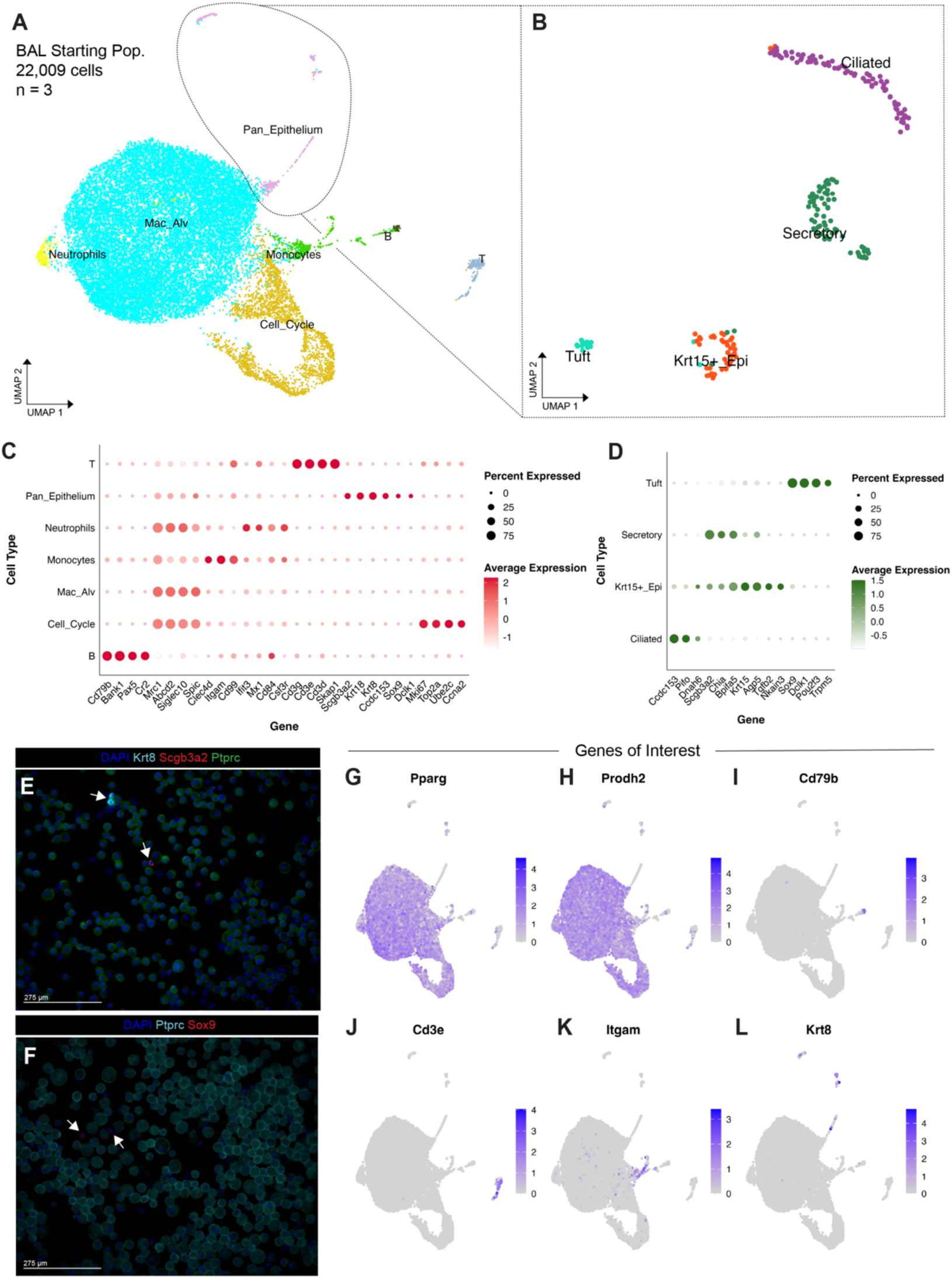
Characterization of the BAL starting population prior to 3D culture. **(A)** UMAP vIsualIzalion of 22,009 single cells derived from bronchoalveolar lavage (BAL) across three biological replicates, identifying major populations including Pan_Epithelium. Macrophages (Mac_Alv). Monocytes, Neutrophils, B cells, and proliferative (Cell_Cycle) cells. **(B)** Sub-clustering of the Pan_Epithelium population from **(A)** reveals distinct epithelial subsets, including Ciliated, Secretory, Krt15+_Epi, and Tuft cells. **(C-D)** Dot plots displaying expression of canonical marker genes across major cell types **(C)** and epithelial subclusters **(D)**. Dot size refiects the percentage of cells expressing the gene; color indicates scaled average expression. **(E-F)** lmmunofluorescence validation of epithelial and immune cell markers in BAL starting population. **(E)** Co-staining for Krt8 (cyan), Scgb3a2 (red), and Plprc (green) confirms the presence of both epithelial and immune cells; **(F)** Plprc (cyan) and Sox9 (red) highlight rare Sox9+ cells within the population. Arrows highlight rare epithelial cells. DAPI (blue) marks nuclei. Scale bars= 275 µm.

**Supplemental Figure 2.**
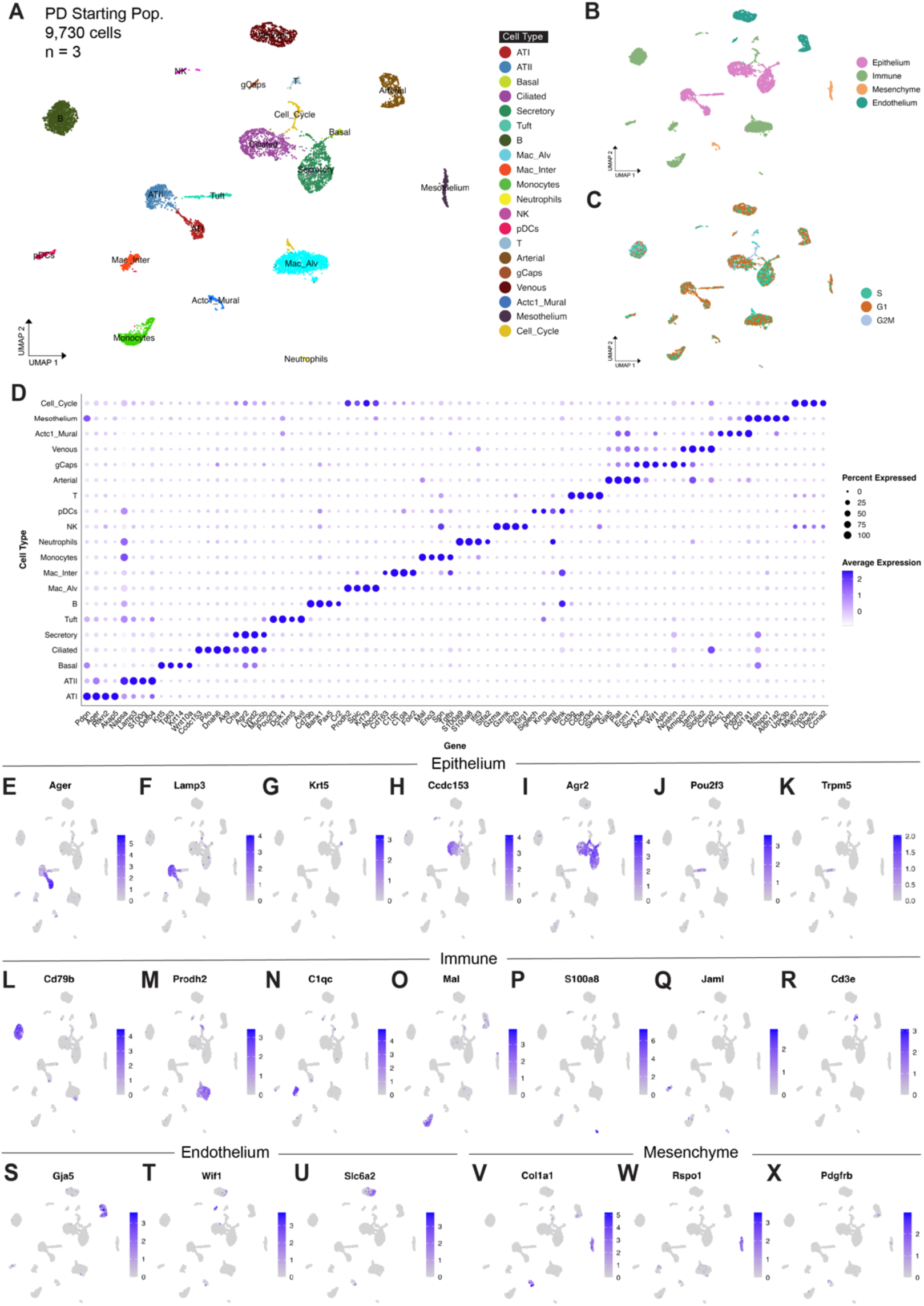
Single-cell characterization of the PD starting population prior to 3D culture. **(A)** UMAP embedding of 9,730 single cells derived from the pulmonary dissociation (PD) starting condition across three biological replicates, colored by cell type annotation. Cell populations include epithelial (ATI, ATII, Basal, Secretory, Ciliated, Tuft), immune (Macrophages, Monocytes, Neutrophils, NK, pDCs, B, T), mesenchymal (Actc1_Mural, Mesothelium), endothelial (gCaps, aCaps, Arterial, Venous), and proliferating (Cell_Cycle) cells. **(B)** UMAP colored by broad cell class: Epithelium, Immune, Endothelium, and Mesenchyme. (C) UMAP showing cell cycle phase assignment (G1, S, G2M). **(D)** Dot plot displaying expression of canonical markers across annotated cell types. Dot size indicates the percentage of cells expressing each gene, and color intensity represents average expression. **(E-K)** Feature plots showing expression of representative epithelial markers. **(L-R)** Feature plots for immune-related genes of interest. **(S-U)** Feature plots showing endothelial marker markers of interest. **(V-X)** Feature plots showing mesenchymal markers that are representative of the population including Col1a1, Rspo1, and Pdgfrb.

**Supplemental Figure 3.**
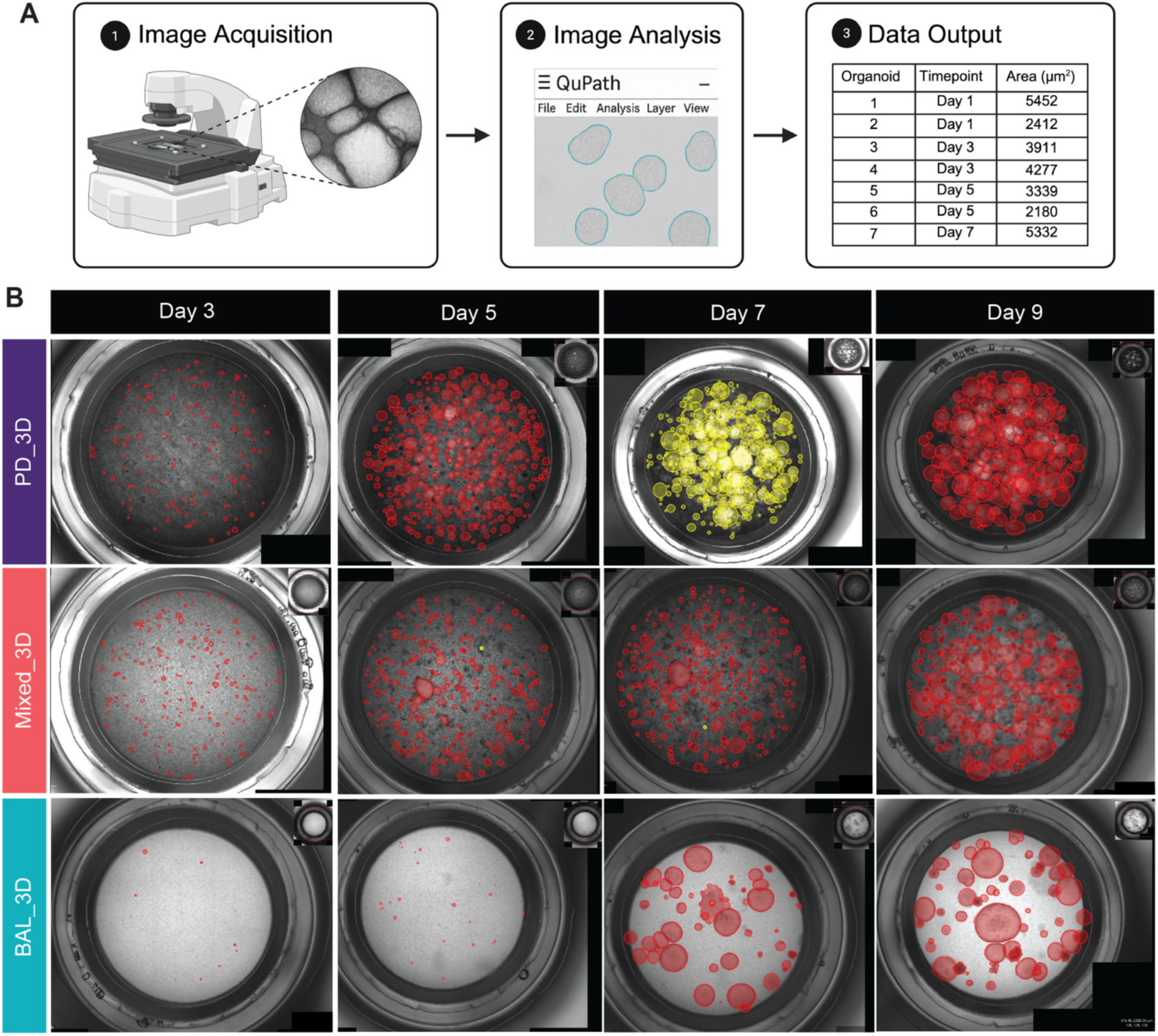
Organoid size quantification using QuPath. **(A)** Schematic overview of the image analysis workflow. Brightfield images were acquired at multiple timepoints (Days 0, 3, 5, 7, and 9) using the EVOS FL Auto 2 microscope (4x objective). Stitched images were imported into QuPath (v0.5.1), where organoid boundaries were annotated using the ellipse and brush tools. Organoid area was calculated using a calibrated scale of 1.63 µm/pixel based on the embedded **1** mm scale bar. Quantified area measurements (µm^2^) were exported for downstream statistical analysis. **(B)** Representative stitched images of entire wells across three culture conditions, PD_3D, Mixed_3D, and BAL_3D, at Days 3, 5, 7, and 9. Red (and yellow, Day 7) overlays indicate detected organoids and their measured boundaries. Variability in organoid number and size is evident across both conditions and timepoints.

**Supplemental Figure 4.**
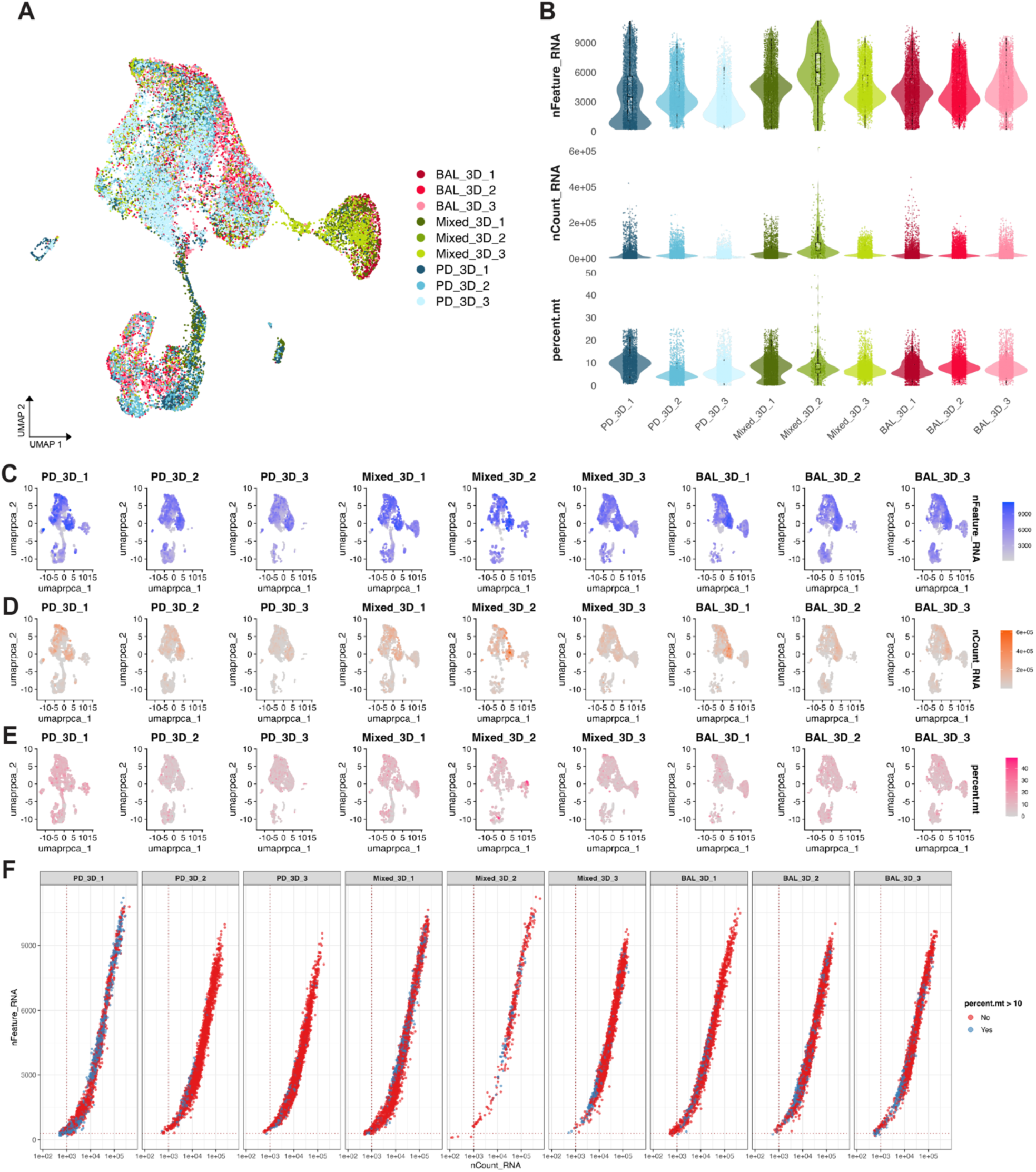
Quality control and sample overview of engineered organoid dataset. **(A)** Integrated UMAP embedding of all engineered samples grouped and colored by experimental replicate (Orig_lD). Samples include BAL-derived (BAL_3D_1-3), Mixed (Mixed_3D_1-3), and PD-derived (PD_3D_1-3) organoids. **(B)** Violin plots showing the distributions of raw (untransformed) values by sample (replicate): number of genes detected per cell (nFeature_RNA), total RNA counts (nCount_RNA), and percentage of mitochondrial transcripts (percent.mt). Boxplots indicate interquartile range and median; individual cells are overlaid as jittered points. **(C-E)** Split UMAP feature plots for each sample showing expression of nFeature_RNA **(C)**, nCount_RNA **(D)**, and percent.mt **(E). (F)** Scatter plots showing nCount_RNA (log10 scale) versus nFeature_RNA (linear scale) for individual cells, faceted by sample (Orig_lD). Cells are colored by mitochondrial transcript abundance, with blue indicating cells exceeding 10% mitochondrial RNA (percent.mt> 10). Red dotted lines indicate commonly used quality control thresholds (nFeature_RNA =300, nCount_RNA = 1,000).

**Supplemental Figure 5.**
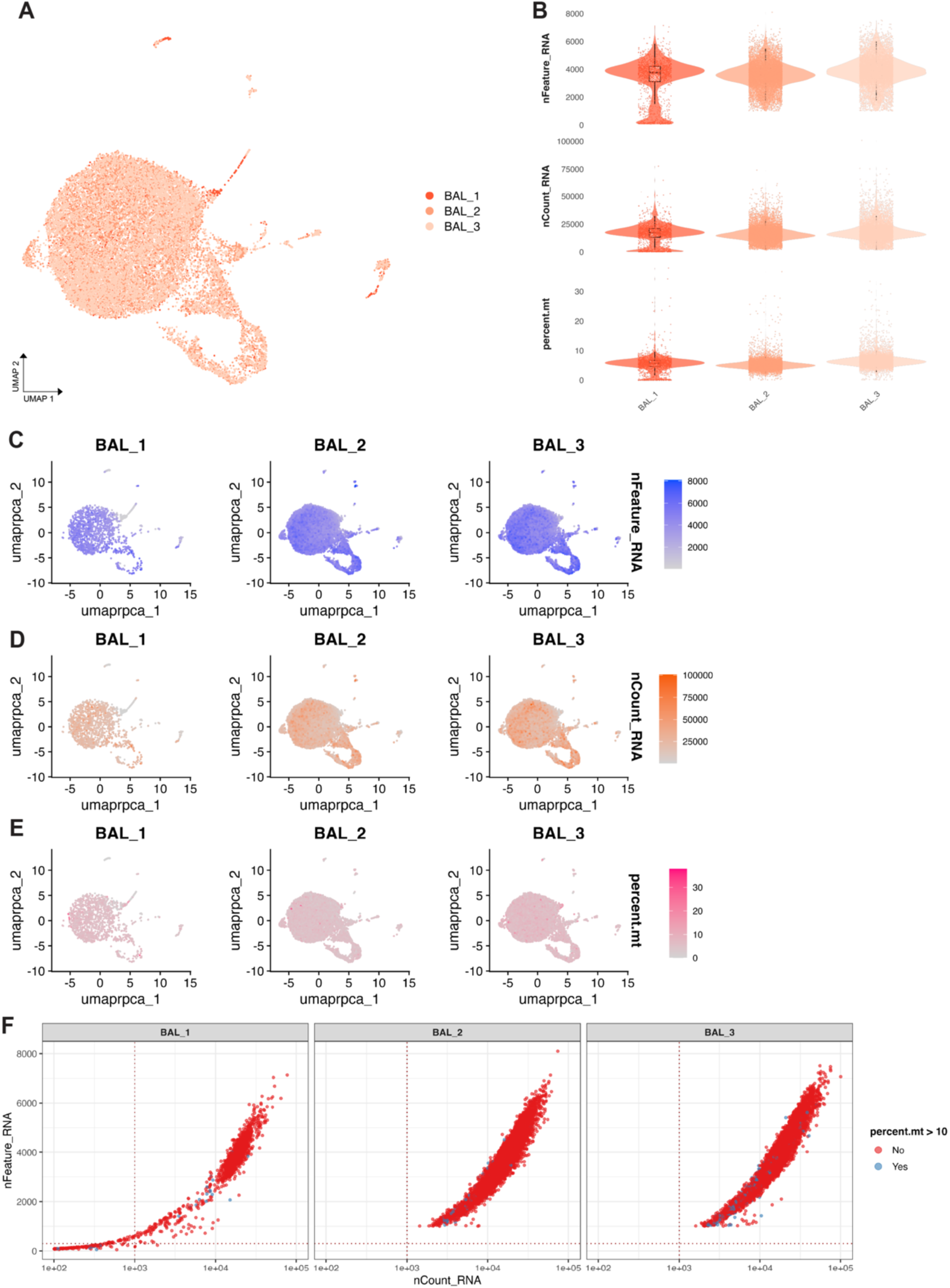
Quality control and sample overview of BAL starting population. **(A)** Integrated UMAP embedding of all BAL starting cells grouped and colored by experimental replicate (Orig_lD; BAL_1-3). **(B)** Violin plots showing the distributions of raw (untransformed) quality control metrics across replicates: number of genes detected per cell (nFeature_RNA), total RNA counts (nCount_RNA), and percentage of mitochondrial transcripts (percent.mt). Boxplots indicate interquartile range and median; individual cells are overlaid as jittered points. **(C-E)** Split UMAP feature plots for each replicate showing expression of nFeature_RNA (C), nCount_RNA **(D),** and percent.mt **(E). (F)** Scatter plots showing nCount_RNA (10910 scale) versus nFeature_RNA (linear scale) for individual cells, faceted by replicate. Cells are colored by mitochondrial transcript abundance, with blue indicating cells exceeding 10% mitochondrial RNA (percent.mt> 10). Red dotted lines indicate quality control thresholds used for filtering (nFeature_RNA = 300, nCount_RNA = 1,000, percent.mt= 10%).

**Supplemental Figure 6.**
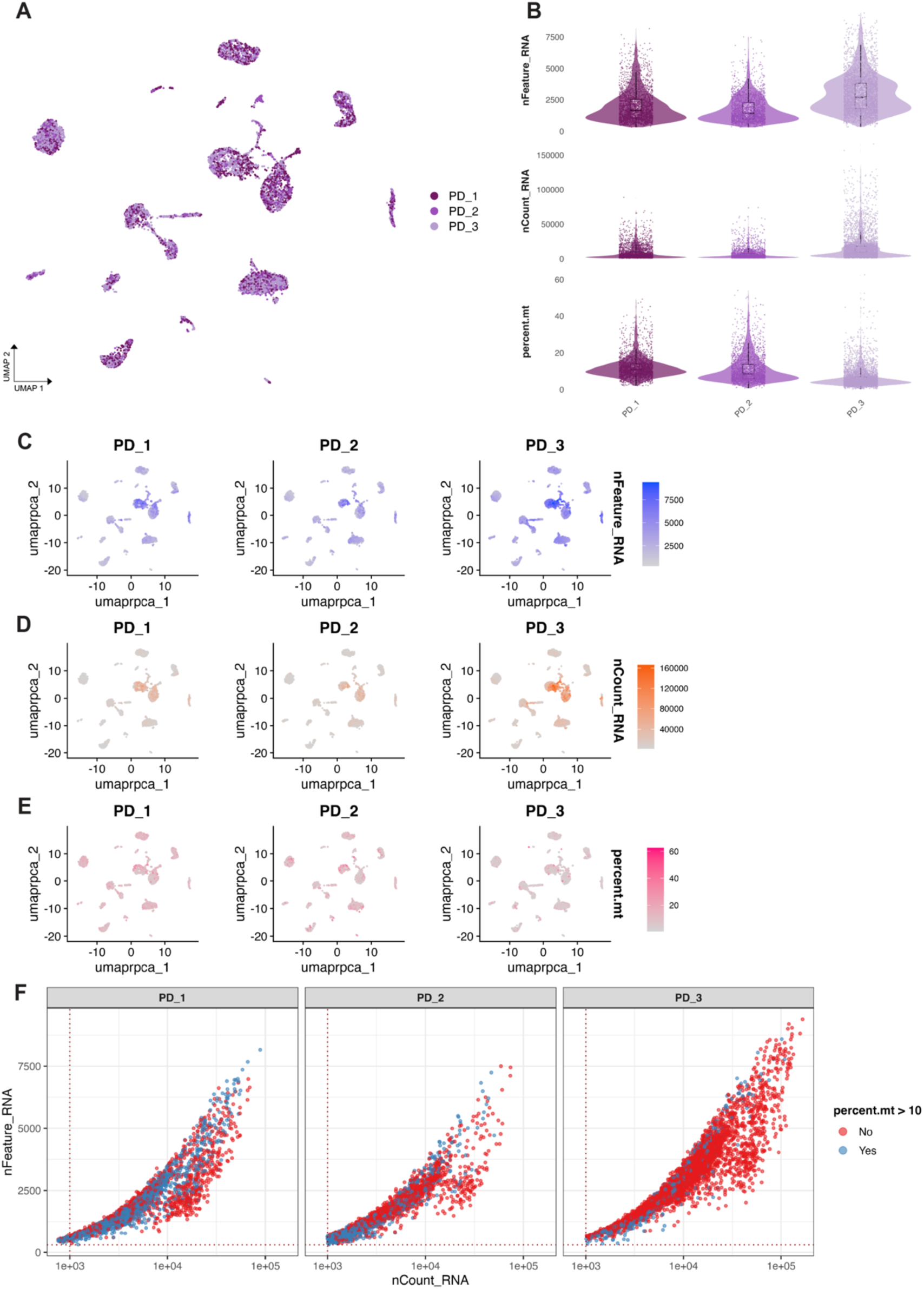
Quality control and sample overview of PD starting population. **(A)** UMAP embedding of all pulmonary dissociation (PD) starting cells, grouped and colored by experimental replicate (Orig_lD; PD_1-3). **(B)** Violin plots showing the distributions of raw (untransformed) quality control metrics across replicates: number of genes detected per cell (nFeature_RNA), total RNA counts (nCount_RNA), and percentage of mitochondrial transcripts (percent.mt). Boxplots indicate interquartile range and median; individual cells are overlaid as jittered points. **(C-E)** Split UMAP feature plots for each replicate showing expression of nFeature_RNA (C), nCount_RNA **(D),** and percent.mt **(E). (F)** Scatter plots showing nCount_RNA (log10 scale) versus nFeature_RNA (linear scale) for individual cells, faceted by replicate (PD_1-3). Cells are colored by mitochondrial transcript abundance, with blue indicating cells exceeding 10% mitochondrial RNA (percent.mt> 10). Red dotted lines indicate commonly used quality control thresholds (nFeature_RNA = 300, nCount_RNA = 1,000).

**Supplemental Figure 7.**
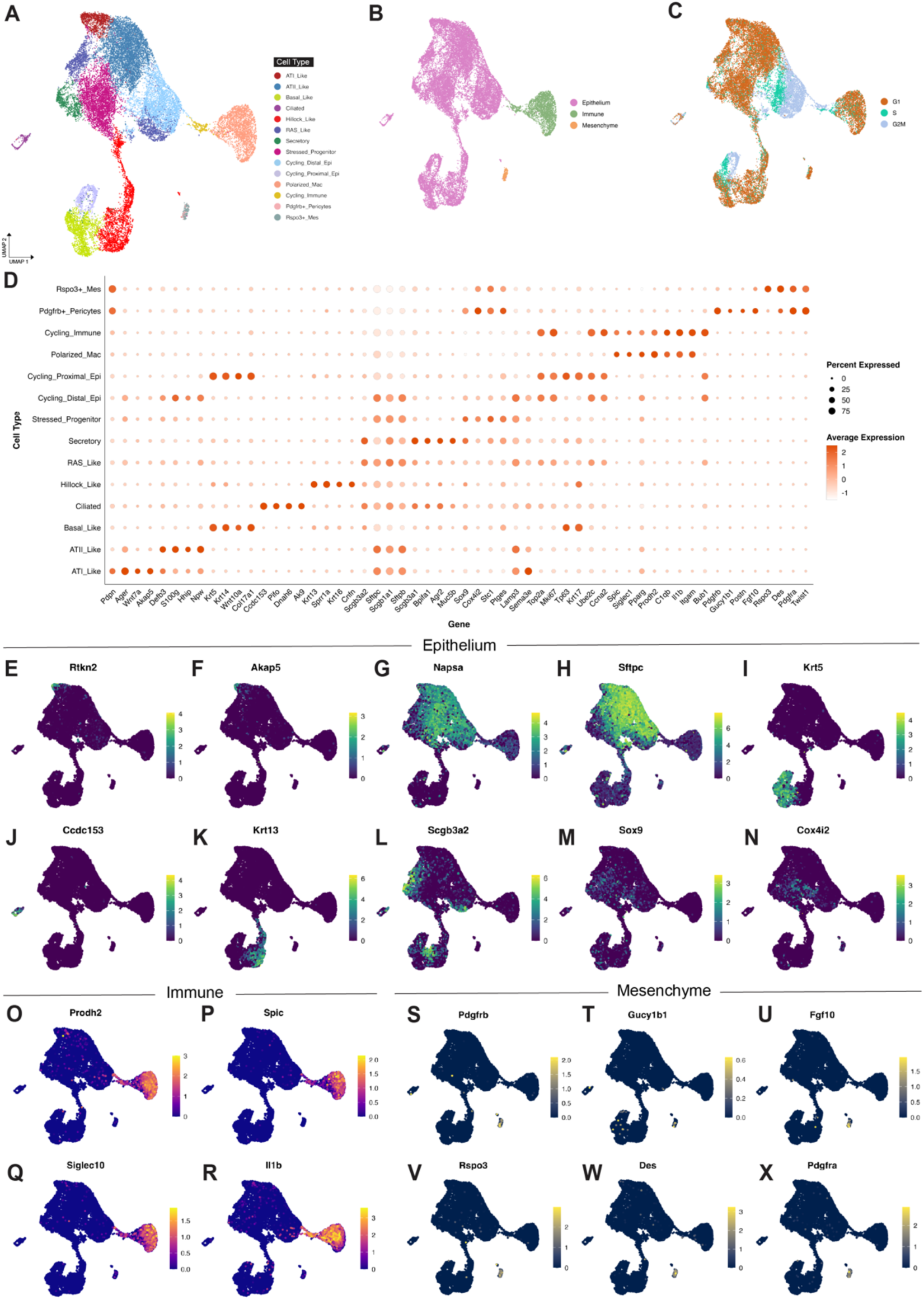
Marker-based cell type annotation in engineered organoid single-cell dataset (global object, all three organoid conditions). **(A)** Integrated UMAP embedding of all cells from engineered organoid samples colored by annotated cell type, including epithelial (e.g., ATI_Like, ATII_Like, Secretory, Hillock_Like. RAS_Like), immune (e.g., Polarized_Mac, Cycling_lmmune), and mesenchymal (e.g., Rspo3+_Mes, Pdgfrb+_Pericytes) populations. **(B)** Same UMAP colored by broad cell class: Epithelium, Immune, and Mesenchyme. (C) UMAP colored by cell cycle phase assignment (G1, S, G2M). **(D)** Dot plot showing expression of canonical marker genes across annotated cell types for justification. Dot size indicates the proportion of cells expressing the gene; color reflects scaled average expression. **(E-N)** Feature plots showing expression of epithelial markers including Rtkn2 (ATI), Akap5 (ATI), Napsa (ATII), Sftpc (ATII), Krt5 (Basal), Ccdc153 (Ciliated), Krt13 (Hillock), Scgb3a2 (Secretory/RAS), Sox9 (Stressed_Progenitor), and Cox4i2 (Stressed_Progenitor). **(O-R)** Expression of immune-associated genes including Prodh2, Spic, Siglec10, and 111b. **(S-X)** Feature plots of mesenchymal markers such as Pdgfrb (pericyte), Gucy1b1 (pericyte), Fgf10, Rspo3, Des, and Pdgfra, supporting identification of mesenchymal subpopulations.

**Supplemental Figure 8.**
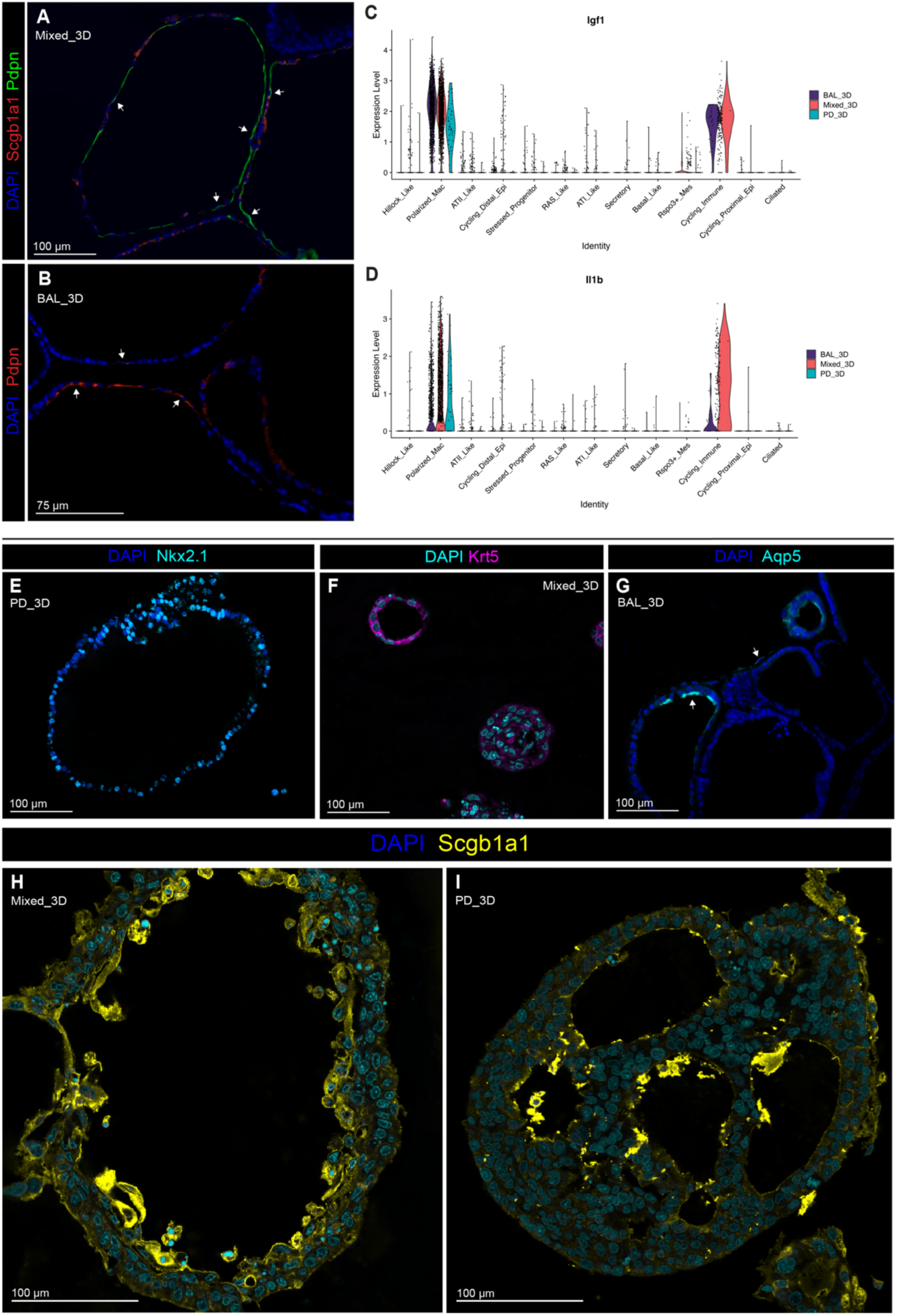
Drivers of ATl-like differentiation and additional epithelial immunofluorescence staining across organoid conditions. **(A-B)** lmmunofluorescence staining of day 10 organoids reveal spatial separation between Pdpn+ alveolar type I (ATI) cells (green) and Scgb1a1+ secretory cells (red), with arrowheads indicating Pdpn+ segments along the epithelial border in **(A)** Mixed_3D and **(B)** BAL_3D cultures. (C-O) Violin plots showing expression of lgf1 and 111b across immune populations. lgf1 expression is highest in the BAL_3D condition. **(E-G)** lmmunofluorescence staining of day 10 organoids for additional epithelial lineage markers: **(E)** Nkx2.1 (cyan) marks ATll-like cells; **(F)** Krt5 (magenta) labels basal-like cells; **(G)** Aqp5 (cyan) marks ATl-like cells, with arrows highlighting localized expression along squamous regions of the epithelial border. **(H-1)** lmmunofluorescence staining for Scgb1a1 (yellow) in **(H)** Mixed_3D and **(I)** PD_3D conditions shows variable distribution of secretory epithelial cells across organoid structures.

## Supplemental Tables

**Supplemental Table S1.** LPM-3D Growth Medium Components (adapted from Nichane et al., 2017)

| Classifier | Component | Concentration | Supplier | Ref. # |
| --- | --- | --- | --- | --- |
| Basal Medium | Advanced DMEM/F12 | N/A | ThermoFisher | 12634010 |
| Growth Factor (GF) | FGF10 | 50 ng/mL | Peprotech | 400-42 |
| GF | FGF9 | 50 ng/mL | Novus Biologics | NBP-35196 |
| GF | EGF | 50 ng/mL | Peprotech | 400-25 |
| Small Molecule (SM) | CHIR99021 (GSK-3 Inhibitor) | 3 μM | Cayman Chemicals | 13122-10 |
| SM | BIRB796 | 1 μM | Cayman Chemicals | 10460-10 |
| SM | Y27632 (Rock Inhibitor) | 10 μM | Cayman Chemicals | 10005583-10 |
| SM | A8301 (ALK-5/TGF-beta/NODAL Inhibitor) | 1 μM | Cayman Chemicals | 9001799-10 |
| Additional Components (AC) | Heparin | 5 μg/mL | Sigma-Aldrich | H3149-10KU |
| AC | Insulin | 10 μg/mL | Sigma-Aldrich | 910077C-100 |
| AC | Transferrin | 15 μg/mL | Sigma-Aldrich | T8158-100 |
| Antibiotics (AB) | PenStrep | 1x | ThermoFisher | 15140122 |
| AB | Glutamine | 1x | ThermoFisher | 25030081 |
| AB | Anti-Anti | 0.25 mg/L | ThermoFisher | 15240062 |
| AB | G418 | 50 μg/mL | ThermoFisher | 10131035 |

**Supplemental Table S2.** Primary Antibodies with Dilutions Used for Immunohistochemistry.

| **Antibody** | **Supplier** | **Product #/**<br>**Identifier** | **Cell Type** | **Dilution** |
| --- | --- | --- | --- | --- |
| Goat anti-Foxj1 | Santa Cruz<br>Biotechnolo<br>gy, Inc. | sc-54371 | Ciliated<br>epithelium<br>(nuclear) | 1:100 |
| Rabbit anti-CD45<br>(Ptprc) | Abcam | ab10558 | Pan-immune | 1:100 |
| Mouse anti-CD45<br>(Ptprc) | BD<br>Biosciences | 554875 | Pan-immune | 1:100 |
| Goat anti-CCSP<br>(Uteroglobin /<br>Scgb1a1) | EMD<br>Millipore | ABS1673 | Secretory cells | 1:100 |
| Goat anti-UGRP1/<br>SCGB3A2 | R&D<br>Systems | AF3545 | Secretory cells | 1:200 |
| Mouse anti-RTI-40<br>(Podoplanin) | Terrace<br>Biotech | TB-11ART1-40 | Alveolar type II<br>cells (rat<br>specific) | 1:40 |
| Mouse anti-Acetylated Tubulin | Sigma | T7451 | Epithelium; cilia<br>specific. | 1:200 |
| Rabbit anti-Krt13 | Abcam | ab92551 | Hillock cells | 1:100 |
| Mouse anti-Krt14 | Abcam | ab9220 | Basal cells | 1:100 |
| Rabbit anti-proSPC | Millipore | AB3786 | ATII cells | 1:100 |
| Rabbit anti-Sox9 | Abcam | ab185966 | BASCs | 1:100 |
| Rabbit anti-Krt8 | Abcam | ab59400 | Transitional<br>epithelium | 1:100 |

**Supplemental Table S3.** Secondary Antibodies with Dilutions Used for Immunohistochemistry.

| **Antibody** | **Supplier** | **Product #/**<br>**Identifier** | **Species** | **Dilution** |
| --- | --- | --- | --- | --- |
| Donkey anti-Rabbit IgG (H+L)<br>Alexa Fluor 647 | Invitrogen | A31573 | Donkey | 1:500 |
| Donkey Anti-Goat IgG (H+L), Alexa Fluor 555 | Invitrogen | A21432 | Donkey | 1:500 |
| Donkey Anti-Mouse IgG (H&L), DyLight 488 | Abcam | ab96875 | Donkey | 1:500 |
| Donkey Anti-Mouse IgG (H+L), Alexa Fluor 488 | Invitrogen | A21202 | Donkey | 1:500 |

**Supplemental Table 4 (S4):** Initial cutoffs and final clustering resolutions used for each individual scRNAseq sample.

| **Sample ID** | **nCount_RNA Initial Cutoff** | **Final Embedding Resolution** |
| --- | --- | --- |
| PD_1 | **750** | **0.5** |
| PD_2 | **1000** | **0.2** |
| PD_3 | **1000** | **0.35** |
| BAL_1 | **100** | **0.5** |
| BAL_2 | **1000** | **0.3** |
| BAL_3 | **1000** | **0.3** |
| PD_3D_1 | **500** | **0.5** |
| PD_3D_2 | **500** | **0.8** |
| PD_3D_3 | **500** | **0.45** |
| Mixed_3D_1 | **500** | **0.42** |
| Mixed_3D_2 | **100** | **0.4** |
| Mixed_3D_3 | **100** | **0.5** |
| BAL_3D_1 | **500** | **0.38** |
| BAL_3D_2 | **500** | **0.5** |
| BAL_3D_3 | **500** | **0.5** |

## References

Abo, K. M., Sainz de Aja, J., Lindstrom-Vautrin, J., Alysandratos, K. D., Richards, A., Garcia-de-Alba, C., Huang, J., Hix, O. T., Werder, R. B., Bullitt, E., Hinds, A., Falconer, I., Villacorta-Martin, C., Jaenisch, R., Kim, C. F., Kotton, D. N., & Wilson, A. A. (2022). Air-liquid interface culture promotes maturation and allows environmental exposure of pluripotent stem cell-derived alveolar epithelium. JCI Insight, 7(6). 10.1172/jci.insight.155589

Ahmed, D. W., Eiken, M. K., DePalma, S. J., Helms, A. S., Zemans, R. L., Spence, J. R., Baker, B. M., & Loebel, C. (2023). Integrating mechanical cues with engineered platforms to explore cardiopulmonary development and disease. iScience, 26(12), 108472. 10.1016/j.isci.2023.108472

Alysandratos, K. D., Herriges, M. J., & Kotton, D. N. (2021). Epithelial Stem and Progenitor Cells in Lung Repair and Regeneration. Annu Rev Physiol, 83, 529–550. 10.1146/annurev-physiol-041520-092904

Anton, O., Batista, A., Millan, J., Andres-Delgado, L., Puertollano, R., Correas, I., & Alonso, M. A. (2008). An essential role for the MAL protein in targeting Lck to the plasma membrane of human T lymphocytes. Journal of Experimental Medicine, 205(13), 3201–3213. 10.1084/jem.20080552

Aras, S., Pak, O., Sommer, N., Finley, R., Jr., Huttemann, M., Weissmann, N., & Grossman, L. I. (2013). Oxygen-dependent expression of cytochrome c oxidase subunit 4-2 gene expression is mediated by transcription factors RBPJ, CXXC5 and CHCHD2. Nucleic Acids Res, 41(4), 2255–2266. 10.1093/nar/gks1454

Ardini-Poleske, M. E., Clark, R. F., Ansong, C., Carson, J. P., Corley, R. A., Deutsch, G. H., Hagood, J. S., Kaminski, N., Mariani, T. J., Potter, S. S., Pryhuber, G. S., Warburton, D., Whitsett, J. A., Palmer, S. M., Ambalavanan, N., & Lung, M. A. P. C. (2017). LungMAP: The Molecular Atlas of Lung Development Program. Am J Physiol Lung Cell Mol Physiol, 313(5), L733–L740. 10.1152/ajplung.00139.2017

Ayaub, E. A., Dubey, A., Imani, J., Botelho, F., Kolb, M. R. J., Richards, C. D., & Ask, K. (2017). Overexpression of OSM and IL-6 impacts the polarization of pro-fibrotic macrophages and the development of bleomycin-induced lung fibrosis. Sci Rep, 7(1), 13281. 10.1038/s41598-017-13511-z

Babaev, V. R., Fazio, S., Gleaves, L. A., Carter, K. J., Semenkovich, C. F., & Linton, M. F. (1999). Macrophage lipoprotein lipase promotes foam cell formation and atherosclerosis in vivo. J Clin Invest, 103(12), 1697–1705. 10.1172/JCI6117

Barnes, J. L., Yoshida, M., He, P., Worlock, K. B., Lindeboom, R. G. H., Suo, C., Pett, J. P., Wilbrey-Clark, A., Dann, E., Mamanova, L., Richardson, L., Polanski, K., Pennycuick, A., Allen-Hyttinen, J., Herczeg, I. T., Arzili, R., Hynds, R. E., Teixeira, V. H., Haniffa, M.,…Nikolic, M. Z. (2023). Early human lung immune cell development and its role in epithelial cell fate. Science Immunology, 8(90), eadf9988. 10.1126/sciimmunol.adf9988

Basil, M. C., Cardenas-Diaz, F. L., Kathiriya, J. J., Morley, M. P., Carl, J., Brumwell, A. N., Katzen, J., Slovik, K. J., Babu, A., Zhou, S., Kremp, M. M., McCauley, K. B., Li, S., Planer, J. D., Hussain, S. S., Liu, X., Windmueller, R., Ying, Y., Stewart, K. M.,…Morrisey, E. E. (2022). Human distal airways contain a multipotent secretory cell that can regenerate alveoli. Nature, 604(7904), 120–126. 10.1038/s41586-022-04552-0

Belperio, J. A., Keane, M. P., Burdick, M. D., Londhe, V., Xue, Y. Y., Li, K., Phillips, R. J., & Strieter, R. M. (2002). Critical role for CXCR2 and CXCR2 ligands during the pathogenesis of ventilator-induced lung injury. J Clin Invest, 110(11), 1703–1716. 10.1172/JCI15849

Beppu, A. K., Zhao, J., Yao, C., Carraro, G., Israely, E., Coelho, A. L., Drake, K., Hogaboam, C. M., Parks, W. C., Kolls, J. K., & Stripp, B. R. (2023). Epithelial plasticity and innate immune activation promote lung tissue remodeling following respiratory viral infection. Nat Commun, 14(1), 5814. 10.1038/s41467-023-41387-3

Block, G. J., Ohkouchi, S., Fung, F., Frenkel, J., Gregory, C., Pochampally, R., DiMattia, G., Sullivan, D. E., & Prockop, D. J. (2009). Multipotent stromal cells are activated to reduce apoptosis in part by upregulation and secretion of stanniocalcin-1. Stem Cells, 27(3), 670–681. 10.1002/stem.20080742

Boon, K., Vanalken, N., Szpakowska, M., Chevigne, A., Schols, D., & Van Loy, T. (2025). High-affinity ELR+ chemokine ligands show G protein bias over beta-arrestin recruitment and receptor internalization in CXCR1 signaling. J Biol Chem, 301(1), 108044. 10.1016/j.jbc.2024.108044

Buckley, S., Shi, W., Barsky, L., & Warburton, D. (2008). TGF-beta signaling promotes survival and repair in rat alveolar epithelial type 2 cells during recovery after hyperoxic injury. Am J Physiol Lung Cell Mol Physiol, 294(4), L739–748. 10.1152/ajplung.00294.2007

Burri, P. H. (1974). The postnatal growth of the rat lung. 3. Morphology. Anat Rec, 180(1), 77–98. 10.1002/ar.1091800109

Cai, X. T., Jia, M., Heigl, T., Shamir, E. R., Wong, A. K., Hall, B. M., Arlantico, A., Hung, J., Menon, H. G., Darmanis, S., Brightbill, H. D., Garfield, D. A., & Rock, J. R. (2024). IL-4-induced SOX9 confers lineage plasticity to aged adult lung stem cells. Cell Rep, 43(8), 114569. 10.1016/j.celrep.2024.114569

Camper, N., Glasgow, A. M., Osbourn, M., Quinn, D. J., Small, D. M., McLean, D. T., Lundy, F. T., Elborn, J. S., McNally, P., Ingram, R. J., Weldon, S., & Taggart, C. C. (2016). A secretory leukocyte protease inhibitor variant with improved activity against lung infection. Mucosal Immunology, 9(3), 669–676. 10.1038/mi.2015.90

Cao, X., Surma, M. A., & Simons, K. (2012). Polarized sorting and trafficking in epithelial cells. Cell Res, 22(5), 793–805. 10.1038/cr.2012.64

Cao, J., Spielmann, M., Qiu, X., Huang, X., Ibrahim, D. M., Hill, A. J., Zhang, F., Mundlos, S., Christiansen, L., Steemers, F. J., Trapnell, C., & Shendure, J. (2019). The single-cell transcriptional landscape of mammalian organogenesis. Nature, 566(7745), 496–502. 10.1038/s41586-019-0969-x

Choi, J., Park, J. E., Tsagkogeorga, G., Yanagita, M., Koo, B. K., Han, N., & Lee, J. H. (2020). Inflammatory Signals Induce AT2 Cell-Derived Damage-Associated Transient Progenitors that Mediate Alveolar Regeneration. Cell Stem Cell, 27(3), 366–382 e367. 10.1016/j.stem.2020.06.020

Church, S. H., Mah, J. L., & Dunn, C. W. (2024). Integrating phylogenies into single-cell RNA sequencing analysis allows comparisons across species, genes, and cells. PLoS Biol, 22(5), e3002633. 10.1371/journal.pbio.3002633

Corro, C., Novellasdemunt, L., & Li, V. S. W. (2020). A brief history of organoids. Am J Physiol Cell Physiol, 319(1), C151–C165. 10.1152/ajpcell.00120.2020

de Kleer, I. M., Kool, M., de Bruijn, M. J., Willart, M., van Moorleghem, J., Schuijs, M. J., Plantinga, M., Beyaert, R., Hams, E., Fallon, P. G., Hammad, H., Hendriks, R. W., & Lambrecht, B. N. (2016). Perinatal Activation of the Interleukin-33 Pathway Promotes Type 2 Immunity in the Developing Lung. Immunity, 45(6), 1285–1298. 10.1016/j.immuni.2016.10.031

Deguchi, E., Lin, S., Hirayama, D., Matsuda, K., Tanave, A., Sumiyama, K., Tsukiji, S., Otani, T., Furuse, M., Sorkin, A., Matsuda, M., & Terai, K. (2024). Low-affinity ligands of the epidermal growth factor receptor are long-range signal transmitters in collective cell migration of epithelial cells. Cell Rep, 43(11), 114986. 10.1016/j.celrep.2024.114986

Di-Luoffo, M., Ben-Meriem, Z., Lefebvre, P., Delarue, M., & Guillermet-Guibert, J. (2021). PI3K functions as a hub in mechanotransduction. Trends Biochem Sci, 46(11), 878–888. 10.1016/j.tibs.2021.05.005

Dong, P., Maddali, M. V., Srimani, J. K., Thelot, F., Nevins, J. R., Mathey-Prevot, B., & You, L. (2014). Division of labour between Myc and G1 cyclins in cell cycle commitment and pace control. Nat Commun, 5, 4750. 10.1038/ncomms5750

Eapen, M. S., Hansbro, P. M., McAlinden, K., Kim, R. Y., Ward, C., Hackett, T. L., Walters, E. H., & Sohal, S. S. (2017). Abnormal M1/M2 macrophage phenotype profiles in the small airway wall and lumen in smokers and chronic obstructive pulmonary disease (COPD). Sci Rep, 7(1), 13392. 10.1038/s41598-017-13888-x

Eislmayr, K., Bestehorn, A., Morelli, L., Borroni, M., Vande Walle, L., Lamkanfi, M., & Kovarik, P. (2022). Nonredundancy of IL-1alpha and IL-1beta is defined by distinct regulation of tissues orchestrating resistance versus tolerance to infection. Sci Adv, 8(9), eabj7293. 10.1126/sciadv.abj7293

Enomoto, Y., Katsura, H., Fujimura, T., Ogata, A., Baba, S., Yamaoka, A., Kihara, M., Abe, T., Nishimura, O., Kadota, M., Hazama, D., Tanaka, Y., Maniwa, Y., Nagano, T., & Morimoto, M. (2023). Autocrine TGF-beta-positive feedback in profibrotic AT2-lineage cells plays a crucial role in non-inflammatory lung fibrogenesis. Nat Commun, 14(1), 4956. 10.1038/s41467-023-40617-y

Filippou, P. S., Karagiannis, G. S., & Constantinidou, A. (2020). Midkine (MDK) growth factor: a key player in cancer progression and a promising therapeutic target. Oncogene, 39(10), 2040–2054. 10.1038/s41388-019-1124-8

Franzen, O., Gan, L. M., & Bjorkegren, J. L. M. (2019). PanglaoDB: a web server for exploration of mouse and human single-cell RNA sequencing data. Database (Oxford), 2019. 10.1093/database/baz046

Frum, T., Hsu, P. P., Hein, R. F. C., Conchola, A. S., Zhang, C. J., Utter, O. R., Anand, A., Zhang, Y., Clark, S. G., Glass, I., Sexton, J. Z., & Spence, J. R. (2023). Opposing roles for TGFbeta- and BMP-signaling during nascent alveolar differentiation in the developing human lung. NPJ Regen Med, 8(1), 48. 10.1038/s41536-023-00325-z

Garcia-Arcos, I., Park, S. S., Mai, M., Alvarez-Buve, R., Chow, L., Cai, H., Baumlin-Schmid, N., Agudelo, C. W., Martinez, J., Kim, M. D., Dabo, A. J., Salathe, M., Goldberg, I. J., & Foronjy, R. F. (2022). LRP1 loss in airway epithelium exacerbates smoke-induced oxidative damage and airway remodeling. J Lipid Res, 63(4), 100185. 10.1016/j.jlr.2022.100185

Ge, Z., Chen, Y., Ma, L., Hu, F., & Xie, L. (2024). Macrophage polarization and its impact on idiopathic pulmonary fibrosis. Front Immunol, 15, 1444964. 10.3389/fimmu.2024.1444964

Ghonim, M. A., Boyd, D. F., Flerlage, T., & Thomas, P. G. (2023). Pulmonary inflammation and fibroblast immunoregulation: from bench to bedside. Journal of Clinical Investigation, 133(17). 10.1172/JCI170499

Ghosh, M. C., Gorantla, V., Makena, P. S., Luellen, C., Sinclair, S. E., Schwingshackl, A., & Waters, C. M. (2013). Insulin-like growth factor-I stimulates differentiation of ATII cells to ATI-like cells through activation of Wnt5a. Am J Physiol Lung Cell Mol Physiol, 305(3), L222–228. 10.1152/ajplung.00014.2013

Giri, J., Das, R., Nylen, E., Chinnadurai, R., & Galipeau, J. (2020). CCL2 and CXCL12 Derived from Mesenchymal Stromal Cells Cooperatively Polarize IL-10+ Tissue Macrophages to Mitigate Gut Injury. Cell Rep, 30(6), 1923–1934 e1924. 10.1016/j.celrep.2020.01.047

Goltsis, O., Bilodeau, C., Wang, J., Luo, D., Asgari, M., Bozec, L., Pettersson, A., Leibel, S. L., & Post, M. (2024). Influence of mesenchymal and biophysical components on distal lung organoid differentiation. Stem Cell Res Ther, 15(1), 273. 10.1186/s13287-024-03890-2

Goodwin, A. T., John, A. E., Joseph, C., Habgood, A., Tatler, A. L., Susztak, K., Palmer, M., Offermanns, S., Henderson, N. C., & Jenkins, R. G. (2023). Stretch regulates alveologenesis and homeostasis via mesenchymal Galphaq/11-mediated TGFbeta2 activation. Development, 150(9). 10.1242/dev.201046

Gray, Z., Shi, G., Wang, X., & Hu, Y. (2018). Macrophage inducible nitric oxide synthase promotes the initiation of lung squamous cell carcinoma by maintaining circulated inflammation. Cell Death Dis, 9(6), 642. 10.1038/s41419-018-0653-3

Greaney, A. M., Adams, T. S., Brickman Raredon, M. S., Gubbins, E., Schupp, J. C., Engler, A. J., Ghaedi, M., Yuan, Y., Kaminski, N., & Niklason, L. E. (2020). Platform Effects on Regeneration by Pulmonary Basal Cells as Evaluated by Single-Cell RNA Sequencing. Cell Rep, 30(12), 4250–4265 e4256. 10.1016/j.celrep.2020.03.004

Greaney, A. M., Sam, M., Obata, T., Wang, J., Adams, T. S., Schupp, J. C., Mizoguchi, S., Edelstein, S., Yuan, Y., Pavlina Baevova, Wang, N., Engler, A., Leiby, K., Tsuchiya, T., Homer, R., Kaminski, N., Langer, R., Niklason, L., & Ruslan Medzhitov. (2024). Engineered Whole Lungs for Tissue Biology. BioRxiv (Cold Spring Harbor Laboratory). 10.1101/2024.10.02.616240

Guan, T., Zhou, X., Zhou, W., & Lin, H. (2023). Regulatory T cell and macrophage crosstalk in acute lung injury: future perspectives. Cell Death Discov, 9(1), 9. 10.1038/s41420-023-01310-7

Hardie, W. D., Le Cras, T. D., Jiang, K., Tichelaar, J. W., Azhar, M., & Korfhagen, T. R. (2004). Conditional expression of transforming growth factor-alpha in adult mouse lung causes pulmonary fibrosis. Am J Physiol Lung Cell Mol Physiol, 286(4), L741–749. 10.1152/ajplung.00208.2003

Hoffmann, K., Obermayer, B., Honzke, K., Fatykhova, D., Demir, Z., Lowa, A., Alves, L. G. T., Wyler, E., Lopez-Rodriguez, E., Mieth, M., Baumgardt, M., Hoppe, J., Firsching, T. C., Tonnies, M., Bauer, T. T., Eggeling, S., Tran, H. L., Schneider, P., Neudecker, J.,…Kessler, M. (2022). Human alveolar progenitors generate dual lineage bronchioalveolar organoids. Commun Biol, 5(1), 875. 10.1038/s42003-022-03828-5

Holtzman, M. J., Morton, J. D., Shornick, L. P., Tyner, J. W., O’Sullivan, M. P., Antao, A., Lo, M., Castro, M., & Walter, M. J. (2002). Immunity, inflammation, and remodeling in the airway epithelial barrier: epithelial-viral-allergic paradigm. Physiol Rev, 82(1), 19–46. 10.1152/physrev.00020.2001

Hurley, K., Ding, J., Villacorta-Martin, C., Herriges, M. J., Jacob, A., Vedaie, M., Alysandratos, K. D., Sun, Y. L., Lin, C., Werder, R. B., Huang, J., Wilson, A. A., Mithal, A., Mostoslavsky, G., Oglesby, I., Caballero, I. S., Guttentag, S. H., Ahangari, F., Kaminski, N.,…Kotton, D. N. (2020). Reconstructed Single-Cell Fate Trajectories Define Lineage Plasticity Windows during Differentiation of Human PSC-Derived Distal Lung Progenitors. Cell Stem Cell, 26(4), 593–608 e598. 10.1016/j.stem.2019.12.009

Huttemann, M., Lee, I., Gao, X., Pecina, P., Pecinova, A., Liu, J., Aras, S., Sommer, N., Sanderson, T. H., Tost, M., Neff, F., Aguilar-Pimentel, J. A., Becker, L., Naton, B., Rathkolb, B., Rozman, J., Favor, J., Hans, W., Prehn, C.,…Grossman, L. I. (2012). Cytochrome c oxidase subunit 4 isoform 2-knockout mice show reduced enzyme activity, airway hyporeactivity, and lung pathology. FASEB J, 26(9), 3916–3930. 10.1096/fj.11-203273

Jeon, H. Y., Choi, J., Kraaier, L., Kim, Y. H., Eisenbarth, D., Yi, K., Kang, J. G., Kim, J. W., Shim, H. S., Lee, J. H., & Lim, D. S. (2022). Airway secretory cell fate conversion via YAP-mTORC1-dependent essential amino acid metabolism. EMBO J, 41(8), e109365. 10.15252/embj.2021109365

Johansson, K., Woodruff, P. G., & Ansel, K. M. (2021). Regulation of airway immunity by epithelial miRNAs. Immunology Reviews, 304(1), 141–153. 10.1111/imr.13028

Jones, D. L., Morley, M. P., Li, X., Ying, Y., Zhao, G., Schaefer, S. E., Rodriguez, L. R., Cardenas-Diaz, F. L., Li, S., Zhou, S., Chembazhi, U. V., Kim, M., Shen, C., Nottingham, A., Lin, S. M., Cantu, E., Diamond, J. M., Basil, M. C., Vaughan, A. E., & Morrisey, E. E. (2024). An injury-induced mesenchymal-epithelial cell niche coordinates regenerative responses in the lung. Science, 386(6727), eado5561. 10.1126/science.ado5561

Joshi, R., Heinz, A., Fan, Q., Guo, S., Monia, B., Schmelzer, C. E. H., Weiss, A. S., Batie, M., Parameshwaran, H., & Varisco, B. M. (2018). Role for Cela1 in Postnatal Lung Remodeling and Alpha-1 Antitrypsin-Deficient Emphysema. Am J Respir Cell Mol Biol, 59(2), 167–178. 10.1165/rcmb.2017-0361OC

Kadur Lakshminarasimha Murthy, P., Sontake, V., Tata, A., Kobayashi, Y., Macadlo, L., Okuda, K., Conchola, A. S., Nakano, S., Gregory, S., Miller, L. A., Spence, J. R., Engelhardt, J. F., Boucher, R. C., Rock, J. R., Randell, S. H., & Tata, P. R. (2022). Human distal lung maps and lineage hierarchies reveal a bipotent progenitor. Nature, 604(7904), 111–119. 10.1038/s41586-022-04541-3

Katsura, H., Kobayashi, Y., Tata, P. R., & Hogan, B. L. M. (2019). IL-1 and TNFalpha Contribute to the Inflammatory Niche to Enhance Alveolar Regeneration. Stem Cell Reports, 12(4), 657–666. 10.1016/j.stemcr.2019.02.013

Khedoe, P., van Schadewijk, W., Schwiening, M., Ng-Blichfeldt, J. P., Marciniak, S. J., Stolk, J., Gosens, R., & Hiemstra, P. S. (2023). Cigarette smoke restricts the ability of mesenchymal cells to support lung epithelial organoid formation. Front Cell Dev Biol, 11, 1165581. 10.3389/fcell.2023.1165581

Kim, C. F., Jackson, E. L., Woolfenden, A. E., Lawrence, S., Babar, I., Vogel, S., Crowley, D., Bronson, R. T., & Jacks, T. (2005). Identification of bronchioalveolar stem cells in normal lung and lung cancer. Cell, 121(6), 823–835. 10.1016/j.cell.2005.03.032

Konigshoff, M., & Eickelberg, O. (2010). WNT signaling in lung disease: a failure or a regeneration signal? Am J Respir Cell Mol Biol, 42(1), 21–31. 10.1165/rcmb.2008-0485TR

Konkimalla, A., Konishi, S., Macadlo, L., Kobayashi, Y., Farino, Z. J., Miyashita, N., El Haddad, L., Morowitz, J., Barkauskas, C. E., Agarwal, P., Souma, T., ElMallah, M. K., Tata, A., & Tata, P. R. (2023). Transitional cell states sculpt tissue topology during lung regeneration. Cell Stem Cell, 30(11), 1486–1502 e1489. 10.1016/j.stem.2023.10.001

Korfhagen, T. R., Swantz, R. J., Wert, S. E., McCarty, J. M., Kerlakian, C. B., Glasser, S. W., & Whitsett, J. A. (1994). Respiratory epithelial cell expression of human transforming growth factor-alpha induces lung fibrosis in transgenic mice. J Clin Invest, 93(4), 1691–1699. 10.1172/JCI117152

Leiby, K. L., Yuan, Y., Ng, R., Raredon, M. S. B., Adams, T. S., Baevova, P., Greaney, A. M., Hirschi, K. K., Campbell, S. G., Kaminski, N., Herzog, E. L., & Niklason, L. E. (2023). Rational engineering of lung alveolar epithelium. NPJ Regen Med, 8(1), 22. 10.1038/s41536-023-00295-2

Lewis, A. E., Kuwahara, A., Franzosi, J., & Bush, J. O. (2022). Tracheal separation is driven by NKX2-1-mediated repression of Efnb2 and regulation of endodermal cell sorting. Cell Rep, 38(11), 110510. 10.1016/j.celrep.2022.110510

Liang, J., Jung, Y., Tighe, R. M., Xie, T., Liu, N., Leonard, M., Gunn, M. D., Jiang, D., & Noble, P. W. (2012). A macrophage subpopulation recruited by CC chemokine ligand-2 clears apoptotic cells in noninfectious lung injury. Am J Physiol Lung Cell Mol Physiol, 302(9), L933–940. 10.1152/ajplung.00256.2011

Li, G., Ji, X. D., Gao, H., Zhao, J. S., Xu, J. F., Sun, Z. J., Deng, Y. Z., Shi, S., Feng, Y. X., Zhu, Y. Q., Wang, T., Li, J. J., & Xie, D. (2012). EphB3 suppresses non-small-cell lung cancer metastasis via a PP2A/RACK1/Akt signaling complex. Nat Commun, 3, 667. 10.1038/ncomms1675

Liberzon, A., Birger, C., Thorvaldsdottir, H., Ghandi, M., Mesirov, J. P., & Tamayo, P. (2015). The Molecular Signatures Database (MSigDB) hallmark gene set collection. Cell Syst, 1(6), 417–425. 10.1016/j.cels.2015.12.004

Lin, B., Shah, V. S., Chernoff, C., Sun, J., Shipkovenska, G. G., Vinarsky, V., Waghray, A., Xu, J., Leduc, A. D., Hintschich, C. A., Surve, M. V., Xu, Y., Capen, D. E., Villoria, J., Dou, Z., Hariri, L. P., & Rajagopal, J. (2024). Airway hillocks are injury-resistant reservoirs of unique plastic stem cells. Nature, 629(8013), 869–877. 10.1038/s41586-024-07377-1

Liu, C., Zheng, S., Lu, Z., Wang, Z., Wang, S., Feng, X., Wang, Y., Sun, N., & He, J. (2022). S100A7 attenuates immunotherapy by enhancing immunosuppressive tumor microenvironment in lung squamous cell carcinoma. Signal Transduct Target Ther, 7(1), 368. 10.1038/s41392-022-01196-4

Liu, F., Lagares, D., Choi, K. M., Stopfer, L., Marinkovic, A., Vrbanac, V., Probst, C. K., Hiemer, S. E., Sisson, T. H., Horowitz, J. C., Rosas, I. O., Fredenburgh, L. E., Feghali-Bostwick, C., Varelas, X., Tager, A. M., & Tschumperlin, D. J. (2015). Mechanosignaling through YAP and TAZ drives fibroblast activation and fibrosis. Am J Physiol Lung Cell Mol Physiol, 308(4), L344–357. 10.1152/ajplung.00300.2014

Liu, G., Wang, Y., Yang, L., Zou, B., Gao, S., Song, Z., & Lin, Z. (2019). Tetraspanin 1 as a mediator of fibrosis inhibits EMT process and Smad2/3 and beta-catenin pathway in human pulmonary fibrosis. J Cell Mol Med, 23(5), 3583–3596. 10.1111/jcmm.14258

Liu, M. Y., Chen, B., Borji, M., Garcia de Alba Rivas, C., Dost, A. F. M., Moye, A. L., Movval Abdulla, N., Paschini, M., Rollins, S. D., Wang, R., Schnapp, L. M., Khalil, H. A., Wu, C. J., Sharma, N. S., & Kim, C. F. (2024). Human Airway and Alveolar Organoids from BAL Fluid. Am J Respir Crit Care Med, 209(12), 1501–1504. 10.1164/rccm.202310-1831LE

Liu, T., Laidlaw, T. M., Feng, C., Xing, W., Shen, S., Milne, G. L., & Boyce, J. A. (2012). Prostaglandin E2 deficiency uncovers a dominant role for thromboxane A2 in house dust mite-induced allergic pulmonary inflammation. Proc Natl Acad Sci U S A, 109(31), 12692–12697. 10.1073/pnas.1207816109

Manneken, J. D., & Currie, P. D. (2023). Macrophage-stem cell crosstalk: regulation of the stem cell niche. Development, 150(8). 10.1242/dev.201510

Martins, L. R., Sieverling, L., Michelhans, M., Schiller, C., Erkut, C., Grunewald, T. G. P., Triana, S., Frohling, S., Velten, L., Glimm, H., & Scholl, C. (2024). Single-cell division tracing and transcriptomics reveal cell types and differentiation paths in the regenerating lung. Nat Commun, 15(1), 2246. 10.1038/s41467-024-46469-4

Matkovic Leko, I., Schneider, R. T., Thimraj, T. A., Schrode, N., Beitler, D., Liu, H. Y., Beaumont, K., Chen, Y. W., & Snoeck, H. W. (2023). A distal lung organoid model to study interstitial lung disease, viral infection and human lung development. Nat Protoc, 18(7), 2283–2312. 10.1038/s41596-023-00827-6

Matsuda, K., Hirayama, D., Hino, N., Kuno, S., Sakaue-Sawano, A., Miyawaki, A., Matsuda, M., & Terai, K. (2023). Knockout of all ErbB-family genes delineates their roles in proliferation, survival and migration. J Cell Sci, 136(16). 10.1242/jcs.261199

Medzhitov, R. (2021). The spectrum of inflammatory responses. Science, 374(6571), 1070–1075. 10.1126/science.abi5200

Meli, V. S., Atcha, H., Veerasubramanian, P. K., Nagalla, R. R., Luu, T. U., Chen, E. Y., Guerrero-Juarez, C. F., Yamaga, K., Pandori, W., Hsieh, J. Y., Downing, T. L., Fruman, D. A., Lodoen, M. B., Plikus, M. V., Wang, W., & Liu, W. F. (2020). YAP-mediated mechanotransduction tunes the macrophage inflammatory response. Sci Adv, 6(49). 10.1126/sciadv.abb8471

Minagawa, S., Lou, J., Seed, R. I., Cormier, A., Wu, S., Cheng, Y., Murray, L., Tsui, P., Connor, J., Herbst, R., Govaerts, C., Barker, T., Cambier, S., Yanagisawa, H., Goodsell, A., Hashimoto, M., Brand, O. J., Cheng, R., Ma, R.,…Nishimura, S. L. (2014). Selective targeting of TGF-beta activation to treat fibroinflammatory airway disease. Sci Transl Med, 6(241), 241ra279. 10.1126/scitranslmed.3008074

Micochova, P., Zhao, N., Shabana, O., Fischer, R., & Gupta, R. K. (2024). CD4 T cell contact drives macrophage cell cycle G0-G1 transition. Signal Transduct Target Ther, 9(1), 348. 10.1038/s41392-024-02053-2

Mo, Y., Kang, H., Bang, J. Y., Shin, J. W., Kim, H. Y., Cho, S. H., & Kang, H. R. (2022). Intratracheal administration of mesenchymal stem cells modulates lung macrophage polarization and exerts anti-asthmatic effects. Scientific Reports, 12(1), 11728. 10.1038/s41598-022-14846-y

Montoro, D. T., Haber, A. L., Biton, M., Vinarsky, V., Lin, B., Birket, S. E., Yuan, F., Chen, S., Leung, H. M., Villoria, J., Rogel, N., Burgin, G., Tsankov, A. M., Waghray, A., Slyper, M., Waldman, J., Nguyen, L., Dionne, D., Rozenblatt-Rosen, O.,…Rajagopal, J. (2018). A revised airway epithelial hierarchy includes CFTR-expressing ionocytes. Nature, 560(7718), 319–324. 10.1038/s41586-018-0393-7

Morrison, T. J., Jackson, M. V., Cunningham, E. K., Kissenpfennig, A., McAuley, D. F., O’Kane, C. M., & Krasnodembskaya, A. D. (2017). Mesenchymal Stromal Cells Modulate Macrophages in Clinically Relevant Lung Injury Models by Extracellular Vesicle Mitochondrial Transfer. Am J Respir Crit Care Med, 196(10), 1275–1286. 10.1164/rccm.201701-0170OC

Morrow, J. D., Yun, J. H., & Hersh, C. P. (2024). Lung Single-Cell Transcriptomics in Alpha-1 Antitrypsin Deficiency. Am J Respir Cell Mol Biol, 71(2), 254–256. 10.1165/rcmb.2024-0064LE

Murrow, L. M., Weber, R. J., & Gartner, Z. J. (2017). Dissecting the stem cell niche with organoid models: an engineering-based approach. Development, 144(6), 998–1007. 10.1242/dev.140905

Nalio Ramos, R., Missolo-Koussou, Y., Gerber-Ferder, Y., Bromley, C. P., Bugatti, M., Nunez, N. G., Tosello Boari, J., Richer, W., Menger, L., Denizeau, J., Sedlik, C., Caudana, P., Kotsias, F., Niborski, L. L., Viel, S., Bohec, M., Lameiras, S., Baulande, S., Lesage, L.,…Helft, J. (2022). Tissue-resident FOLR2(+) macrophages associate with CD8(+) T cell infiltration in human breast cancer. Cell, 185(7), 1189–1207 e1125. 10.1016/j.cell.2022.02.021

Negretti, N. M., Son, Y., Crooke, P., Plosa, E. J., Benjamin, J. T., Jetter, C. S., Bunn, C., Mignemi, N., Marini, J., Hackett, A. N., Ransom, M., Garg, S., Nichols, D., Guttentag, S. H., Pua, H. H., Blackwell, T. S., Zacharias, W., Frank, D. B., Kozub, J. A.,…Sucre, J. M. (2025). Epithelial outgrowth through mesenchymal rings drives lung alveologenesis. JCI Insight, 10(4). 10.1172/jci.insight.187876

Neri, T., Armani, C., Pegoli, A., Cordazzo, C., Carmazzi, Y., Brunelleschi, S., Bardelli, C., Breschi, M. C., Paggiaro, P., & Celi, A. (2011). Role of NF-kappaB and PPAR-gamma in lung inflammation induced by monocyte-derived microparticles. Eur Respir J, 37(6), 1494–1502. 10.1183/09031936.00023310

Nichane, M., Javed, A., Sivakamasundari, V., Ganesan, M., Ang, L. T., Kraus, P., Lufkin, T., Loh, K. M., & Lim, B. (2017). Isolation and 3D expansion of multipotent Sox9(+) mouse lung progenitors. Nature Methods, 14(12), 1205–1212. 10.1038/nmeth.4498

Nikolic, M. Z., & Rawlins, E. L. (2017). Lung Organoids and Their Use To Study Cell-Cell Interaction. Curr Pathobiol Rep, 5(2), 223–231. 10.1007/s40139-017-0137-7

Nolte, R. T., Wisely, G. B., Westin, S., Cobb, J. E., Lambert, M. H., Kurokawa, R., Rosenfeld, M. G., Willson, T. M., Glass, C. K., & Milburn, M. V. (1998). Ligand binding and co-activator assembly of the peroxisome proliferator-activated receptor-gamma. Nature, 395(6698), 137–143. 10.1038/25931

Nwokoye, P. N., & Abilez, O. J. (2024). Bioengineering methods for vascularizing organoids. Cell Rep Methods, 4(6), 100779. 10.1016/j.crmeth.2024.100779

Obata, T., Mizoguchi, S., Greaney, A. M., Adams, T., Yuan, Y., Edelstein, S., Leiby, K. L., Rivero, R., Wang, N., Kim, H., Yang, J., Schupp, J. C., Stitelman, D., Tsuchiya, T., Levchenko, A., Kaminski, N., Niklason, L. E., & Brickman Raredon, M. S. (2024). Organ Boundary Circuits Regulate Sox9+ Alveolar Tuft Cells During Post-Pneumonectomy Lung Regeneration. bioRxiv. 10.1101/2024.01.07.574469

O’Reilly, M., & Thebaud, B. (2014). Animal models of bronchopulmonary dysplasia. The term rat models. Am J Physiol Lung Cell Mol Physiol, 307(12), L948–958. 10.1152/ajplung.00160.2014

Ostrin, E. J., Little, D. R., Gerner-Mauro, K. N., Sumner, E. A., Rios-Corzo, R., Ambrosio, E., Holt, S. E., Forcioli-Conti, N., Akiyama, H., Hanash, S. M., Kimura, S., Huang, S. X. L., & Chen, J. (2018). beta-Catenin maintains lung epithelial progenitors after lung specification. Development, 145(5). 10.1242/dev.160788

Otto, Y., Shipkovenska, G. G., Liu, Q., Sun, J., Hariri, L. P., & Rajagopal, J. (2025). The identification and characterization of hillocks in the postnatal mouse and human airway. biorxiv (Cold Spring Laboratory). 10.1101/2025.04.15.648833

Patel, D. F., Peiro, T., Shoemark, A., Akthar, S., Walker, S. A., Grabiec, A. M., Jackson, P. L., Hussell, T., Gaggar, A., Xu, X., Trevor, J. L., Li, J., Steele, C., Tavernier, G., Blalock, J. E., Niven, R. M., Gregory, L. G., Simpson, A., Lloyd, C. M., & Snelgrove, R. J. (2018). An extracellular matrix fragment drives epithelial remodeling and airway hyperresponsiveness. Sci Transl Med, 10(455). 10.1126/scitranslmed.aaq0693

Radomir, L., Kramer, M. P., Perpinial, M., Schottlender, N., Rabani, S., David, K., Wiener, A., Lewinsky, H., Becker-Herman, S., Aharoni, R., Milo, R., Mauri, C., & Shachar, I. (2021). The survival and function of IL-10-producing regulatory B cells are negatively controlled by SLAMF5. Nature Communications, 12(1), 1893. 10.1038/s41467-021-22230-z

Raja, M. R. K., Gupta, G., Atkinson, G., Kathrein, K., Armstrong, A., Gower, M., Roninson, I., Broude, E., Chen, M., Ji, H., Lim, C., Wang, H., Fan, D., Xu, P., Li, J., Zhou, G., & Chen, H. (2024). Host-derived Interleukin 1alpha induces an immunosuppressive tumor microenvironment via regulating monocyte-to-macrophage differentiation. bioRxiv. 10.1101/2024.05.03.592354

Ramilowski, J. A., Goldberg, T., Harshbarger, J., Kloppmann, E., Lizio, M., Satagopam, V. P., Itoh, M., Kawaji, H., Carninci, P., Rost, B., & Forrest, A. R. (2015). A draft network of ligand-receptor-mediated multicellular signalling in human. Nat Commun, 6, 7866. 10.1038/ncomms8866

Raredon, M. S. B., Yang, J., Kothapalli, N., Lewis, W., Kaminski, N., Niklason, L. E., & Kluger, Y. (2023). Comprehensive visualization of cell-cell interactions in single-cell and spatial transcriptomics with NICHES. Bioinformatics, 39(1). 10.1093/bioinformatics/btac775

Rockich, B. E., Hrycaj, S. M., Shih, H. P., Nagy, M. S., Ferguson, M. A., Kopp, J. L., Sander, M., Wellik, D. M., & Spence, J. R. (2013). Sox9 plays multiple roles in the lung epithelium during branching morphogenesis. Proc Natl Acad Sci U S A, 110(47), E4456–4464. 10.1073/pnas.1311847110

Russo, R. C., Guabiraba, R., Garcia, C. C., Barcelos, L. S., Roffe, E., Souza, A. L., Amaral, F. A., Cisalpino, D., Cassali, G. D., Doni, A., Bertini, R., & Teixeira, M. M. (2009). Role of the chemokine receptor CXCR2 in bleomycin-induced pulmonary inflammation and fibrosis. Am J Respir Cell Mol Biol, 40(4), 410–421. 10.1165/rcmb.2007-0364OC

Sachs, N., Papaspyropoulos, A., Zomer-van Ommen, D. D., Heo, I., Bottinger, L., Klay, D., Weeber, F., Huelsz-Prince, G., Iakobachvili, N., Amatngalim, G. D., de Ligt, J., van Hoeck, A., Proost, N., Viveen, M. C., Lyubimova, A., Teeven, L., Derakhshan, S., Korving, J., Begthel, H.,…Clevers, H. (2019). Long-term expanding human airway organoids for disease modeling. EMBO J, 38(4). 10.15252/embj.2018100300

Salwig, I., Spitznagel, B., Vazquez-Armendariz, A. I., Khalooghi, K., Guenther, S., Herold, S., Szibor, M., & Braun, T. (2019). Bronchioalveolar stem cells are a main source for regeneration of distal lung epithelia in vivo. EMBO J, 38(12). 10.15252/embj.2019102099

Sanchez-Esteban, J., Cicchiello, L. A., Wang, Y., Tsai, S. W., Williams, L. K., Torday, J. S., & Rubin, L. P. (2001). Mechanical stretch promotes alveolar epithelial type II cell differentiation. J Appl Physiol (1985), *91*(2), 589-595. 10.1152/jappl.2001.91.2.589

Sapudom, J., Tipay, P., & Teo, J. C. (2024). Mechano-mediated M2 macrophage polarization and immune suppression in stiffened tumor microenvironment. biorxiv (Cold Spring Laboratory). 10.1101/2024.07.29.605566

Seo, H. R., Jeong, H. E., Joo, H. J., Choi, S. C., Park, C. Y., Kim, J. H., Choi, J. H., Cui, L. H., Hong, S. J., Chung, S., & Lim, D. S. (2016). Intrinsic FGF2 and FGF5 promotes angiogenesis of human aortic endothelial cells in 3D microfluidic angiogenesis system. Scientific Reports, 6, 28832. 10.1038/srep28832

Shao, X., Taha, I. N., Clauser, K. R., Gao, Y. T., & Naba, A. (2020). MatrisomeDB: the ECM-protein knowledge database. Nucleic Acids Res, 48(D1), D1136–D1144. 10.1093/nar/gkz849

Shen, W. K., Chen, S. Y., Gan, Z. Q., Zhang, Y. Z., Yue, T., Chen, M. M., Xue, Y., Hu, H., & Guo, A. Y. (2023). AnimalTFDB 4.0: a comprehensive animal transcription factor database updated with variation and expression annotations. Nucleic Acids Res, 51(D1), D39–D45. 10.1093/nar/gkac907

Shiraishi, K., Shah, P. P., Morley, M. P., Loebel, C., Santini, G. T., Katzen, J., Basil, M. C., Lin, S. M., Planer, J. D., Cantu, E., Jones, D. L., Nottingham, A. N., Li, S., Cardenas-Diaz, F. L., Zhou, S., Burdick, J. A., Jain, R., & Morrisey, E. E. (2023). Biophysical forces mediated by respiration maintain lung alveolar epithelial cell fate. Cell, 186(7), 1478–1492 e1415. 10.1016/j.cell.2023.02.010

Sintes, J., Romero, X., de Salort, J., Terhorst, C., & Engel, P. (2010). Mouse CD84 is a pan-leukocyte cell-surface molecule that modulates LPS-induced cytokine secretion by macrophages. J Leukoc Biol, 88(4), 687–697. 10.1189/jlb.1109756

Stahl, S., Kaminskyy, V. O., Efazat, G., Hyrslova Vaculova, A., Rodriguez-Nieto, S., Moshfegh, A., Lewensohn, R., Viktorsson, K., & Zhivotovsky, B. (2013). Inhibition of Ephrin B3-mediated survival signaling contributes to increased cell death response of non-small cell lung carcinoma cells after combined treatment with ionizing radiation and PKC 412. Cell Death Dis, 4(1), e454. 10.1038/cddis.2012.188

Stouch, A. N., McCoy, A. M., Greer, R. M., Lakhdari, O., Yull, F. E., Blackwell, T. S., Hoffman, H. M., & Prince, L. S. (2016). IL-1beta and Inflammasome Activity Link Inflammation to Abnormal Fetal Airway Development. J Immunol, 196(8), 3411–3420. 10.4049/jimmunol.1500906

Street, K., Risso, D., Fletcher, R. B., Das, D., Ngai, J., Yosef, N., Purdom, E., & Dudoit, S. (2018). Slingshot: cell lineage and pseudotime inference for single-cell transcriptomics. BMC Genomics, 19(1), 477. 10.1186/s12864-018-4772-0

Strunz, M., Simon, L. M., Ansari, M., Kathiriya, J. J., Angelidis, I., Mayr, C. H., Tsidiridis, G., Lange, M., Mattner, L. F., Yee, M., Ogar, P., Sengupta, A., Kukhtevich, I., Schneider, R., Zhao, Z., Voss, C., Stoeger, T., Neumann, J. H. L., Hilgendorff, A.,…Schiller, H. B. (2020). Alveolar regeneration through a Krt8+ transitional stem cell state that persists in human lung fibrosis. Nat Commun, 11(1), 3559. 10.1038/s41467-020-17358-3

Suezawa, T., Kanagaki, S., Moriguchi, K., Masui, A., Nakao, K., Toyomoto, M., Tamai, K., Mikawa, R., Hirai, T., Murakami, K., Hagiwara, M., & Gotoh, S. (2021). Disease modeling of pulmonary fibrosis using human pluripotent stem cell-derived alveolar organoids. Stem Cell Reports, 16(12), 2973–2987. 10.1016/j.stemcr.2021.10.015

Sun, D., Llora Batlle, O., van den Ameele, J., Thomas, J. C., He, P., Lim, K., Tang, W., Xu, C., Meyer, K. B., Teichmann, S. A., Marioni, J. C., Jackson, S. P., Brand, A. H., & Rawlins, E. L. (2022). SOX9 maintains human foetal lung tip progenitor state by enhancing WNT and RTK signalling. EMBO J, 41(21), e111338. 10.15252/embj.2022111338

Suwara, M. I., Green, N. J., Borthwick, L. A., Mann, J., Mayer-Barber, K. D., Barron, L., Corris, P. A., Farrow, S. N., Wynn, T. A., Fisher, A. J., & Mann, D. A. (2014). IL-1alpha released from damaged epithelial cells is sufficient and essential to trigger inflammatory responses in human lung fibroblasts. Mucosal Immunology, 7(3), 684–693. 10.1038/mi.2013.87

Tata, P. R., & Rajagopal, J. (2017). Plasticity in the lung: making and breaking cell identity. Development, 144(5), 755–766. 10.1242/dev.143784

Thomas, K., Rossaint, J., Ludwig, N., Mersmann, S., Kotting, N., Grenzheuser, J., Schemmelmann, L., Oguama, M., Margraf, A., Block, H., Henke, K., Hellenthal, K., Mirakaj, V., Gerke, V., Hansen, U., Gaher, K., Engelhardt, M., Roth, J., Eble, J.,…Zarbock, A. (2025). Alveolar epithelial and vascular CXCR2 mediates transcytosis of CXCL1 in inflamed lungs. Nat Commun, 16(1), 4846. 10.1038/s41467-025-60174-w

Toth, A., Kannan, P., Snowball, J., Kofron, M., Wayman, J. A., Bridges, J. P., Miraldi, E. R., Swarr, D., & Zacharias, W. J. (2023). Alveolar epithelial progenitor cells require Nkx2-1 to maintain progenitor-specific epigenomic state during lung homeostasis and regeneration. Nat Commun, 14(1), 8452. 10.1038/s41467-023-44184-0

Van den Berge, K., Roux de Bezieux, H., Street, K., Saelens, W., Cannoodt, R., Saeys, Y., Dudoit, S., & Clement, L. (2020). Trajectory-based differential expression analysis for single-cell sequencing data. Nat Commun, 11(1), 1201. 10.1038/s41467-020-14766-3

Vining, K. H., & Mooney, D. J. (2017). Mechanical forces direct stem cell behaviour in development and regeneration. Nat Rev Mol Cell Biol, 18(12), 728–742. 10.1038/nrm.2017.108

Wang, Y., Dede, M., Mohanty, V., Dou, J., Li, Z., & Chen, K. (2024). A statistical approach for systematic identification of transition cells from scRNA-seq data. Cell Rep Methods, 4(12), 100913. 10.1016/j.crmeth.2024.100913

Wang, C., Yu, Q., Song, T., Wang, Z., Song, L., Yang, Y., Shao, J., Li, J., Ni, Y., Chao, N., Zhang, L., & Li, W. (2022). The heterogeneous immune landscape between lung adenocarcinoma and squamous carcinoma revealed by single-cell RNA sequencing. Signal Transduct Target Ther, 7(1), 289. 10.1038/s41392-022-01130-8

Wang, H., Hosakote, Y. M., Boor, P. J., Yang, J., Zhang, Y., Yu, X., Gonzales, C., Levine, C. B., McLellan, S., Cloutier, N., Xie, X., Shi, P. Y., Ren, P., Hu, H., Sun, K., Soong, L., Sun, J., & Liang, Y. (2024). The alarmin IL-33 exacerbates pulmonary inflammation and immune dysfunction in SARS-CoV-2 infection. iScience, 27(6), 110117. 10.1016/j.isci.2024.110117

Warburton, D., Schwarz, M., Tefft, D., Flores-Delgado, G., Anderson, K. D., & Cardoso, W. V. (2000). The molecular basis of lung morphogenesis. Mechanisms of Development, 92(1), 55–81. 10.1016/s0925-4773(99)00325-1

Warren, R., Klinkhammer, K., Lyu, H., Knopp, J., Yuan, T., Yao, C., Stripp, B., & De Langhe, S. P. (2024). Cell competition drives bronchiolization and pulmonary fibrosis. Nat Commun, 15(1), 10624. 10.1038/s41467-024-54997-2

Wickham, H. (2012). A Layer Grammar of Graphics. Journal of Computational and Graphical Statistics, 19(1), 3–28. 10.1198/jcgs.2009.07098

Willart, M. A., Deswarte, K., Pouliot, P., Braun, H., Beyaert, R., Lambrecht, B. N., & Hammad, H. (2012). Interleukin-1alpha controls allergic sensitization to inhaled house dust mite via the epithelial release of GM-CSF and IL-33. J Exp Med, 209(8), 1505–1517. 10.1084/jem.20112691

Worsdorfer, P., Dalda, N., Kern, A., Kruger, S., Wagner, N., Kwok, C. K., Henke, E., & Ergun, S. (2019). Generation of complex human organoid models including vascular networks by incorporation of mesodermal progenitor cells. Scientific Reports, 9(1), 15663. 10.1038/s41598-019-52204-7

Wu, T., Hu, E., Xu, S., Chen, M., Guo, P., Dai, Z., Feng, T., Zhou, L., Tang, W., Zhan, L., Fu, X., Liu, S., Bo, X., & Yu, G. (2021). clusterProfiler 4.0: A universal enrichment tool for interpreting omics data. Innovation (Camb), 2(3), 100141. 10.1016/j.xinn.2021.100141

Xu, B., Chen, C., Chen, H., Zheng, S. G., Bringas, P., Jr., Xu, M., Zhou, X., Chen, D., Umans, L., Zwijsen, A., & Shi, W. (2011). Smad1 and its target gene Wif1 coordinate BMP and Wnt signaling activities to regulate fetal lung development. Development, 138(5), 925–935. 10.1242/dev.062687

Zepp, J. A., Zacharias, W. J., Frank, D. B., Cavanaugh, C. A., Zhou, S., Morley, M. P., & Morrisey, E. E. (2017). Distinct Mesenchymal Lineages and Niches Promote Epithelial Self-Renewal and Myofibrogenesis in the Lung. Cell, 170(6), 1134–1148 e1110. 10.1016/j.cell.2017.07.034

Zhang, H., Tsui, C. K., Garcia, G., Joe, L. K., Wu, H., Maruichi, A., Fan, W., Pandovski, S., Yoon, P. H., Webster, B. M., Durieux, J., Frankino, P. A., Higuchi-Sanabria, R., & Dillin, A. (2024). The extracellular matrix integrates mitochondrial homeostasis. Cell, 187(16), 4289–4304 e4226. 10.1016/j.cell.2024.05.057

Zhang, S., Zhang, L., Wang, L., Wang, H., Wu, J., Cai, H., Mo, C., & Yang, J. (2023). Machine learning identified MDK score has prognostic value for idiopathic pulmonary fibrosis based on integrated bulk and single cell expression data. Front Genet, 14, 1246983. 10.3389/fgene.2023.1246983

Zhang, Y., Black, K. E., Phung, T. N., Thundivalappil, S. R., Lin, T., Wang, W., Xu, J., Zhang, C., Hariri, L. P., Lapey, A., Li, H., Lerou, P. H., Ai, X., Que, J., Park, J. A., Hurley, B. P., & Mou, H. (2023). Human Airway Basal Cells Undergo Reversible Squamous Differentiation and Reshape Innate Immunity. Am J Respir Cell Mol Biol, 68(6), 664–678. 10.1165/rcmb.2022-0299OC

Zhang, Y., Dube, P. E., Washington, M. K., Yan, F., & Polk, D. B. (2012). ErbB2 and ErbB3 regulate recovery from dextran sulfate sodium-induced colitis by promoting mouse colon epithelial cell survival. Lab Invest, 92(3), 437–450. 10.1038/labinvest.2011.192

Zhao, L., Yee, M., & O’Reilly, M. A. (2013). Transdifferentiation of alveolar epithelial type II to type I cells is controlled by opposing TGF-beta and BMP signaling. Am J Physiol Lung Cell Mol Physiol, 305(6), L409–418. 10.1152/ajplung.00032.2013

Zheng, L., Zhou, Z., Lin, L., Alber, S., Watkins, S., Kaminski, N., Choi, A. M., & Morse, D. (2009). Carbon monoxide modulates alpha-smooth muscle actin and small proline rich-1a expression in fibrosis. Am J Respir Cell Mol Biol, 41(1), 85–92. 10.1165/rcmb.2007-0401OC

Zhou, C., Gao, Y., Ding, P., Wu, T., & Ji, G. (2023). The role of CXCL family members in different diseases. Cell Death Discov, 9(1), 212. 10.1038/s41420-023-01524-9

Zimmermann, N., Doepker, M. P., Witte, D. P., Stringer, K. F., Fulkerson, P. C., Pope, S. M., Brandt, E. B., Mishra, A., King, N. E., Nikolaidis, N. M., Wills-Karp, M., Finkelman, F. D., & Rothenberg, M. E. (2005). Expression and regulation of small proline-rich protein 2 in allergic inflammation. Am J Respir Cell Mol Biol, 32(5), 428–435. 10.1165/rcmb.2004-0269OC

